# A Stochastic Mechanism Drives Fast Substrate Translocation in the AAA+ Machine ClpB

**DOI:** 10.1101/2025.03.13.643017

**Authors:** Remi Casier, Dorit Levy, Inbal Riven, Yoav Barak, Gilad Haran

## Abstract

How biological machines harness ATP to drive mechanical work remains a crucial question. Structural studies of protein-translocating AAA+ machines proposed a coupled and sequential translocation process, whereby ATP hydrolysis events lead to short threading steps. Yet, direct real-time observation of these events remains elusive. Here, we employ single-molecule FRET spectroscopy to track substrate translocation through ClpB, a quality control AAA+ machine. We isolate ClpB and its substrate within lipid vesicles and find that translocation events, while dependent on ATP, take milliseconds, much faster than ATP hydrolysis times. Surprisingly, the translocation rate depends weakly on temperature and ATP concentration. Using three-color FRET experiments, we find that translocation events can occur bidirectionally but are not always complete. Replacing ATP with the slowly hydrolysable analog ATPγS abolishes both rapid translocation and directionality. These results indicate a fast, stochastic Brownian-motor-like mechanism, redefining how ATP is coupled with mechanical action in AAA+ machines.

## Introduction

The ubiquitous superfamily of AAA+ (ATPases associated with diverse cellular activity) proteins participates in essential cellular activities, including protein disaggregation and degradation, DNA replication and recombination, organelle biogenesis, and cellular remodeling (*1-3*). Despite their remarkable functional diversity, they typically assemble into ring-like hexamers and share conserved nucleotide-binding domains (NBDs) that couple ATP turnover to mechanical work (*3*). However, how ATP energy is utilized to promote the activity of these proteins remains an active area of study. In recent years, cryogenic electron microscopy has yielded numerous 3D structures of AAA+ machines, with a particular focus on those that unfold or disaggregate client proteins and then translocate them through their central lumen for protein quality control activities. Rather than adopting a fully symmetric ring, many of the AAA+ assemblies were found to form a spiral-like configuration (*6-9*). These observations, based on static structural information, have led to the proposed hand-over-hand model for substrate translocation. In this model, the top subunit sequentially moves downwards following ATP hydrolysis, pulling the substrate protein in a *power-stroke-like* manner (*6, 10-15*); each ATP hydrolysis event directly generates a single forceful translocation step of two amino-acid (*aa*) residues (*10*). However, complementary biochemical and biophysical studies point towards a more stochastic and rapid mechanism. For instance, experiments on the AAA+ machines ClpA and ClpX demonstrated that they can function in translocating proteins even when up to three of their subunits are defective in ATP hydrolysis (*16, 17*), suggesting that full coordination is not strictly required. Further, optical tweezer experiments on the AAA+ disaggregation machine ClpB (*18, 19*) found that it could translocate more than 500 *aa’*s in a second (*20*), outpacing the hydrolysis rate by more than two orders of magnitude. Additional studies have also pointed to the viable possibility of partial substrate threading by ClpB (*21*).

These observations underscore the need for direct, real-time monitoring of substrate translocation to obtain decisive evidence for the mechanism of action of these machines. Here, we address this challenge by using single-molecule Förster resonance energy transfer (smFRET) experiments to directly track, in real-time, the motion of a substrate protein through the lumen of ClpB. As a model substrate, we use κ-casein, which has a high degree of intrinsic disorder (*22, 23*). This property allows us to bypass the unfolding requirement of structured proteins, which has been studied elsewhere (see e.g. (*24-26*)) and thereby isolate the translocation process itself, which is expected to be faster and is a central component of ClpB-mediated disaggregation and AAA+ machine interaction with proteins in general. Studying this step in isolation facilitates probing the fundamental mechanism underlying substrate movement through the central pore. Our experiments are performed on ClpB molecules isolated with casein substrates within surface-immobilized lipid vesicles. They lead us to propose a stochastic, ATP-dependent translocation mechanism in which the consumption of ATP regulates the directionality of substrate diffusion within the lumen. This *Brownian-motor-like* mechanism (*14*) enables efficient translocation, achieving complete threading of an unfolded substrate with the consumption of only one or two ATP molecules.

## Results

### Observing substrate translocation events by ClpB

To gain insight into the molecular mechanism of ClpB (*Thermus thermophilus*), we utilized real-time smFRET to monitor the translocation of the model substrate κ-casein, an unstructured protein (*22, 23*). We inserted individual copies of the fluorescently tagged proteins inside surface-tethered lipid vesicles (Fig. 1A), which afforded several distinct advantages (*27*). The confinement into ca. 120 nm diameter vesicles (Fig. S1) created a high local protein concentration (ca. 2 – 3 μM) which facilitated the bimolecular interactions between the weakly associating species (*K*_D,app_ (ATP) = 0.6 ± 0.1 μM, Fig. S2), while still allowing the proteins to diffuse freely (Fig. S3). The immobilized vesicles also provided extended observation times for individual ClpB molecules, allowing us to monitor repeated translocation events. The liposomes were prepared from 1,2-dimyristoyl-sn-glycero-3-phosphocholine (DMPC), a lipid that is near its gel-to-liquid crystalline transition at room temperature (22.5 °C), allowing for the permeation of ions such as ATP (*28, 29*). The rapid and reversible permeability of the membrane (see *liposome permeability* and Fig. S4 in the SI) facilitated the maintenance of constant ATP levels.

**Figure 1.**
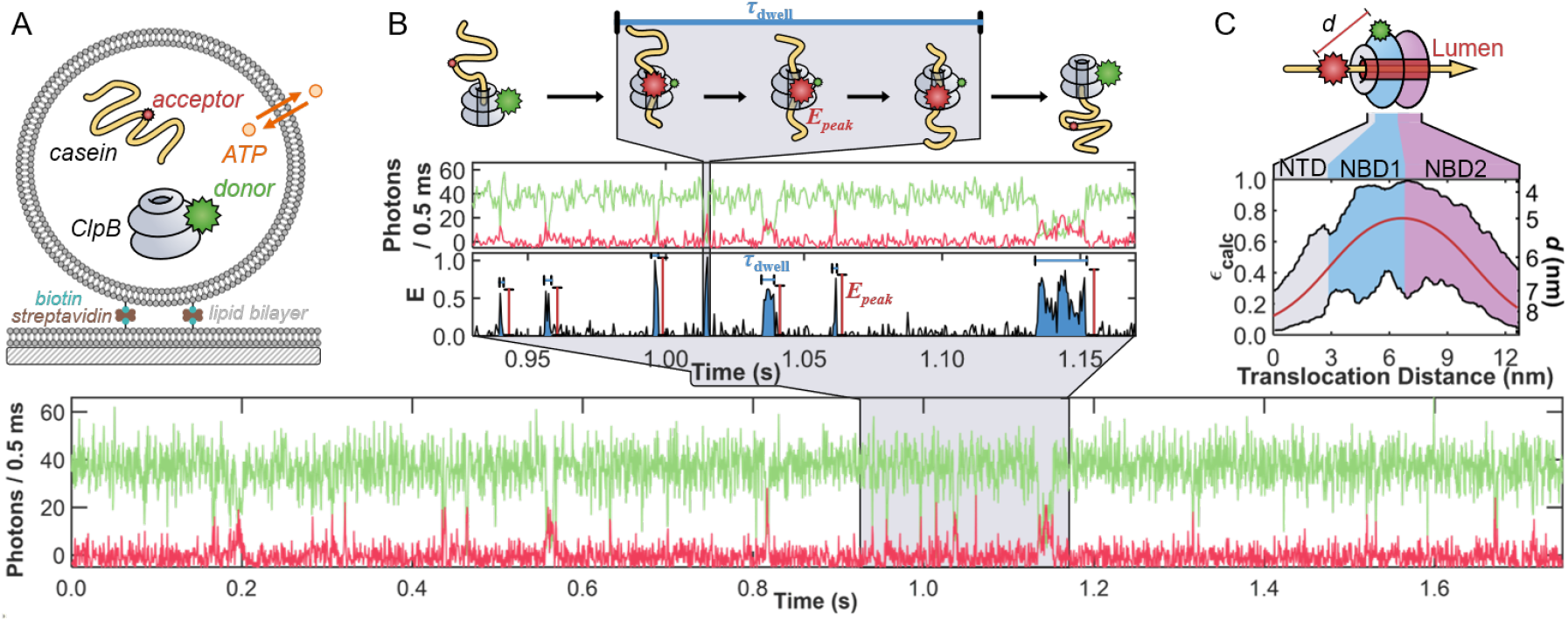
Observation of multiple fast events of translocation through the lumen of ClpB facilitated by porous vesicles. (**A**) Schematic overview of a ClpB and substrate-loaded liposome (diameter ca. 120 nm) immobilized on a glass-supported lipid bilayer through a biotin-streptavidin linkage. Only liposomes containing individual copies of a fluorescently labeled ClpB hexamer (grey, with a green donor, Cy3b, on residue 359) and casein (yellow, with a red acceptor, CF660R) were included in the analysis. Liposomes remained permeable to ATP (orange) at 22.5 °C, ensuring replenishment of the ATP supply consumed by ClpB. (**B**) A sample fluorescence trajectory (bottom) of a ClpB-casein pair inside a liposome in the presence of 2 mM ATP (donor signal, green, and acceptor signal, red). A zoomed-in region of the trajectory (middle) depicts several short events of ClpB and casein interaction. These events are seen as the sudden appearance and disappearance of spikes in the apparent FRET efficiency, *E*_app_ (= *N*_A_/(*N*_D_ + *N*_A_)), where *N*_D_ and *N*_A_ are donor and acceptor counts, respectively. Each spike reaches a maximum value of *E*_peak_ and is characterized by a time *τ*_dwell_. A cartoon (top) illustrates a translocation event of casein through the lumen of ClpB. (**C**) A map of the expected FRET efficiency (*ε*_calc_) for an acceptor-labeled segment as it passes through ClpB with a donor on the NBD1. The solid red line represents the centerline passing directly through the pore, while the black lines represent the boundaries of the pore. The colored regions correspond to the *N*-terminal domain (NTD, grey) and the two nucleotide-binding domains (NBD1 and NBD2, blue and purple, respectively). The secondary *y*-axis denotes the inter-dye distance (*d*).

We labeled casein on a native cysteine and ClpB on a cysteine inserted into the nucleotide binding domain 1 (NBD1) of only one of its six subunits (Figs. S5 and S6). A sample fluorescence trajectory containing multiple events of interaction between ClpB and casein in the presence of 2 mM ATP is shown in Fig. 1B. Most of the time, the trajectory shows no energy transfer between the dyes as ClpB and casein independently diffuse within the liposome. However, there are sudden, brief appearances of high FRET efficiency, indicating that casein has entered the lumen of ClpB (blue, Fig. 1 B inset). The peaks in FRET efficiency were characterized by the temporal length of the events (*τ*_dwell_) and the maximum FRET efficiency values reached (*E*_peak_). As illustrated at the top of Fig. 1B, the 76 nm (= 190 *aa* × 0.4 nm per *aa* (*30*)) contour length of casein far exceeds the ca. 13 nm length of ClpB’s lumen (hexameric model based on PDB 1QVR (*31*)) and the 6 nm Förster radius (*R*_0_) of the dye pair. Therefore, *τ*_dwell_ does not correspond to the translocation process of the entire chain of casein but rather to a temporal segment during which the acceptor-labeled portion of casein passes through ClpB. To help illustrate this, Fig. 1C presents a calculated map of the expected FRET efficiency (*ε*_calc_) as the acceptor traverses the central pore of ClpB. As the fluorescently labeled residue enters the *N*-terminal domain (NTD), *ε*_calc_ is less than 0.3, depending on the exact location within the cross-section of the 2 – 3 nm diameter pore. As the dye travels deeper into the lumen, *ε*_calc_ increases, potentially reaching a value close to unity around the interface between the NBDs. *ε*_calc_ then decreases as the labeled segment exits the trans side past the NBD2. Thus, *τ*_dwell_ values represent the time that it takes a contour length of ∼18 *aa*’s of casein to pass through ClpB (where *ε*_calc_ > 0.5), and *E*_peak_ corresponds to the minimum proximity between the two labels during an event.

### smFRET reveals ultrafast translocation of casein

From the sample trajectories given in Figs. 1B and 2A (ClpB labeled on the NBD1), it can be readily seen that the events occur very quickly, lasting only a few milliseconds. The 1535 events measured across 166 trajectories resulted in a power-law distribution of dwell times, 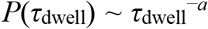, with a power *a* =1.69 ± 0.02 (Fig. 2B). Although the *τ*_dwell_ values spanned several orders of magnitude, the majority (89%) of the events occurred in less than 10 ms. Power laws with a power less than two do not have well-defined means (*32*), so an estimate of the average dwell time (<*τ*_dwell_>) in our experiments was obtained from an exponential fit to the short (< 10 ms) translocation times. The fit revealed a <*τ*_dwell_> value of only 1.6 ± 0.1 ms, nearly 3 orders of magnitude faster than the ATP hydrolysis time of the machine (1.14 ± 0.03 ATP/s, Fig. S5). To preclude the possibility that the short *τ*_dwell_ values were due to acceptor blinking (a photophysical artifact), we verified that the acceptor remained fluorescently active throughout the entire trajectory (Fig. S7). We also verified that, as demonstrated before, ClpB retained its hexameric assembly (Fig. S5) under these conditions, necessitating that casein can only enter ClpB through the lumen openings. Given that the millisecond *τ*_dwell_ values are much longer than the timescale of the dynamics of an unfolded protein chain (10’s – 100’s ns (*33*)), there is ample time for the chain segment to explore the entire cross-sectional area of the pore at any given point during the translocation process. For the sake of simplicity, we discuss the *ε*_calc_ values corresponding to a chain passing through the center of the lumen (red center line in Fig. 1C). Comparing the *E*_peak_ values (Fig. 2C) to the *ε*_calc_ map (Fig. 1C), it becomes clear that the moderate *E*_peak_ values of ca. 0.5 correspond to instances in which the tagged residue of casein only begins to enter the pore (> 3 nm depth). If entry is via the NTD, it indicates that casein has reached the NBD1 but has not made it past the first of three sets of flexible loops extending into the lumen (pore loops 1, PL1, located at a depth of ca. 3.4 nm). On the other hand, those 45% of the events with *E*_peak_ values ≥ 0.7 must originate from cases where casein reaches deep into the lumen of ClpB, near the interface between the two NBDs. This could be accomplished by casein entering through the NTD ring, reaching NBD1, and passing through (at least) PL1 and PL2 (the depth of the latter is ca. 4.4 nm), corresponding to a translocation depth of at least ca. 5 nm into the lumen.

**Figure 2.**
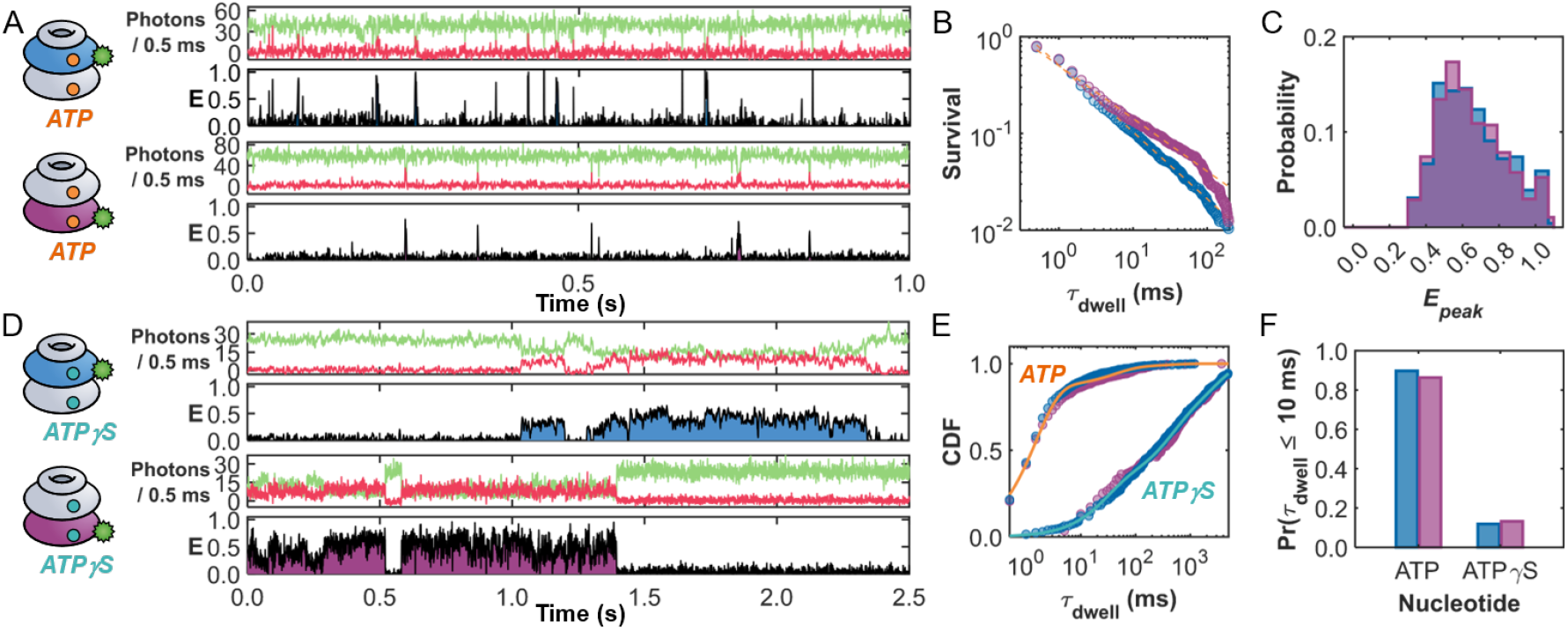
ATP, but not ATPγS, sustains rapid substrate processing. Sample trajectories in the presence of (**A**) 2 mM ATP or (**D**) 2 mM ATPγS. ClpB was labeled with a donor either on residue 359 in NBD1 (top trajectory) or residue 771 in NBD2 (bottom trajectory). (**B**)The survival probability functions of the events for NBD1-(blue) and NBD2-(purple) labeled ClpB are very similar, with power-law fits (dashed lines) yielding powers of 1.69 ± 0.02 and 1.55 ± 0.02, respectively. (**C**) Histogram of the peak FRET efficiency, *E*_peak_, for the events in (B). (**E**) Cumulative distribution function (CDF) of the event times for the two labeling locations in the presence of ATP and ATPγS. The solid lines represent triexponential fits of the dwell times in the presence of (orange) ATP and (teal) ATPγS, providing average values of 21.8 ± 0.3 and 749 ± 4 ms, respectively. (**F**) The presence of ATPγS suppresses the fraction of short (< 10 ms) events in both labeling locations, demonstrating that the fast translocation events are ATP-dependent.

To examine whether translocation events culminated in a complete passage through the lumen of ClpB, we repositioned the donor molecule to the NBD2 of ClpB (Fig. 2A, bottom). In this position, high FRET efficiency events can only occur if casein passes through the NBD2 ring (see Fig. S8). The sample trajectory shown in Fig. 2A bottom (in purple) is remarkably similar to those obtained with the NBD1-labeled ClpB (Fig. 2A top). In fact, the 1214 events measured across 137 trajectories displayed a similar power-law distribution (*a* =1.55 ± 0.02, Fig. 2B), and identical average event time (<*τ*_dwell_> = 1.6 ± 0.1 ms) and *E*_peak_ values (Fig. 2C) compared to those of the NBD1-labeled protein. Additional sample trajectories for both labeling locations can be found in Figs. S9 and S10, and a summary of statistics is provided in Tables S1 and S2. The high *E*_peak_ events in the histogram (Fig. 2C), constituting 45% of all events, necessitate that the acceptor-labeled segment has completed passage through the NBD2 and through the third ring of pore loops (PL3, depth of ca. 7.8nm). The low *E*_peak_ values of ca. 0.5 might correspond to events where casein has made its way past the first two PL rings but has not completed translocation across NBD2. The similar <*τ*_dwell_> values for NBD1- and NBD2-labeled ClpB suggest that the longitudinal motions (estimated to correspond to distances of 8.1 and 6.2 nm, respectively, based on the region in the *ε*_calc_ maps where *ε*_calc_ > 0.5), occur rapidly across both NBDs. While these fast events only represent snapshots of motion corresponding to contour lengths of ca. 16 – 20 *aa*’s along casein, the overwhelming frequency of these short events points towards a rapid translocation mechanism spanning both NBDs.

### Millisecond translocation is ATP-dependent

The threading velocity through ClpB (∼ 20 *aa*’s in 2 ms) estimated from the above experiments is much faster than expected based on its overall ATPase activity (1.14 ± 0.03 ATP/s, Fig. S5), begging the question as to whether the events measured here truly represent ATP-dependent translocation. While it would be tempting to simply remove ATP to abolish ClpB activity, the hexameric assembly of ClpB necessitates the presence of ATP or an analog (*34*). The slowly hydrolysable ATPγS (adenosine-5′-o-3-thio-triphosphate) is known to hamper the translocation activity of ClpB, resulting in a *holdase*-like behavior (*35*). We, therefore, tested events of interaction between ClpB and casein in the presence of 2 mM ATPγS, collecting 337 events from 147 molecules with the NBD1 labeling location and 227 events from 113 molecules with the NBD2 labeling location. Notably, compared to the same labeling locations in the presence of ATP, there was a > 4-fold reduction in the average number of events (compare to >1200 events for a similar number of molecules with ATP, Table S2). Fig. 2D provides sample trajectories for both labeling locations on ClpB (NBD1 in blue, NBD2 in purple) in the presence of 2 mM ATPγS, demonstrating a substantial increase in the substrate dwell time (see also Figs. S11 and S12). A cumulative dwell time histogram of both labeling locations in the presence of ATP and ATPγS is given in Fig. 2E. While the presence of ATP leads to nearly 90% of the events occurring in less than 10 ms (Fig. 2F), ATPγS suppresses these short events, reducing their frequency to < 15%. Fitting the cumulative distribution of dwell times (solid lines in Fig. 2E, Table S3) revealed that the weighted average overall dwell time increased from 21.8 ± 0.3 to 749 ± 4 ms when the nucleotide was switched from ATP to ATPγS, corresponding to an increase by a factor of ∼34.

In support of these findings, the apparent dissociation constants (*K*_D,app_) of casein in the presence of both nucleotides were determined by fluorescence anisotropy (Fig. S2). A simple bimolecular binding curve revealed that ATPγS decreased *K*_D,app_ (= 20 ± 10 nM) by a factor of 30 compared to that of ATP (= 600 ± 10 nM). Importantly, due to the active translocation and disengagement by ClpB, these *K*_D,app_ values do not represent the true dissociation constant but rather depend on the interaction times with casein. Therefore, the agreement between the 30-fold increase in dwell time and apparent equilibrium constants supports the conclusion that the events probed in the single-molecule experiments indeed report on the translocation activity of ClpB. Overall, the presence of ATP generates fast translocation events that last only a few milliseconds. Despite these events occurring on a timescale much faster than the bulk ATPase activity, the presence of ATPγS hinders them considerably, suggesting that with this nucleotide, substrates are not quickly released by the machine but rather dwell inside for long periods of time.

### Translocation times depend weakly on temperature

To glean further insight into the mechanism of translocation, the temperature dependence of the measured events was investigated by repeating the smFRET experiments at two additional temperatures, 10 and 32°C (see Methods for more details). At each temperature, over 60 ClpB molecules were sampled, leading to the observation of hundreds (> 500) of events (Table S1). Sample trajectories are provided in Fig. S13. For both labeling locations, the impact of temperature on *τ*_dwell_ was surprisingly small, leading to similar power laws, *E*_peak_ histograms, and fraction of short events (Fig. S14, Table S2). In fact, only a slight increase in the <*τ*_dwell_> value from 1.4 to 1.7 ms was observed as the temperature decreased from 32 to 10 °C (Table S2). An Arrhenius plot combining data from both labeling locations (Fig. 3A) revealed a very low activation energy for translocation of *E*_A_ = 1.87 ± 0.69 k_B_T. For comparison, the molecular motors kinesin and actomyosin, which consume ATP to ‘*walk’* along microtubules, exhibit significantly larger energy barriers of 20 and 40 k_B_T, respectively (*36, 37*). These classically established *power-stroke* machines exhibit large barriers due to large conformational changes driving the mechanical work (*14, 15, 38*), as would also be expected for ClpB if ATP were directly coupled with discrete translocation steps. Instead, the substantially lower activation energy measured for ClpB demonstrates that it must utilize a mechanism for translocation that does not rely on large structural shifts, such as a *Brownian ratchet* (*14, 39, 40*) (see discussion below).

**Figure 3.**
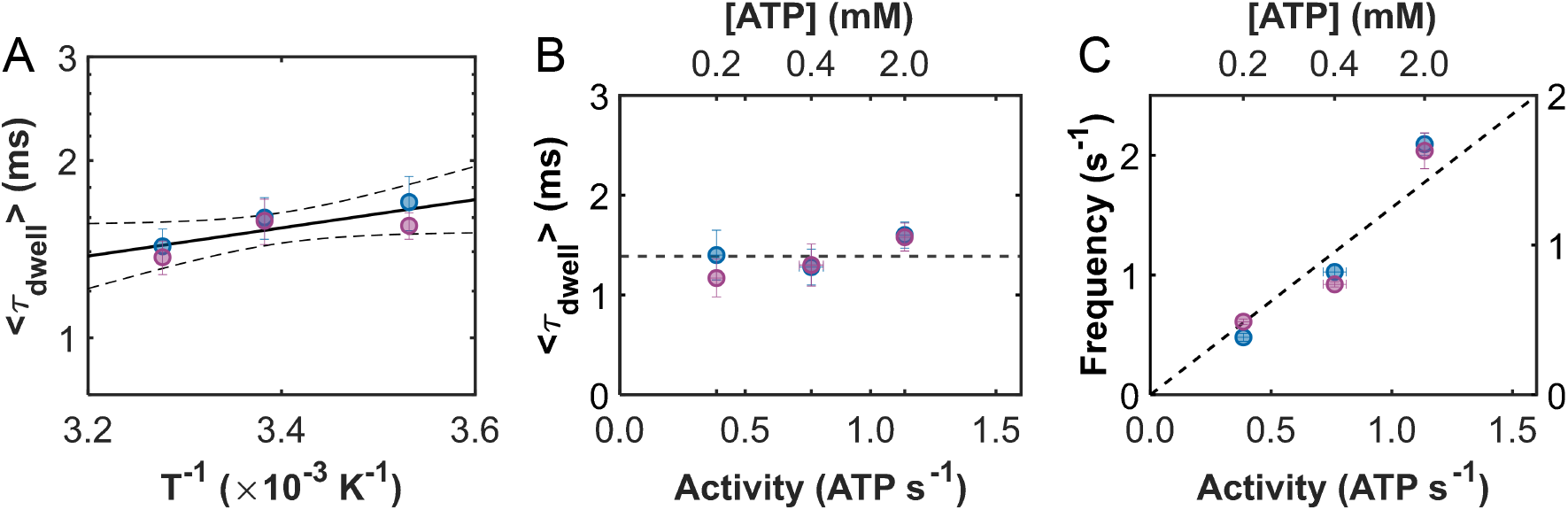
Temperature and ATP concentration dependence measurements support a ratcheting mechanism. (**A**) Arrhenius plot (logarithm of <*τ*_dwell_> vs. the reciprocal of temperature) of the average dwell times for NBD1- (blue) and NBD2-(purple) labeled ClpB. Linear regression (solid black line) returns an activation energy of 1.87 ± 0.69 k_B_T. The 95% confidence intervals are provided as dashed lines. (**B**) <*τ*_dwell_> values for NBD1- (blue) and NBD2-(purple) labeled ClpB as a function of ATPase activity. The dashed line represents the average dwell time of 1.4 ± 0.2 ms. The ATPase activity of ClpB (bottom axis, Fig. S5) was controlled by varying the ATP concentration (top axis) in the smFRET experiments. (**C**) The frequency of events for NBD1- (blue, left axis) and NBD2- (purple, right axis) labeled ClpB as a function of ATPase activity. Linear regressions return 1.65 ± 0.18 and 1.29 ± 0.15 events per hydrolysed ATP based on the bulk ATPase activity. Note the non-linear scale in ATP concentration in panels B and C, top axes.

### Event frequency directly correlates with bulk ATPase activity

At a saturating concentration of ATP (2.0 mM, Fig. S5), the bulk casein-stimulated ATPase activity of ClpB is only 1.14 ± 0.03 ATP/s. How then is the slow consumption of ATP related to the two to three orders of magnitude faster translocation events? To begin exploring this question, we repeated single-molecule experiments under conditions where the ATPase activity of ClpB was reduced. By lowering the concentrations of ATP from 2.0 to 0.4 and 0.2 mM, the bulk ATPase activity was lowered from 1.14 ± 0.03 to 0.76 ± 0.05 and 0.39 ± 0.03 ATP/s, respectively (Fig. S5). Translocation events from over 100 individual ClpB molecules were collected at each ATP concentration and labeling location (Table S1), and sample trajectories are shown in Fig. S15. Despite the significant reduction in ATPase activity with decreasing ATP concentration, we observed only small changes in the average dwell times (Fig. 3B), dwell time distributions, power-law exponents, and *E*_peak_ histograms (Tables S2 and S4, Fig. S16). There was, however, a significant reduction in the frequency of events (= # of events/trajectory length) as the ATPase activity decreased. Exponential fits of the event frequency distribution (Fig. S17) revealed that the frequency decreased linearly with decreasing activity (Table S5). As seen in Fig. 3C, both labeling locations exhibited very strong correlations (*R* = 0.98 and 0.95, respectively) between the event frequency and the bulk ATPase activity. The unanticipated result that decreasing ATP concentrations strongly affects event frequency and only weakly event dwell times reveals the important result that ATP controls the recruitment of protein substrates by the machine. Earlier studies on hexameric AAA+ machines suggest that ATP may widen the lumen entrance (*41, 42*), perhaps explaining the facilitation of substrate recruitment. From the slopes of the results in Fig. 3C, we find that the average numbers of events per hydrolysed ATP molecule are 1.65 ± 0.18 and 1.29 ± 0.15 for NBD1- and NBD2-labeled ClpB, respectively. The larger-than-unity values are perhaps not unexpected, since it is possible that substrate recruitment by ClpB might be affected by the dwell times of ATP products (ADP or P_i_) on one or more of its 12 binding sites, rather than the ATP hydrolysis events themselves.

### Three-color smFRET studies reveal bidirectional translocation

While the two-color smFRET experiments demonstrated that casein can rapidly translocate past both NBDs, the question remained if these events represented complete translocation. Further, while previous studies suggested that the interaction with substrates starts at the *N*-terminus, there is no direct proof that translocation can take place only in this direction. To address these aspects, we designed a three-color FRET experiment whereby a donor dye (Cy3B) was attached to the substrate casein, and two acceptor dyes were positioned on ClpB. Using bioorthogonal labeling techniques to ensure location specificity (see Methods), we labeled the NBD1 and NBD2 of ClpB with AF647 (acceptor 1) and CF680R (acceptor 2), respectively. The specific positions were chosen to maximize the sensitivity to the motion through the lumen while simultaneously minimizing unwanted energy transfer between the acceptors (Fig. S18). A map of the *ε*_calc_ in Fig. 4A depicts how the FRET efficiency between the dye on casein and each of the dyes on ClpB is expected to change as a function of position throughout the lumen. Given the relative locations of the dyes, acceptor 1 (orange) contributes more while casein traverses the NTD and NBD1, while energy transfer to acceptor 2 (red) becomes dominant as casein passes through the NBD2 (see Methods for details on FRET efficiency calculations). We collected 1445 signals of events across 135 ClpB molecules with ATP, and 135 events across 110 molecules with ATPγS. Sample trajectories are provided in Fig. S19. In the presence of ATP, the dwell-time distributions (decaying with a power law of *a* = 1.68 ± 0.2), the fraction of short events (91% of the events persisted for less than 10 ms), and *E*_peak_ (= max(*E*_app_)) values were all in close agreement with the two-color experiments (Fig. S20, Table S2). As expected, the addition of ATPγS significantly reduced the number of observed events and generated a 20-fold increase in dwell time (Fig. S21), demonstrating that the three-color experiments were sampling the same rapid ATP-dependent translocation events. However, with the addition of the second dye, we now had the ability to further dissect the results, providing unprecedented detail to the picture of translocation.

**Figure 4.**
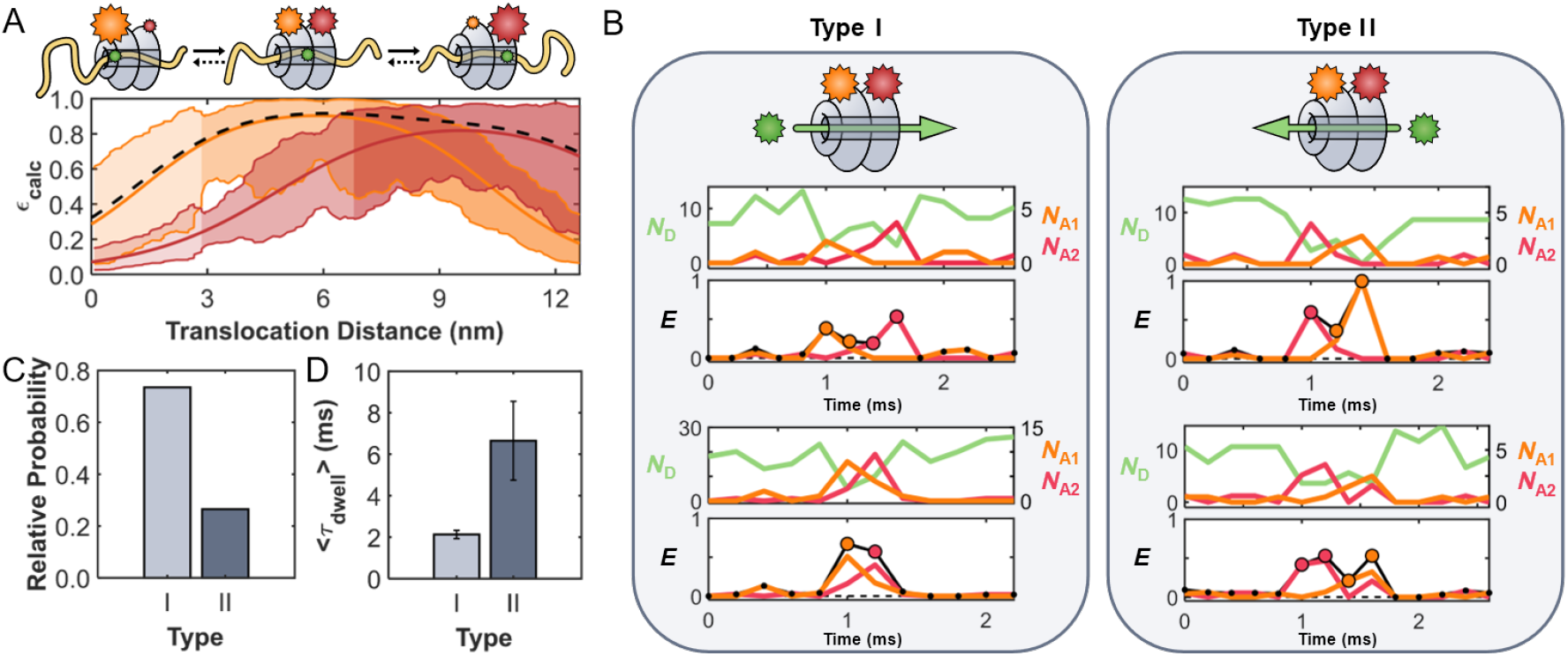
Translocation of casein in both the forward and backward directions is observed in three-color experiments. (**A**) Map of the calculated FRET efficiency as a function of location within the lumen of ClpB. Acceptors are located on the NBDs of ClpB (acceptor 1: AF647 (R_0_ ∼ 6.2 nm), orange, res. 176, and acceptor 2: CF680R (R_0_ ∼ 7.2 nm), red, res. 771). The thick orange, red, and black lines represent the calculated FRET efficiency through the center of the pore to acceptors 1 and 2, and the total FRET efficiency, respectively. The shaded areas represent the limits of the FRET efficiencies based on the boundaries of the pore. (**B**) Complete translocation events through ClpB. Type I – *forward* translocation events starting at the NTD and exiting past the NBD2. Type II – *backward* translocation events starting at the NBD2 and exiting past the NTD. Samples of the fluorescence of the donor (green, *N*_D_) and acceptors (acceptor 1: orange, *N*_A1_ and acceptor 2: red, *N*_A2_) and apparent FRET efficiency (*E*_A1_ = *N*_A1_ /(*N*_D_ + *N*_A1_ + *N*_A2_), *E*_A2_ = *N*_A2_ /(*N*_D_ + *N*_A1_ + *N*_A2_), and overall apparent efficiency as the black trace *E*_app_ = *E*_A1_ + *E*_A2_) are provided. Each point in time was assigned a color based on which acceptor was more prevalent (i.e., which acceptor experienced greater energy transfer from the donor, see Methods) as indicated by the coloring of the data points in the FRET efficiency trace. (**C**) The fractions of complete translocation events in the *forward* and *backward* directions when considering only complete translocation events, and (**D**) the average event times (Fig. S20).

Three-color FRET experiments uncovered a total of six types (I-VI) of translocation events. As shown in Fig. 4, translocation events often took place in the expected, *forward* direction (type I, starting at the *N*-terminus and proceeding to the *C*-terminus). Representative type I events in Fig. 4B, left, and S22 demonstrate that the FRET events begin with acceptor 1 (orange) and terminate with acceptor 2 (red), indicating that casein has completely passed through ClpB. An exponential fit of the *E*_app_ dwell time distribution of type I events revealed a <*τ*_dwell_> value of 2.1 ± 0.2 ms (Fig. 4d). While this value was very similar to the <*τ*_dwell_> values found in the two-color experiments, the three-color experiments confirm that complete translocation indeed occurs in such a short timeframe. Based on the placement of the two dyes on ClpB, the <*τ*_dwell_> values for translocation correspond to a lateral motion of 11.6 nm (which is the distance in Fig. 4A where *ε*_calc_ > 0.5), translating into an overall average translocation velocity of 14 *aa*/ms in the *forward* direction. Based on this estimate, the 190 *aa* long casein, corresponding to a contour length of ca. 76 nm, would take only ∼14 ms to be fully translocated through the lumen of ClpB. While not directly observed, both bulk stopped-flow mixing experiments (*24*) and single-molecule optical tweezer assays (*20*) support the existence of such fast translocation events. Although the short range of FRET limits our ability to rule out a combination of translocation steps and pauses, as proposed by single-molecule optical tweezer assays (*20, 43, 44*), the lack of intermediate FRET efficiency values in our data points towards a continuous translocation mechanism. In fact, sustained rapid translocation with speeds greater than 2.8 nm/ms (equivalent to ∼ 7 *aa*/ms) has been observed in DNA-translocating AAA+ proteins (*45*).

In addition to *forward* translocation events, there were also events that proceeded in the unanticipated *backward* direction (type II, starting at the *C*-terminus and continuing to the *N*-terminus, Fig. 4B, and Fig. S23). Of all complete translocation events, 25% occurred in the *backward* direction, where casein entered ClpB from the NBD2 side (generating fluorescence in the second acceptor, marked red), moved past all three sets of pore loops, and exited through the NTD (generating fluorescence in the first acceptor, marked orange). While it has been assumed so far that translocation occurs only in the forward direction, this study not only reveals the existence of *retrograde* threading events but also demonstrates that they occur in relative abundance, highlighting their significance in the translocation activity of ClpB. These type II events differed from their type I counterparts in their <*τ*_dwell_> value of 6.6 ± 1.9 ms (Fig. S20), which was about three times slower, pointing to a preferential *forward* bias in motion through ClpB’s lumen.

### Partial threading events

In addition to complete *forward* and *backward* translocation, the three-color smFRET experiments also identified partial translocation events. Types III and IV events in Fig. 5A (also in Fig. S24) correspond to incomplete translocation events where casein reached deep into the lumen, arriving at both NBDs and thus exhibiting energy transfer to both acceptors. However, the signals began and terminated with the same acceptor, demonstrating that casein entered and exited from the same side of ClpB. While partial threading events have been inferred from previous studies (*21, 46*), we directly confirm their existence, also noting that they occur in a bidirectional manner. As in the case of type I and II events, a notable feature is that the *forward* type III events were significantly more abundant than *backward* type IV events (Fig. 5B). Type III events typically took 11.3 ± 3.1 ms, over 5 times longer than the type I events (Fig. 5C). However, the *backward* type IV events, with a <*τ*_dwell_> of 6.2 ± 0.5 ms, lasted for nearly identical times as the *backward* type II complete translocation events. Another interesting feature is that the relatively long dwell times of both type III and IV events appear to originate from rapid back-and-forth motions of casein, as seen in the frequent interconversion between the two acceptors (orange and red) in the sample trajectories given in the top of Fig. 5A. Such a feature suggests that the long dwell times of these events are related to back-and-forth motions of the substrate in the lumen before exiting. A correlation plot between *τ*_dwell_ and the number of back-and-forth motions revealed a correlation coefficient of *R* = 0.982 (Fig. 5D).

**Figure 5.**
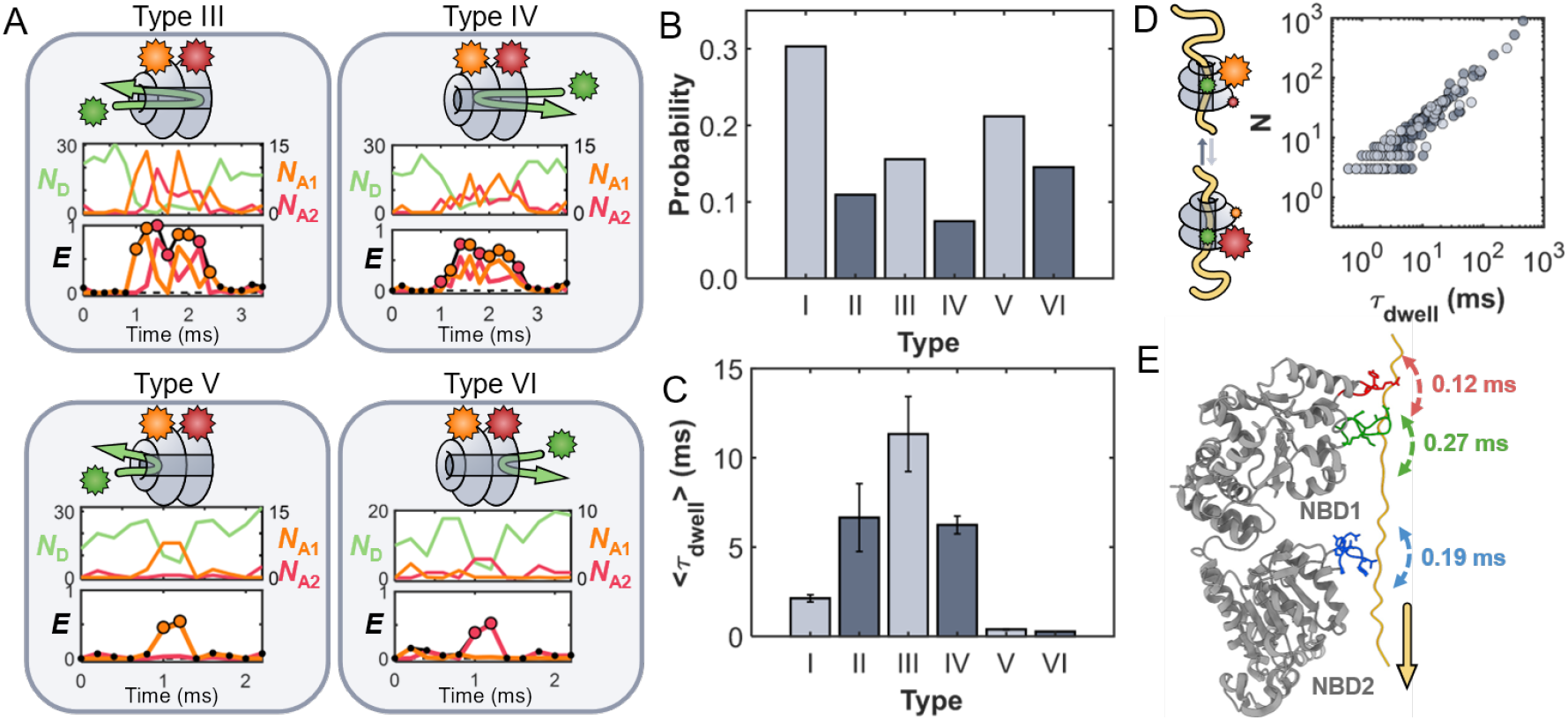
ClpB exhibits partial translocation events. (**A**) Sample incomplete translocation events. Type III (*forward*) and IV (*backward*) partial translocation events are characterized by casein reaching both NBDs but exiting from the same side it entered from. Type V (*forward*) and VI (*backward*) fleeting events are characterized by an interaction with only a single NBD. (**B**) Probability of occurrence of the different translocation events. (**C**) Average time for each event type (Fig. S20). (**D**) Correlation between the number (*N*) of back-and-forth motions and the dwell times for type III (light) and IV (dark) events. The correlation coefficient of *R* = 0.98 demonstrates that the long dwell times are directly proportional to *N*. The cartoon inlay illustrates the back-and-forth motions of casein. (**E**) Schematic of the previously determined sub-millisecond dynamics of the pore loops (PLs) in the presence of ATP and casein (*4*). The three PLs (red, green, and blue) on each subunit directly engage the substrate and rapidly move up-and-down (PDB 6OAX (*5*)). These fast dynamics modulate the energy landscape between two (or more) rapidly interconverting states. ATP allosterically regulates the dynamics and conformational states of the PLs (*4*), generating a landscape in which the substrate preferentially diffuses *forward* from NBD1 towards NBD2.

Lastly, type V (*forward*) and VI (*backward*) events (Figs. 5A and S25) were attributed to casein superficially binding, but not entering the lumen, resulting in the appearance of energy transfer to only one of the acceptors. Both type V and VI events exhibited very brief dwell times of under 0.5 ms (Fig. 5C), suggesting that these events represent instances where casein attempts to enter the lumen but fails to make it past one of the NBDs (which would result in energy transfer to both acceptors) before dissociation.

Overall, only two-thirds of all events started at the *N*-terminus of ClpB, while the rest started at the *C*-terminus. It is possible that in the presence of ClpB’s co-chaperones, such as DnaK and DnaJ, which are typically required for aggregate recognition, a larger fraction of translocation events would originate from the *N*-terminal side (*6*). However, even without these co-chaperones, the NTD binding groove on ClpB is known to bind substrates such as casein (*47*), and therefore likely contributes to the observed two-fold increase in probability for events that begin at the *N*-terminus.

In the presence of ATPγS, we found that there was a five-fold reduction in the number of events per ClpB molecule, similar to the four-fold reduction in interactions observed in the two-color experiments. Fitting the overall cumulative distributions of dwell times (Fig. S26), it was found that ATPγS increased the overall dwell time by a factor of 20 compared to that of ATP. The lower factor than the one found in the two-color and fluorescence anisotropy experiments (∼30) can be attributed to premature photobleaching of the donor before substrate disengagement due to the increased laser power used in these experiments (see Methods). Despite the limited number of only 135 events (across 110 ClpB molecules) registered in the presence of ATPγS, we nonetheless found that ClpB exhibits all six types of events. However, the fractions of *forward* and *backward* events were nearly equivalent to each other in all cases, indicating that with ATPγS, the *forward* vs. *backward* bias vanishes. Importantly, this result demonstrates that ATP is necessary to rectify translocation in the *forward* direction. A complete summary of the frequency of each event type alongside their corresponding dwell times can be found in Tables S6 and S7. We also note that the strict nucleotide dependence and the orderly sequence of FRET signals observed in the three-color experiments further confirm that we are monitoring genuine substrate translocation events through the ClpB lumen, rather than via side entry or nonspecific interactions at the protein surface.

## Discussion

### ClpB drives diffusive-like translocation

The translocation activity of ClpB demonstrated in this work is surprisingly rich and very different than expected based only on structural studies of AAA+ machines. Translocation events are short and demonstrate a preferred directionality. These characteristics are abolished in the presence of ATPγS, indicating significant dependence on ATP. The bidirectional nature of translocation suggests that the energy landscape throughout the translocation process must be shallow. Indeed, our temperature study revealed an activation energy of only ∼ 2 k_B_T. The experiments further demonstrate that ClpB facilitates both partial and complete translocation events in a bidirectional manner, albeit with a significant preference for *forward* motion.

We can gain deeper insight into the landscape by considering the dwell times and frequency of the different types of events. First, we find that the complete *forward* translocation events are, on average, over five times faster than the incomplete or *backward* translocation events. Based on this ratio, transition state theory (*48*) suggests that the difference in free energy between the *forward* and *backward* threading directions (*ΔΔG* = *ΔG*_forward_ – *ΔG*_backward_ = –ln(<*τ*_dwell_>_forward_ / <*τ*_dwell_>_backward_)) is no greater than 1 – 2 k_B_T. Second, considering the scenario where casein has entered the lumen (i.e. not including the superficial types V and VI events), we find that casein is only slightly more likely to exit past NBD2 (Pr(‘*exit via NBD2’*) = 0.59, types I and IV) than past the NTD (Pr(‘*exit via NTD’*) = 0.41, types II and III). The Curtin–Hammett principle (*49*) suggests that the difference in transition-state energies (ΔΔ*G*^‡^ = –ln(Pr(‘*exit via NBD2*’)/Pr(‘*exit via NTD*’)) between the *forward* and *backward* directions is less than 1 *k*_B_*T*.

Our experiments paint a picture of a shallow landscape where, at any given time, casein is nearly as likely to diffuse in either direction. Further evidence for diffusive-like translocation arises from the type III and IV events, which exhibit hallmarks of rapid back-and-forth motions of casein within the lumen and show a direct correspondence between the number of these motions and the duration of the events. Notably, we observe that diffusion must occur quickly, resulting in translocation within just a few milliseconds. Remarkably, limiting the supply of ATP did not significantly alter the translocation time but rather reduced the frequency of translocation events, indicating that ATP must regulate substrate recruitment in addition to translocation directionality and velocity. Moreover, we find that there are, on average, ∼ 1.5 events per hydrolysed ATP. Based on the 41% probability of complete threading, we calculate that each complete threading event correlates with the consumption of only 1.65 ± 0.29 ATP molecules. Together, these features suggest a mechanism where the utilization of just one or two ATP molecules is sufficient to drive rapid bidirectional translocation events. A fully detailed understanding of this feature requires knowledge about the average dwell time of ATP and its hydrolysis products on all 12 binding sites of ClpB, which is lacking at this point in time. It is important to note that ClpB activity may involve both releasing a protein from an aggregate and the subsequent translocation through the central pore, potentially requiring distinct mechanisms of action (*1, 24*). Our experiments with the intrinsically disordered casein focus on the latter and isolate the translocation process in the absence of any potential rate limitation due to the disaggregation step. While our study does not directly probe substrate unfolding, we emphasize that translocation is an essential and indispensable component of the overall disaggregation mechanism.

### ClpB as a Brownian motor

Two mechanisms have been proposed for molecular machine driven translocation: the *power-stroke* and the *Brownian motor* (*14, 15, 40, 50-52*). While these two mechanisms can share some nuanced characteristics (*50*), they are classically quite different from one another. The *power-stroke* mechanism involves significant, fuel-driven conformational changes in the machine that generate irreversible translocation steps along a downhill free energy landscape. In contrast, a *Brownian motor* does not depend on large, coordinated motions. Instead, substrate movement is driven by thermal noise (Brownian motion), while an external energy source introduces some form of asymmetrical changes within the free energy landscape. This asymmetry restricts diffusion to one direction, resulting in directed motion. The mechanistic features utilized by ClpB discussed above align more closely with those expected of a *Brownian motor* (*14, 40*). Notably, the rapid translocation by ClpB is more akin to findings on transmembrane pores (e.g., α-hemolysin (*53*) and Cytotoxin-K (*54*), both with a similar 2 nm pore diameter to ClpB), though in these cases, electric potential differences drive biomacromolecules across a membrane. We note that in our experiments, ClpB’s central pore (∼13 nm in length) is much shorter than the full contour length of casein (∼76 nm), and casein is labeled on a central cysteine. Consequently, the early stages of threading, where work against the entropy of casein is expected to be greatest and therefore may slow translocation, have limited resolution. However, if entropic effects were a major hurdle, we would expect long dwell times at low FRET efficiencies (E ∼ 0.2, based on Figs. 1C and 4A), due to binding at the entrance of ClpB’s pore, which are absent in our data. Instead, we find that dwell times longer than 10 ms are rare (<15%) and, when present, show FRET efficiencies similar to the rapid events. Thus, while an entropic barrier (expected to be on the order of ca. 9 k_B_T (*55*)) may hinder binding or the earliest threading steps, we have no indication that it significantly impacts the overall translocation time, though it may lower the overall number of interaction events.

Similarly, our findings do not exclude the possibility of subunit motions predicted by structural models, but they do, however, demonstrate that these motions alone cannot fully account for the translocation mechanism. This raises a fundamental question: How does ClpB harness the energy from ATP to drive directional translocation? Our earlier studies (*4, 56*) revealed that the substrate-engaging pore loops (PLs), which line ClpB’s lumen, exhibit sub-millisecond up-and-down motions (Fig. 5E), suggesting a dynamic interaction that may regulate substrate translocation. Importantly, these stochastic ultrafast PL dynamics persist under all nucleotide conditions and even in the presence of casein. However, the populations of the PL conformational states are allosterically regulated by the nucleotide, leading to nucleotide-dependent asymmetries in their orientations (*4*). Within this framework, we propose a ratcheting mechanism in which the substrate encounters an energy landscape with a preferred direction generated by PL dynamics. The rapid and asymmetric interconversion between PL states facilitates the rapid motion of casein within the lumen while rectifying translocation in the *forward* direction. This interpretation is consistent with prior findings on the AAA+ protease FtsH, where thermal motions were shown to drive subunit dynamics, and ATP acted to bias the conformational probability distribution (*57*). Together, these observations reinforce the view that ATP functions not as a direct power source for large conformational strokes, but rather as an asymmetry generator that rectifies Brownian motion into productive translocation.

To explore whether the previously observed sub-millisecond dynamics of the PLs correlate with the millisecond translocation events observed in this study, we calculated the average time and amplitude (of motion) for each of the three sets of pore loops to complete their up-and-down motions in the lumen (Table S8). As depicted in Fig. 5E, PLs 1, 2, and 3 take an average time of 0.12, 0.27, and 0.19 ms, respectively (*4*), to complete one up-and-down lateral motion of 1.4 – 1.7 nm (*56*). By multiplying the time to complete one up-and-down ‘cycle’ by the amplitude of motion, we find that the PL dynamics correspond to motions of 16 – 28 *aa*/ms (for PL2, (1.7 nm amplitude of motion) × (1 *aa* per 0.4 nm) / (0.27 ms per ‘cycle’)). These fast motions represent the expected limiting speed for a substrate traversing the rapidly evolving PLs and closely align with the ∼ 10 *aa*/ms velocity estimated from the two-color experiments, as well as the 14 *aa*/ms velocity found for the type I events. In the presence of ATPγS, the PLs remain dynamic, but they lose their asymmetry (*4*), thereby abolishing the asymmetric landscape required for rapid directional translocation. Lastly, simulations demonstrate that rapid motions of asymmetrical restrictions in a pore, such as the up-and-down movements of the PLs, produce directional translocation as a result of entropic forces (*58, 59*), supporting our model. It is noteworthy that entropically driven translocation is weakly dependent on temperature (*60*), further agreeing with our findings.

## Conclusions

This study demonstrates that substrate translocation through the AAA+ machine ClpB occurs within milliseconds. ATP is essential for these rapid translocation events, with a limited supply of ATP reducing translocation frequency but not velocity, underscoring its role in substrate recruitment rather than direct force generation. Crucially, we reveal that translocation operates bidirectionally, with a 3:1 preference for *forward* threading. Substituting ATP with ATPγS abolishes directionality and hinders translocation, reinforcing ATP’s essential role in regulating movement through ClpB’s lumen. Our findings suggest that translocation cannot be exclusively governed by discrete power strokes following ATP hydrolysis. Instead, the observed diffusive-like behavior and low energy barriers associated with bidirectional motion support a mechanism akin to a *Brownian motor*, where ATP facilitates directionality rather than generating discrete steps. *Brownian motors*, which use an energy source to rectify diffusive motion (*14*), have been broadly studied as physical models for dissipative dynamics (*61*), yet direct observation of such motors in the context of biological nanomachines remains rare. Our results with ClpB may have broader implications for understanding other biological machines that couple ATP hydrolysis to substrate translocation, including protein disaggregases, unfoldases, and translocases across diverse cellular systems. By directly observing the interplay between translocation dynamics and ATP, this study establishes a framework for exploring mechanochemical coupling in other AAA+ proteins and beyond. Future studies using folded substrates will be important to directly probe the mechanochemical coupling between unfolding, translocation, and ATP. Such experiments would build on our current findings by revealing how force generation and substrate remodeling contribute to directional translocation in physiological contexts.

## Materials and Methods

### Protein expression and purification

*Thermus thermophilus* ClpB with a six-residue histidine tag at its *C*-terminus was cloned into a pET28b vector. All ClpB mutants were generated by site-directed mutagenesis and standard cloning techniques and were verified by DNA sequencing. The wild-type (WT) and single mutant ClpB proteins (359C and 771C) were expressed and purified as previously reported (*31*). A double-mutant ClpB incorporated the unnatural amino acid *N*-ε-(prop-2-ynyloxycarbonyl)-*L*-lysine (ProK, Iris Biotech) at residue 771, and a cysteine at residue 176. ProK was coded with the Amber stop codon, allowing for co-transfection of the double mutant 176C + 771TAG ClpB in pET28b together with pEvol PylRS/tRNApyl (*62*) (Addgene, Plasmid #127411, back mutated to A346N/A348C) into *E. Coli* BL21 (DE3) pLysS cells (Invitrogen) using heat shock. Cells were recovered and incubated in 10 mL Luria Broth (LB, Thermo Fisher) media containing kanamycin (Sigma, 50 μg/mL) and chloramphenicol (Sigma, 25 μg/mL) for 16 h at 37 ºC, 250 rpm. The overnight culture was used to inoculate 1 L of LB supplemented with corresponding antibiotics and arabinose (Formedium, 0.2 w/v%). After reaching an optical density (OD) of 0.5, the temperature was reduced to 25°C, and ProK (2 mM) was added. After reaching an OD of 0.8, ClpB expression was induced by the addition of isopropyl ß-D-1-thiogalactopyranoside (Sigma, 1 mM) and incubated for 24 hours. Purification was conducted in a similar manner as above. Briefly, cells were harvested and lysed using a French press, followed by heat shock treatment (80°C, 10 min.). ClpB was recovered from the supernatant on a Ni Sepharose column (HisTrap FF, Cytiva), and dialyzed (50 kDa molecular weight cutoff, Gene Bio-Application Ltd.) overnight.

### Protein labeling and subunit mixing

Single-mutant ClpB was labeled with cyanine 3B maleimide (Cy3B, Cytiva) following a previously reported procedure (*31*). κ-casein (Sigma) was labeled in a similar manner with either Cy3B or the cyanine-based dye 660R maleimide (CF660R, Biotium). The probability of labeling casein (containing two native cysteines) with two dyes was minimized by reducing the dye ratio to 0.6 eq. per protein molecule (see *distribution of labeled casein inside liposomes* in the SI for more details). Excess dye was removed using a desalting column (Sephadex G25 Superfine, Cytiva), followed by overnight dialysis (8 kDa, Gene Bio-Application Ltd.) against HEPES buffer (25 mM HEPES (J.T. Baker), 25 mM KCl (Sigma), 10 mM MgCl_2_ (Sigma), pH 8). The ClpB double mutant was first labeled with Alexa Fluor 647 maleimide (AF647, Thermo Fisher), after which the unnatural amino acid was reacted with CF680R azide (Biotium) through copper-mediated click chemistry. Briefly, CF680R (4 eq.) was added to a solution of AF647 labeled ClpB (500 μL, 13 μM) in phosphate buffer (Sigma, 100 mM, pH 7). Next, CuSO_4_ (Sigma, 200 μM) and tris-hydroxypropyltriazolylmethylamine (Lumiprobe, 1 mM) were added. The labeling reaction was then initiated with the addition of sodium ascorbate (Sigma, 5 mM) and the protein was left to react for 2 hours. Afterwards, the reaction was quenched by the addition of ethylenediaminetetraacetic acid (EDTA, 2 mM). ClpB was separated from the reaction mixture using a desalting column (HiTrap, Cytiva). Any remaining copper or small molecules were removed by overnight dialysis (50 kDa) against a HEPES buffer containing EDTA (2 mM).

Hexameric assemblies of ClpB containing at most one single fluorescently labeled subunit were prepared by reassembling a mixture of labeled ClpB (1 eq.) with an excess (20 eq.) of wild-type ClpB. Homogeneous mixing of the subunits was ensured by performing the procedure in the presence of the guanidinium chloride (GdmCl, 6M, Sigma) overnight. The denaturant was slowly removed by repeated dialysis (50 kDa) against HEPES buffers containing 4, 2, 1, 0.5, and 0 M GdmCl in four-hour intervals. The reassembled subunits were then dialyzed further against a HEPES buffer containing ATP (Sigma, 2 mM). The samples were then filtered (0.1μm, Whatman Anotop), aliquoted, flash-frozen, and stored at –80°C until further use. The low salt of the HEPES buffer (25 mM HEPES, 25 mM KCl, 10 mM MgCl_2_, pH 8) and the persistence of nucleotide after reassembly ensured that ClpB remained in a hexameric assembly in all experiments (*31, 63, 64*). Native gel demonstrating the reassembled ClpB is provided in Fig. S5. The rotational freedom of the covalently bound dyes are quantified in the *fluorescence anisotropy decay experiments* section in the SI.

## ATPase Activity

The ATPase activity of ClpB was measured according to a previously published procedure (*31*) using a coupled colorimetric assay (*65*). Briefly, ClpB (ca. 1 μM) was incubated in a buffer (25 mM HEPES, 25 mM KCl, 0.01% tween-20, pH 8) with an enzymatic ATP regeneration system (2.5 mM phospheonol pyruvate (Sigma), 10 units/mL pyruvate kinase and 15 units/mL lactate dehydrogenase (from rabbit, Sigma)), to which were added 1,4-dithioerythritol (2 mM, Sigma), EDTA (2 mM), nicotinamide-adenine dinucleotide (NADH, reduced form, 0.25 mM), and ATP (varied from 0 to 2 mM) at 25°C. The reaction was started by the addition of MgCl_2_ (12 mM). ATP hydrolysis was measured by monitoring the decrease of NADH absorption (340 nm) in time on a microplate reader (CLARIOstar, BMG Labtech). The ATPase activity of ClpB in the presence of casein (3 μM) and in its absence is shown in Fig. S5.

### Surface-tethered lipid vesicle preparation

Following a previously published procedure (*66*), the fluorescently labeled protein molecules were encapsulated inside unilamellar vesicles, which were subsequently immobilized on a glass-supported lipid bilayer through a biotin-streptavidin linkage. Briefly, vesicles used for the formation of a glass-supported lipid bilayer were prepared from a lyophilized mixture (5 mg) of egg-phosphatidylcholine (Avanti) and 0.2 mol% 1,2-dioleoyl-sn-glycero-3-phosphoethanolamine-*N*-(biotinyl) (18:1 biotinyl PE, Avanti), which was reconstituted into a HEPES buffer (1 mL) by mixing. The suspension was extruded 75 times through a 0.1 μm filter (Whatman Anotop) to form unilamellar vesicles. The vesicle solution was inserted into a glass flow cell for 15 minutes, then rinsed with HEPES buffer, leaving behind a supported lipid bilayer on the interior surfaces of the cell. The cell was flushed with a solution of streptavidin (1 mg/mL), which attached to the biotinylated lipids. Vesicles containing ClpB and casein were prepared by reconstituting a lyophilized mixture (3 mg) of 1% mol biotinyl PE in 1,2-dimyristoyl-sn-glycero-3-phosphocholine (DMPC, Avanti) with a HEPES buffer solution containing ATP (2 mM), ClpB (200 nM), casein (3 μM), ascorbic acid (AA, 2 mM, Sigma) and protocatechuic acid (PCA, 1.5 mM, Sigma). The mixture was then extruded 75 times through a 0.1 μm filter to generate unilamellar vesicles with a ca. 120 nm diameter. Unencapsulated casein was removed by passing the liposome mixture through a MicroSpin S-400 HR column (Cytiva). Importantly, while the initial sample contained a 150-fold excess of casein, this ratio was carefully chosen so that the resulting liposomes would only contain a single molecule of casein and be sparsely populated with ClpB, leading to a near 1:1 ratio between the proteins in the ClpB-containing liposomes (see *Calculation of the expected number of proteins inside a liposome* in the SI). The vesicle solution was diluted (1:100) and incubated inside the lipid-bilayer coated flow cell for 5 minutes to allow binding to the immobilized streptavidin. Unbound vesicles were removed by washing (3 x 200 μL) with the HEPES buffer containing ATP (2 mM, adjusted for experiments conducted at different concentrations). The HEPES buffer (100 μL, outgassed with *N*_2_ to remove oxygen) with the addition of an oxygen-scavenging system (*67, 68*) (1.5 mM PCA and ca. 5 nM protocatechuate 3,4-dioxygenase), a triplet quenching and radical scavenging system (*68, 69*), (2 mM AA, 1 mM methyl viologen) and ATP (2 mM, adjusted for experiments conducted at different concentrations) was added to the flow cell before sealing with silicone grease. Experiments in the presence of ATPγS (Roche, 2 mM) were prepared in a similar manner but were left to incubate with ATPγS for 40 minutes before reconstitution into liposomes.

### smFRET data acquisition

Single-molecule fluorescence data were acquired using a PicoQuant Microtime 200 confocal microscope. Two-color measurements were conducted in a continuous wave mode using a circularly polarized 561 nm laser with a power of ca. 400 nW (polarization and power were measured at the back focal plane of the objective (Olympus, UPlanFLN 100X/1.30 Oil Iris)). The positions of liposomes containing fluorescently labeled species were identified by raster scanning a piezo-driven stage (Physik Instrument, P-721.CLQ and P-733.2CL), and the fluorescence of individual liposomes was monitored until photobleaching of the donor dye. The instrument was equipped with a ZT405/488/561/640rpcv2 (Chroma) primary dichroic mirror, the emission was split between the donor (Cy3B) and acceptor (CF660R) channels using a T635lpxr dichroic mirror (Chroma) and filtered using a 593/46 Brightline (Semrock) or a 731/137 Brightline bandpass filter (Semrock).

To increase the time resolution for the three-color experiments, we elected to perform the experiments under increased laser power (from 400 to 700 nW), which nevertheless still allowed obtaining fluorescence trajectories of a reasonable length. Three-color experiments were conducted using 40 MHz pulsed lasers with a sequence of one pulse of the acceptor laser (640 nm, ca. 50 nW) followed by three pulses of the donor laser (530 nm, ca. 700 nW). The excitation was circularly polarized, as measured at the objective. The positions of liposomes containing labeled ClpB were identified using an acceptor-only raster scan with a power of ca. 500 nW. The instrument was equipped with a TZ532/640rpc primary dichroic mirror (Chroma), and the emission was split into donor and acceptor by a T635lpxr dichroic mirror (Chroma). The acceptor emission was further split by a T685lpxr dichroic mirror (Chroma) into two acceptor channels (AF647 and CF680R). Before reaching the detector, the donor emission was filtered with a 582/75 Brightline bandpass filter (Semrock). The acceptor channels were filtered with a ET667/30m or a ET720/60m bandpass filter (Chroma) for the AF647 and CF680R channels, respectively.

### smFRET data analysis

The apparent FRET efficiency (*E*_app_ = *N*_A_ / (*N*_D_ + *N*_A_)) of the two-color experiments was calculated using the photon counts (binned to 0.5 ms) from the acceptor (*N*_A_) and donor (*N*_D_) after corrections for background photons and channel crosstalk (i.e. acceptor emission detected in the donor channel and donor emission detected in the acceptor channel). Background flux was determined by the residual photon count in each channel after photobleaching of the donor, and channel crosstalk was found from trajectories of isolated donor- and acceptor-labeled molecules within liposomes. The acceptor had negligible crosstalk, while the donor had 13% of its emission appear in the acceptor channel. Further discussion on channel crosstalk can be found in the *derivation of leak and background corrections* and the *leak-factor determination* sections of the SI. Translocation events in the trajectories were identified by peaks in the *E*_app_ trajectories where *E*_app_ exceeded 0.3. Since this method did not account for the rising and falling edges of the events, the edges were extended until the *E*_app_ value was no longer statistically different (1.5 σ) than the local background values (using a sliding window of 7 bins). The dwell time *τ*_dwell_ of each event was taken as the length of each agglomerated region. *E*_peak_ values were taken as the maximum *E*_app_ value from the identified events.

While an explicit calculation of bona fide FRET efficiency values in multicolor experiments can be performed (*70*), for the purpose of this work, we elected to calculate apparent FRET efficiencies in simpler terms, directly using the experimentally obtained photon count rates of the donor (*N*_D_) and the two acceptors (*N*_A1_ and *N*_A2_) (*70, 71*). The apparent FRET efficiencies in three-color experiments were calculated using the photon counts from 0.2 ms bins after background correction. Due to the relatively low photon counts (∼ 10 – 30 photons / bin), applying additional correction factors such as channel crosstalk to the individual trajectories introduced a large amount of unwanted noise and, therefore, was avoided. Further discussion on the correction factors, including inter-acceptor energy transfer, can be found in the *leak-factor determination* and the *determining inter-acceptor energy transfer* sections of the SI. The apparent *E*_app_ to each acceptor was calculated as *E*_Ai_ = *N*_Ai_ / (*N*_D_ + *N*_A1_ + *N*_A2_), where Ai represents acceptor 1 or 2, and the total apparent FRET as *E*_app_ = (*N*_A1_ + *N*_A2_) / (*N*_D_ + *N*_A1_ + *N*_A2_). Following a similar procedure as in the two-color analysis, interaction events were identified using the total *E*_app_ values, such that the extracted *τ*_dwell_ values represent the total observed dwell time regardless of which acceptor is more prominent. Once identified, each bin in an event was then assigned a color based on which acceptor was most prominent. While we elected not to apply spectral crosstalk corrections directly to the trajectories, acceptor 1 (AF647) had a moderately large fraction of its emission (37%) appear in the acceptor 2 channel (CF680R). As such, a bin was assigned to acceptor 2 if *E*_A2_ contributed over 60% of the total *E*_app_; otherwise, it was assigned to acceptor 1. The *E*_A2_/*E*_app_ > 0.6 cut-off value was chosen because it corresponds to the situation where the donor is experiencing equivalent energy transfer to both acceptors (i.e., after correction for crosstalk, *ε*_A1_ ∼ *ε*_A2_).

### Event dwell-time characterization

The length of each event was characterized by the value *τ*_dwell_ as discussed in the *smFRET data analysis* section. To determine the exponent α of the power law distribution of *τ*_dwell_ values, 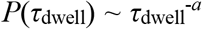, the survival probability that *τ*_dwell_ has a value greater than *τ*, S(*τ*), was calculated for each point in the distribution, and eq. 1 was fitted to the resulting data (*32*). In this equation, *P* is the probability distribution of *τ*_dwell_ values, *C* is a normalization constant, and α is the power for *τ*_dwell_ values obeying the power law (*32*)

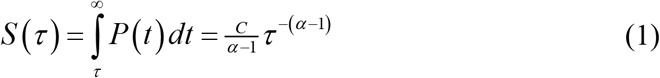

Distributions obeying power laws with α values less than two, as those reported herein, have a divergent mean and variance. Therefore, a representative overall average event time based on the collected statistics was found from the triexponential fit of the cumulative distribution (CDF, eq. 2) of *τ*_dwell_ values.

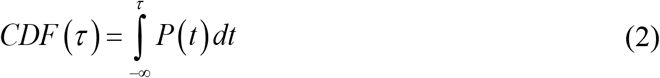

In the presence of ATP, the short dwell times, representing the vast majority of events (see Table S2), were characterized by an exponential fit of all events occurring in less than 10 ms. In doing so we noted that the few very long interactions did not spuriously skew the average dwell time reflecting the bulk of the short translocation events. Model parameters (and their corresponding standard errors) were optimized with the ‘fitnlm’ function in Matlab utilizing the Levenberg-Marquardt nonlinear least squares algorithm.

### Temperature-dependent smFRET experiments

In these experiments, the temperature inside the fluorescence cell was controlled by equipping the confocal microscope with a custom-built Peltier temperature unit, consisting of four Peltier elements controlling the temperature of an aluminum housing surrounding the cell, and another Peltier element controlling the temperature of an aluminum block directly coupled to the objective. The temperature inside the confocal volume was determined by measuring the change in the diffusion coefficient of Cy3B using fluorescence correlation spectroscopy (FCS). Further information can be found in the *fluorescence correlation spectroscopy* section and Fig. S27 in the SI. The constituents of the vesicular membranes were changed to lipids with transition temperatures close to those of the experiments (18:0 – 18:1 PC and 15:0 PC with transition temperatures of 6 and 35°C, respectively), ensuring the membranes remained permeable to ATP (Figs. S28 and S29). Linear regressions and standard errors were computed using the method of ordinary least squares with the ‘fitlm’ function in Matlab.

## Acknowledgments

We thank Drs. Pierre Goloubinoff, Hagen Hofmann, Axel Mogk, David Scheerer, Ilya Kuprov, and Michal Haran for reading and commenting on the manuscript. We thank Drs. Hagen Hofmann and Tanya Lasitza-Male for the use of their homebuilt Peltier control unit, and Dr. David Scheerer for the fluorescently labeled enzyme adenylate kinase.

## Author Contributions

RC, IR, and GH conceptualized and designed the experiments. RC conducted the experiments. RC, DL, IR, and BY worked on protein expression, purification, and labeling. RC and GH analyzed the results and wrote the manuscript. All authors contributed to editing and discussions.

## Data and materials availability

A summary of the data in this manuscript is provided in the Supplementary Materials. Additional data supporting the findings of this manuscript are available from the corresponding author upon request.

## Supplementary Materials

Supplementary Materials are available for this paper; additional methods section, supplementary Figures S1–S32, and Tables S1–S14.

## Supporting Methods

### Fluorescence polarization experiments

The motional freedom of the encapsulated proteins was assessed using fluorescence polarization experiments (1). Vesicles were loaded with either ClpB molecules fluorescently labeled with Cy3B or κ-casein labeled with CF660R and immobilized on a glass-supported lipid-bilayer surface (see methods section). The dyes were excited with circularly polarized light to avoid photo selection, and the fluorescence was passed through a polarizing cube to split the beam into a horizontal (*I*_H_) and vertical (*I*_V_) component. The fluorescence polarization (P = (*I*_V_ – *I*_H_)/(*I*_V_ + *I*_H_)) was calculated for each 20 ms bin until photobleaching occurred, then averaged on a per molecule basis. The resulting polarization histograms in Figure S3 (blue) exhibit a very narrow distribution of polarization values centered about zero, demonstrating the ability of ClpB and casein to tumble rapidly within the liposome. For comparison purposes, ClpB and casein were non-specifically bound to a glass surface without passivation, resulting in significantly broader polarization distributions (Figure S3 grey, 3 – 5 times greater variance), demonstrating the reduction in tumbling freedom of an immobilized protein.

### Liposome permeability

Liposomes constituting lipids near their liquid-to-gel transition temperature are known to remain permeable to small molecules, such as nucleotides, while retaining larger molecules, such as DNA and proteins (2). The permeability of DMPC liposomes to ATP at the phase transition temperature of 22.5 °C was verified using surface-immobilized liposomes containing a fluorescently labeled adenylate kinase variant (AK 82V, mutations: 82V, 73C, 142C, dyes: AF488, AF594) inside a flow cell. AK 82V displays a large conformational shift from a primarily low FRET efficiency state, in the absence of ATP, to a high FRET efficiency state upon the addition of ATP (3), making it an ideal sensor for the presence of ATP. After encapsulation and immobilization without ATP (50 mM TRIS, 100 mM KCl, 15 mM MgCl_2_, pH 8), trajectories of liposomes containing a double-labeled AK were monitored until photobleaching occurred using a circularly polarized 485 nm laser (ca. 200 nW). The histogram of apparent FRET efficiency values revealed a major population of AK in the low FRET efficiency state (E ∼ 0.4, Figure S4a). The flow cell was then flushed with a buffer solution containing 2 mM ATP, resulting in a shift in the FRET efficiency to 0.68 (Figure S4b) and demonstrating that ATP readily penetrated the liposomes. The reversibility of the process was then demonstrated by flushing with a buffer solution without ATP (Figure S4c), resulting in a shift back to low FRET efficiency. This process could then be repeated with the same outcome (Figure S4d).

### Calculation of the expected FRET efficiencies

The expected FRET efficiencies (*ε*_calc_) were calculated as a function of translocation depth through the lumen of ClpB using a hexameric model of the protein based on PDB 1QVR (4). The translocation depth was measured as the distance from the ‘top’ of the NTD (i.e., at a depth of 0 nm) to the ‘bottom’ of the NBD2, corresponding to a depth of 12.63 nm. *ε*_calc_ values were determined for the line passing directly through the center of the lumen, as well as the limiting values based on the boundaries of the pore. The distance *d* separating the reference labeling location on ClpB and the interior of the lumen was measured in the following manner. First, the structure of ClpB was oriented such that the *z*-axis passed directly through the center of the lumen. Next, the structure of ClpB was divided into 150 even slices orthogonal to the *z*-axis. For each slice, the distances from the reference labeling location on ClpB to the center of the lumen and the atoms lining the interior of the pore were determined. The distances corresponding to the atoms closest and furthest from the reference point in each slice were taken as the limiting boundaries of the pore. *ε*_calc_ values were determined by inputting the measured distances into

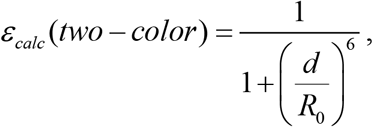

where *R*_0_ is the Förster radius provided in the main text. The total *ε*_calc_ value in the three-color map was determined as

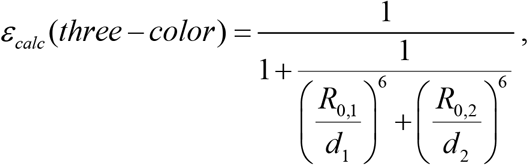

where the subscripts 1 and 2 refer to the two acceptors (5).

### Calculation of the expected number of proteins inside a liposome

Liposomes were prepared by the reconstitution of dried lipids using a solution containing a mixture of ClpB and casein. Assuming the proteins are randomly distributed throughout the solution, the number of proteins contained inside an individual liposome obeys a Poisson distribution (*P*(*i,λ*) = *λ*^*i*^e^−*λ*^/*i*!), where *i* is the number of proteins in a given liposome and *λ* is the average number of proteins contained per liposome. The expectation value, λ, is readily calculated using the concentration of proteins in solution (*c*_P_) and the solution volume contained inside a liposome (*V*_L_) with the relation λ = *c*_P_×*V*_L_. The diameter of the liposomes (= 119 nm, see Figure S1) was used to estimate the interior volume of the liposomes. Assuming a spherical shape, the volume was calculated using the radius of the liposome minus the thickness of the lipid bilayer (= 3.7 nm at 30 °C for DMPC (6)). An example calculation for the expected number of caseins per liposome, prepared from an initial solution of 3 μM casein, is provided below.

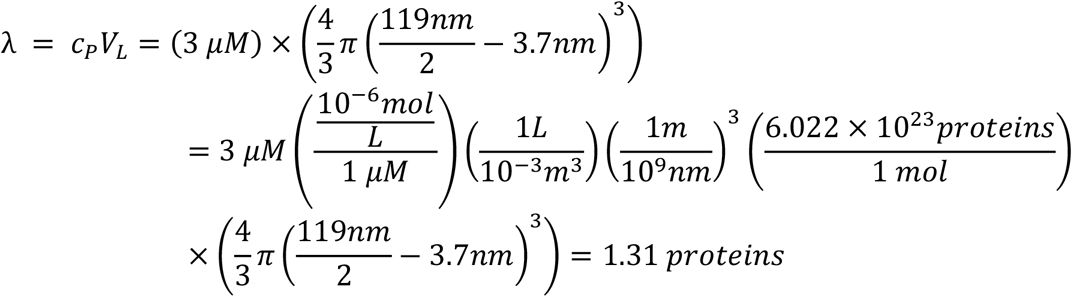

Therefore, the 3 μM solution is expected to produce liposomes that contain an average of 1.3 caseins. Following a similar procedure, liposomes prepared using a solution of 200 nM ClpB are expected to have a *λ* of only 0.09, that is, less than 1 in 10 liposomes contain ClpB. While the above method is of course approximate, liposomes prepared from a solution containing both ClpB and casein are expected, on average, to be sparsely populated with ClpB and contain only a single casein. We note that our smFRET experiments focus only on liposomes containing ClpB, and therefore, the excess of liposomes that do not contain ClpB are not monitored in our experiments.

### Steady-state fluorescence anisotropy

Steady-state fluorescence anisotropy measurements were performed on a Fluorolog-3 spectrofluorometer (Horiba) equipped with automated polarizers. Solutions of the fluorescently labeled proteins (ca. 200 nM) in HEPES buffer were measured in triplicate using a 5 s integration time. Samples containing Cy3B were excited at 550 nm and the emission was recorded at 570 nm. Samples containing CF680R were excited at 670 nm and the emission was recorded at 690 nm. Both the excitation and emission monochromators were set to slit widths of 5 nm. Anisotropy values were calculated using the built-in FluorEssence software. The fluorescence anisotropy values of Cy3B labeled casein in the presence of increasing ClpB concentration are reported in Figure S2. The fluorescence anisotropy of fluorescently labeled ClpB is provided in Table S10. The alignment of the polarizers was verified by confirming that the anisotropy of light scattered by a dilute Ludox suspension (= 0.9887 ± 0.0002) was near unity.

### Fluorescence anisotropy decay experiments

Solutions of the fluorescently labeled proteins (ca. 200 nM) in HEPES buffer were excited (510 nm pulsed laser diode) using vertically polarized light and monitored (570 nm, 16 nm slit width) along the vertical (*I*_VV_) and horizontal (*I*_VH_) axes on a FluoroHub time-correlated single-photon counting instrument (Horiba). Photons were collected in 4096 channels with a time-per-channel of 13.65 ps. Following a previously reported procedure (7), the fluorescence anisotropy was quantified by globally analyzing *I*_VV_ and *I*_VH_ according to Equations S1 – S2 using an in-house Matlab script.

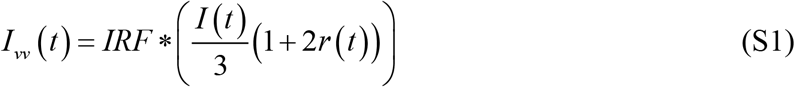

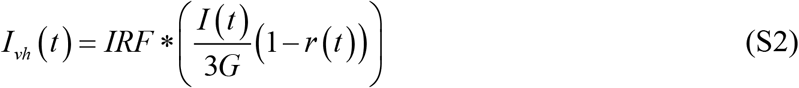

In Equations S1 and S2, IRF is the instrument response function, which was determined using a Ludox solution, ^*^ denotes the discrete convolution of the *IRF* with the model, the G-factor (*G*) is the deviation of the detectors from uniform detection of the two polarization, which was determined using Horiba software through integration of the *I*_HV_ and *I*_HH_ channels, *I*(*t*) is the fluorescence decay of the dye in the absence of anisotropy, and *r*(t) is the time-dependent anisotropy given in Equation S3. Equation S3 assumes the fluorophore bound to the protein experiences segmental motion with a fast-tumbling time *ϕ*_F_, while the protein experiences a slower tumbling time *ϕ*_P_ (8). The preexponential factor α is related to the extent of the hindrance of the fluorophore motion, and *r*_0_ (= 0.40) is the limiting anisotropy at time zero.

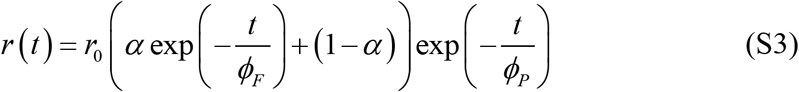

The dye Cy3B required a single fluorescence lifetime (*τ*_D_), as shown in Equation S4.

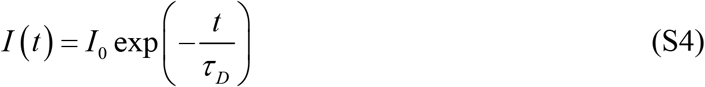

The parameters were optimized with the trust-region-reflective algorithm (9), and the uncertainties were estimated using nonlinear regression estimators. The goodness-of-fit was quantified using the normalized χ^2^ term, where unity reflects a perfect fit with Poisson noise. A list of all retrieved parameters is given in Tables S10 and S11. An example fit of a fluorescence decay is given in Figure S30. The steady-state fluorescence anisotropy (*r*_SS,calc_) was estimated from the retrieved parameters according to Equation S5.

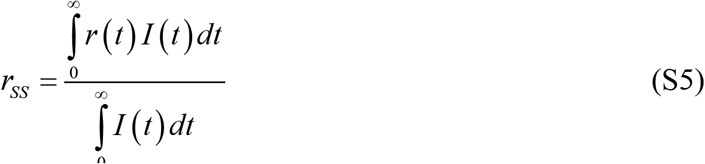

### Distribution of labeled casein inside liposomes

Proper interpretation of the smFRET results requires knowledge of the number of dyes covalently bound to casein. In the three-color experiments, casein was labeled with the donor, allowing us to directly observe the number of dyes inside the liposome by counting the number of donor photobleaching steps and selecting only liposomes that contained a single step. On the other hand, casein was labeled with the acceptor CF660R in the two-color experiments, making the number of photobleaching steps difficult to determine using continuous wave donor excitation. To minimize the probability of double-labeling casein, labeling was carried out with a sub-stoichiometric amount of dye of ca. 0.6 eq. Furthermore, to demonstrate that the majority of casein molecules were single labeled, the labeled casein was loaded into liposomes at the relatively low concentration of 1 μM (with respect to the dye), corresponding to an average of 0.44 (= [dye]×liposome volume) dyes per liposome. As the vesicles were formed, the casein molecules distributed themselves according to a Poisson distribution, with an expectation value equal to the average number of molecules per liposome. If casein contained a significant fraction of double-labeled molecules, the dye would not obey the same distribution, as there would be a significant increase in the occurrence of two or more acceptors per liposome. Figure S6 demonstrates that the distribution of CF660R dyes obeys the Poisson distribution, as expected for single-labeled casein.

### Derivation of leak and background corrections

While the correction of channel crosstalk is relatively straightforward for two-color FRET experiments, we derived general matrix-based equations for the crosstalk and background correction for an arbitrary number of fluorescent species. For the sake of simplicity, derivation is shown for the case of three dyes, as is the case for our three-color experiments. The final equation used is Equation S11, which corrects all channels in a single operation.

Let *O*_*i*_ be the total observed fluorescence intensity in channel *i*, given by the fluorescence of each species in the channel, plus a background term.

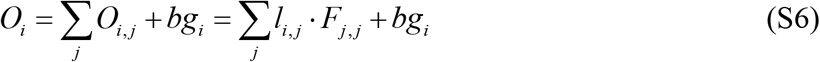

Where *O*_*i, j*_ is the observed fluorescence of species *j* in channel *i, bg*_*i*_ is the background in channel *i, l*_*i, j*_ is the leak factor of species *j* in channel *i* (*i*_j,j_ = *F*_i,j_/*F*_j,j_, and noting that *l*_j,j_ = 1), and *F*_*i, j*_ is the fluorescence intensity of species *j* in channel *i*. In matrix notation, Equation S6 becomes,

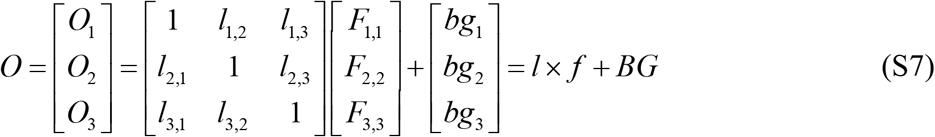

Where O is the matrix of observed fluorescence in each channel, *l* is the matrix of leak values, *f* is the vector of *F*_j,j_ values, and BG is the vector of background values in each channel. The total fluorescence of species *j* is given by the sum of its contributions to each channel.

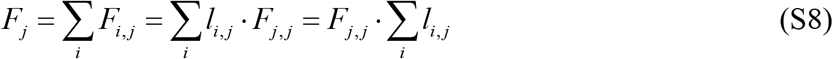

In matrix notation,

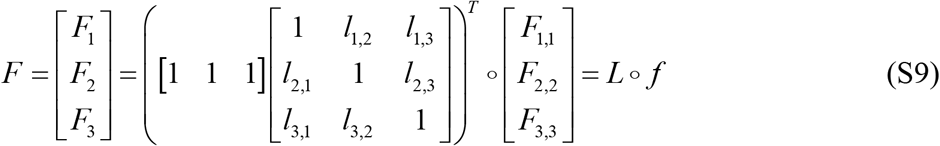

where ○ denotes the element-wise product and *L* is given by,

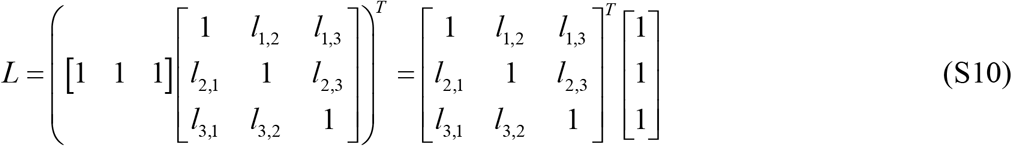

Substituting *f* in (S7) into (S9) yields the corrected fluorescence as S11.

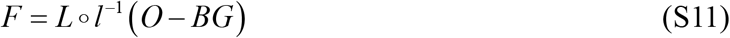

### Leak factor determination

The leak factors *l*_j_ for each species *j* are determined by acquiring fluorescence from single-labeled molecules. The matrix *l* is decomposed into separate elements for each species *j*.

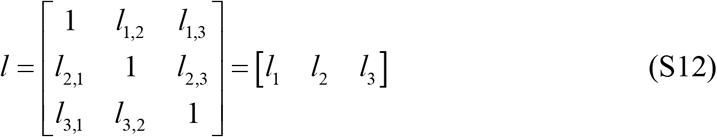

Rearranging Equation (S7), we know,

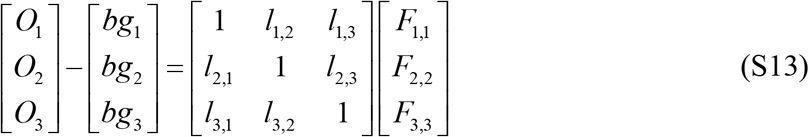

Given that for a single-labeled molecule, the only contribution to each channel arises from the fluorescence of species *j*, the corresponding *l*_j_ element of the leak matrix can be found using Equation S14.

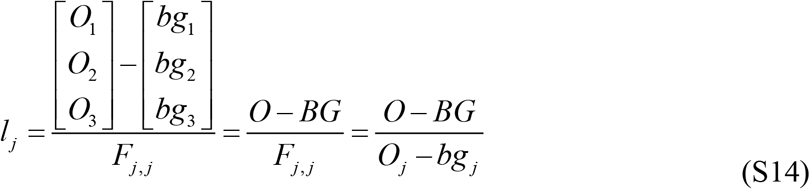

A plot of the molecule-averaged leak factors for Cy3B in the two-color experiments is given in Figure S31. The emission of CF660R was sufficiently separated that its emission in the Cy3B channel was negligible (i.e., leak factor = 0). The corresponding leak factor matrix is given by 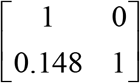. Figure S32 plots the leak factors for Cy3B, AF647, and CF680R in the 3-color experiments. The corresponding leak factor matrix is given by 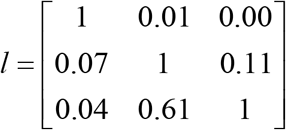.

### Determining inter-acceptor energy transfer

The presence of FRET between the two acceptors (with a separation of 6.8 nm) in the three-color experiments can potentially affect the interpretation of the results. The fluorescence of double-labeled ClpB molecules was collected after acceptor excitation to assess the extent of inter-acceptor FRET. After leak correction, the presence of inter-acceptor energy transfer was assessed by comparing the photon flux of acceptor 2 (CF680R, the more red-shifted dye) before and after photobleaching of acceptor 1 (AF647). When energy transfer occurs between the dyes (from A1 to A2), the fluorescence intensity of A2 would be larger before the photobleaching of A1. However, as seen in Figure S18, the lack of such an effect demonstrates the absence of energy transfer between acceptors.

### Fluorescence correlation spectroscopy

Data was collected using continuous wave excitation (560 nm, 30 μW). The fluorescence emission was split with a 50/50 beam-splitting cube and recorded on two detectors to avoid after-pulsing effects. FCS curves were generated using the PicoQuant SymPhoTime 64 software (Figure S27). The curves were fitted to Equation S15 below using a weighted least-squared function, where the weight of each point was determined by its standard deviation. In this equation, *ω* is the aspect ratio of the confocal volume, *N* is the average number of molecules in the confocal volume, *τ*_D_ is the diffusion time of a molecule through the confocal volume, *τ*_T_ is the triplet lifetime, *f*_T_ is the fraction of triplet state, and *G*_∞_ is the autocorrelation at a very long time. The retrieved parameters are provided in Table S12.

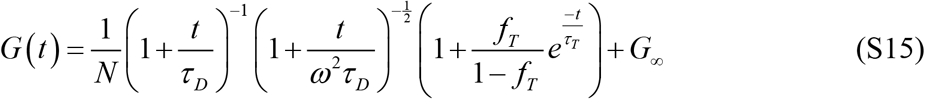

Using the known diffusion coefficient of Cy3B (*D*_0_ = 334 μm^2^/s at *T*_0_ = 24 °C) (10, 11) and a confocal volume (*V*_eff_) of 0.286 fL (calculated using the known *D*_0_), the temperature *T* in the confocal volume was calculated using Equation S16, where the viscosity (*η*) was corrected using the Vogel Relation given in Equation S17. In Equation S17, temperature (*T*) is in K, and the empirical constants a, b, and c are equal to –4.5318, –220.57, and 149.39, respectively (12).

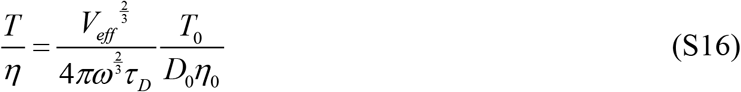

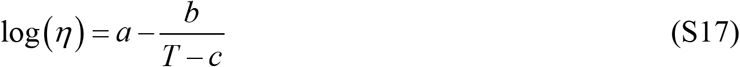

Equation S16 is derived starting from the relationship between effective confocal volume (*V*_eff_) and *D* given in Equation S18 (13). The change in diffusion coefficient with temperature (Equation S19) is determined using the Stokes-Einstein equation for diffusion. *D* in Equation S18 was substituted into S19, which, after rearrangement, yielded S16.

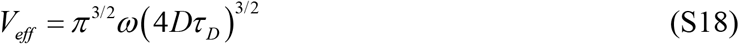

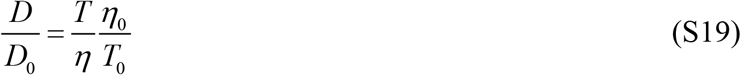

## Supporting Figures

**Figure S1.**
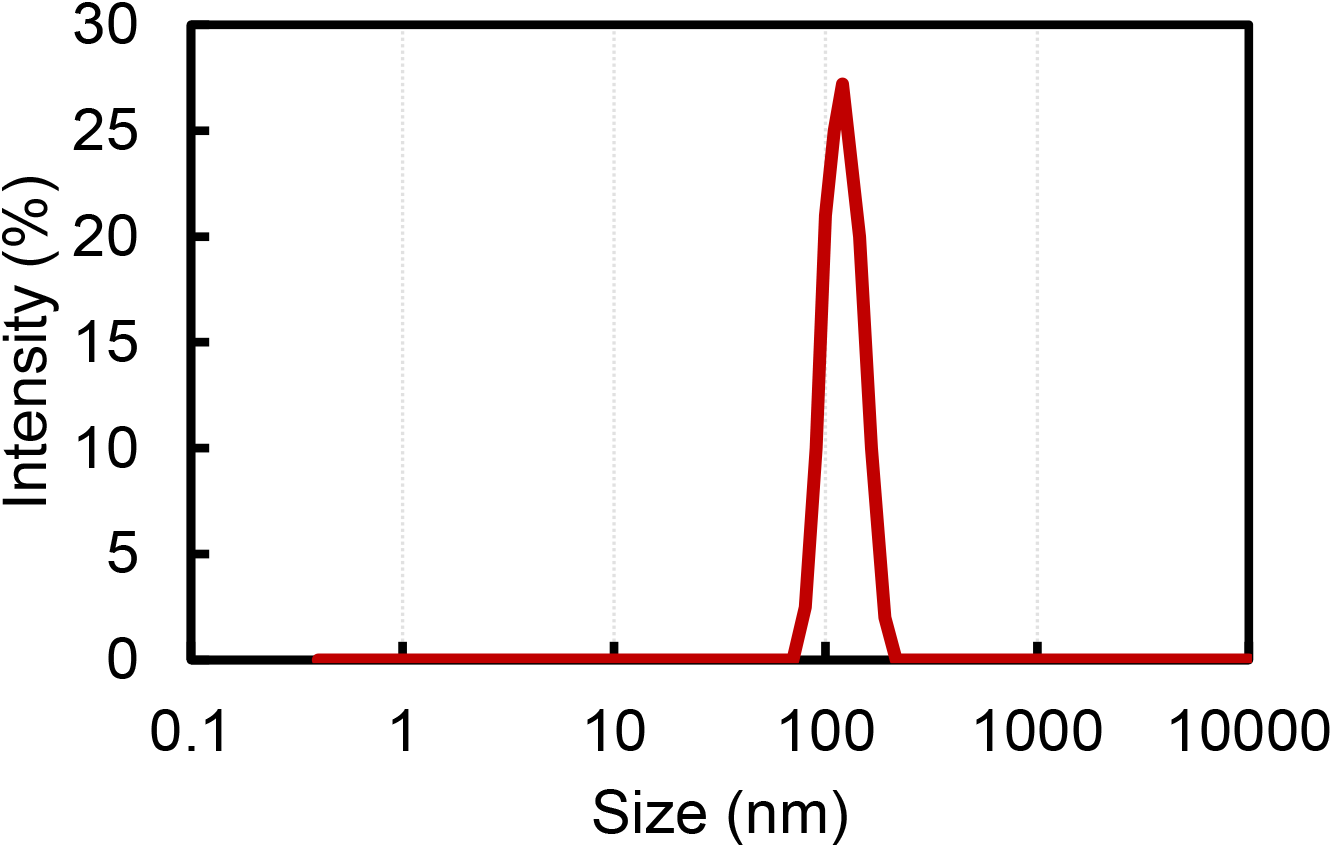
Extrusion produces monodisperse large unilamellar vesicles. Sample distribution of the sizes of 99:1 DMPC: Biotinyl PE liposomes prepared by extrusion through a 0.1 μm membrane, as measured by dynamic light scattering (DLS, Malvern Zetasizer). Extrusion produced vesicles of a unimodal distribution with a *z*-average size of 119 ± 1 nm and a dispersity of 0.05 ± 0.01. Measurements were repeated in triplicates, see Table S9.

**Figure S2.**
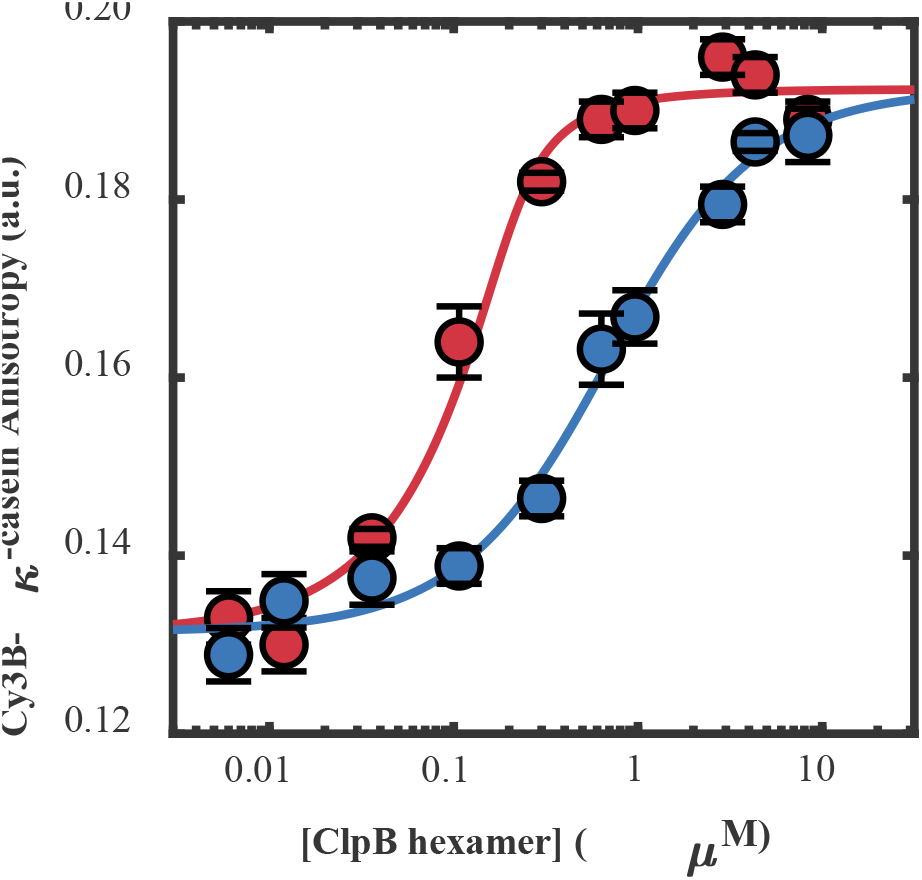
ATPγS increases the apparent affinity of casein to ClpB. Steady-state anisotropy of Cy3B labeled κ-casein (200 nM) in the presence of increasing concentrations of WT ClpB, as well as (blue) 2 mM ATP or (red) 2 mM ATPγS. All measurements were repeated in triplicates. The anisotropy values were fitted (lines) with a bimolecular binding equilibrium curve, yielding apparent dissociation constants of 600 ± 100 nM and 20 ± 10 nM in the presence of ATP and ATPγS, respectively. The apparent constants are affected by the different times of interaction of casein with ClpB in the presence of ATP or ATPγS; smFRET experiments found a similar 34-fold increase in dwell time in the presence of ATPγS.

**Figure S3.**
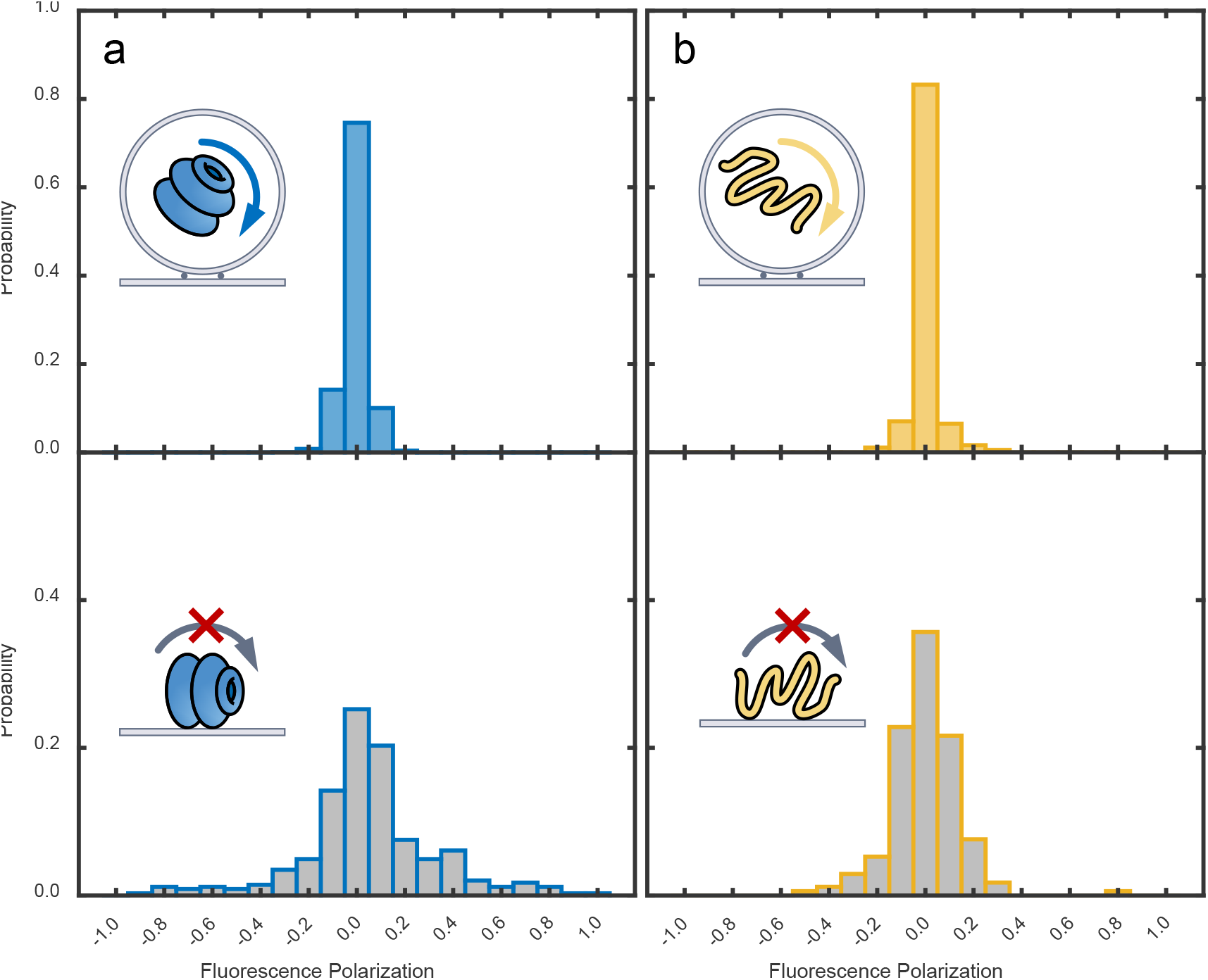
ClpB and κ-casein freely diffuse inside immobilized liposomes. Distributions of the fluorescence polarization (P = (*I*_V_–*I*_H_)/(*I*_V_+*I*_H_)) of fluorescently labeled (**a**) ClpB (359C, Cy3B) and (**b**) κ-casein (sigma, CF660R) trapped inside surface-immobilized DMPC liposomes (blue) and directly immobilized on a glass surface (grey). The polarization distribution for ClpB (σ = 0.05, *N* = 185) and casein (σ = 0.07, *N* = 153) inside the vesicles is much narrower than for protein molecules non-specifically immobilized on a glass surface (ClpB: σ = 0.14, *N* = 463, casein: σ = 0.35, *N* = 342). The narrow distribution demonstrates that ClpB and casein have rotational freedom and can freely diffuse within the vesicles.

**Figure S4.**
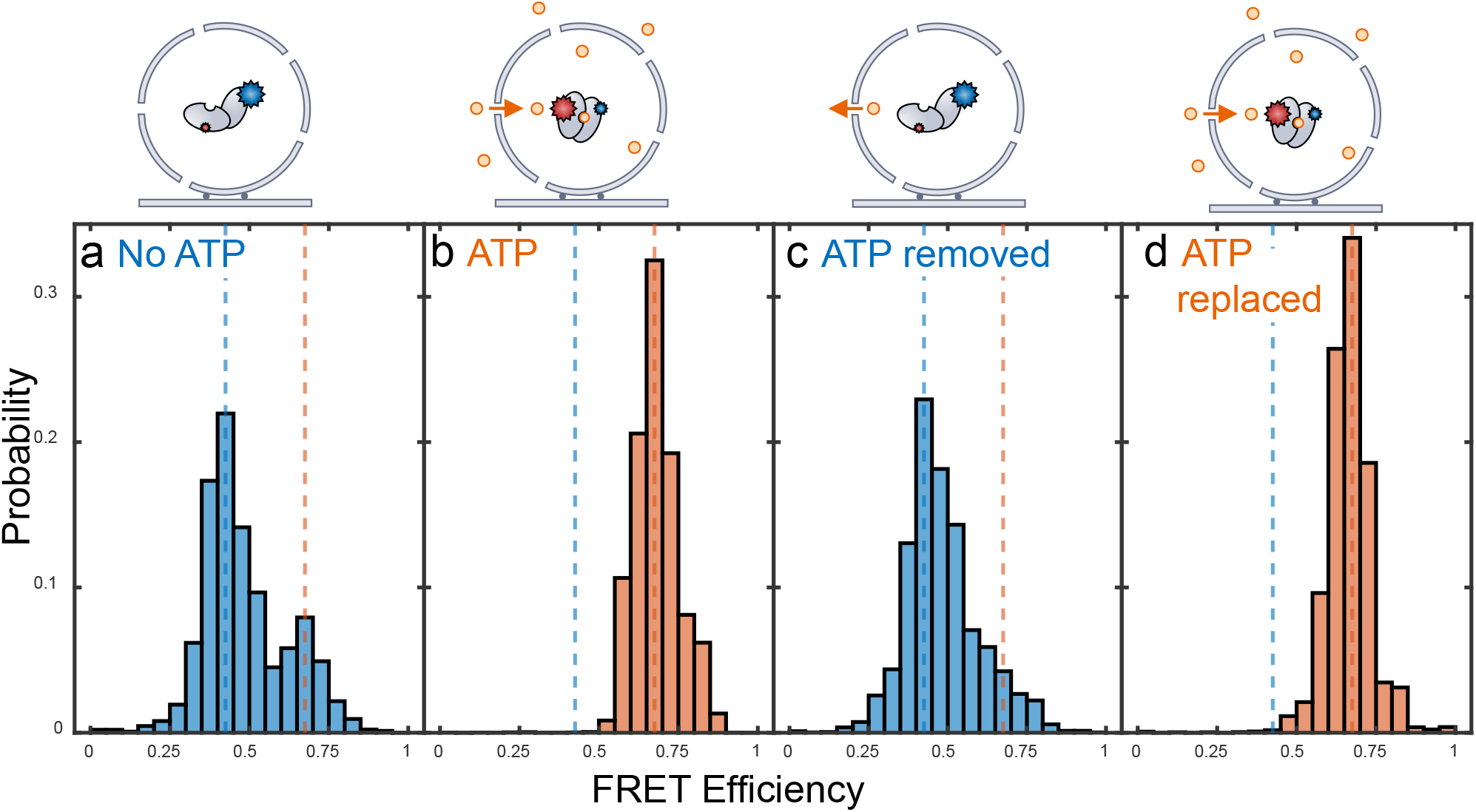
ATP readily diffuses in and out of DMPC liposomes. Histograms of the FRET efficiency values of fluorescently-labeled adenylate kinase variant AK82V (3) encapsulated in surface-immobilized DMPC (14:0 PC) liposomes at 22.5 °C without (**a**,**c**) and with (**b**,**d**) 2 mM ATP. (**a**) The protein was encapsulated without ATP (50 mM TRIS, 100 mM KCl, 15 mM MgCl_2_, pH = 8) and the vesicles were immobilized onto a glass surface inside a flow cell. The histogram (*N* = 116 molecules) displays a major population at low FRET efficiency (ca. 0.4), as expected for apo AK (3). (**b**) Upon the addition of 2 mM ATP, the entire population (*N* = 97) shifted to a high FRET efficiency (ca. 0.68), demonstrating that ATP could permeate through the DMPC membranes and reach the encapsulated protein. (**c**) The reversibility of the process was demonstrated by the recovery of the low FRET population (*N* = 55) when the cell was flushed with buffer without ATP. (**d**) The high FRET population (*N* = 65) reappeared upon the repeated addition of ATP.

**Figure S5.**
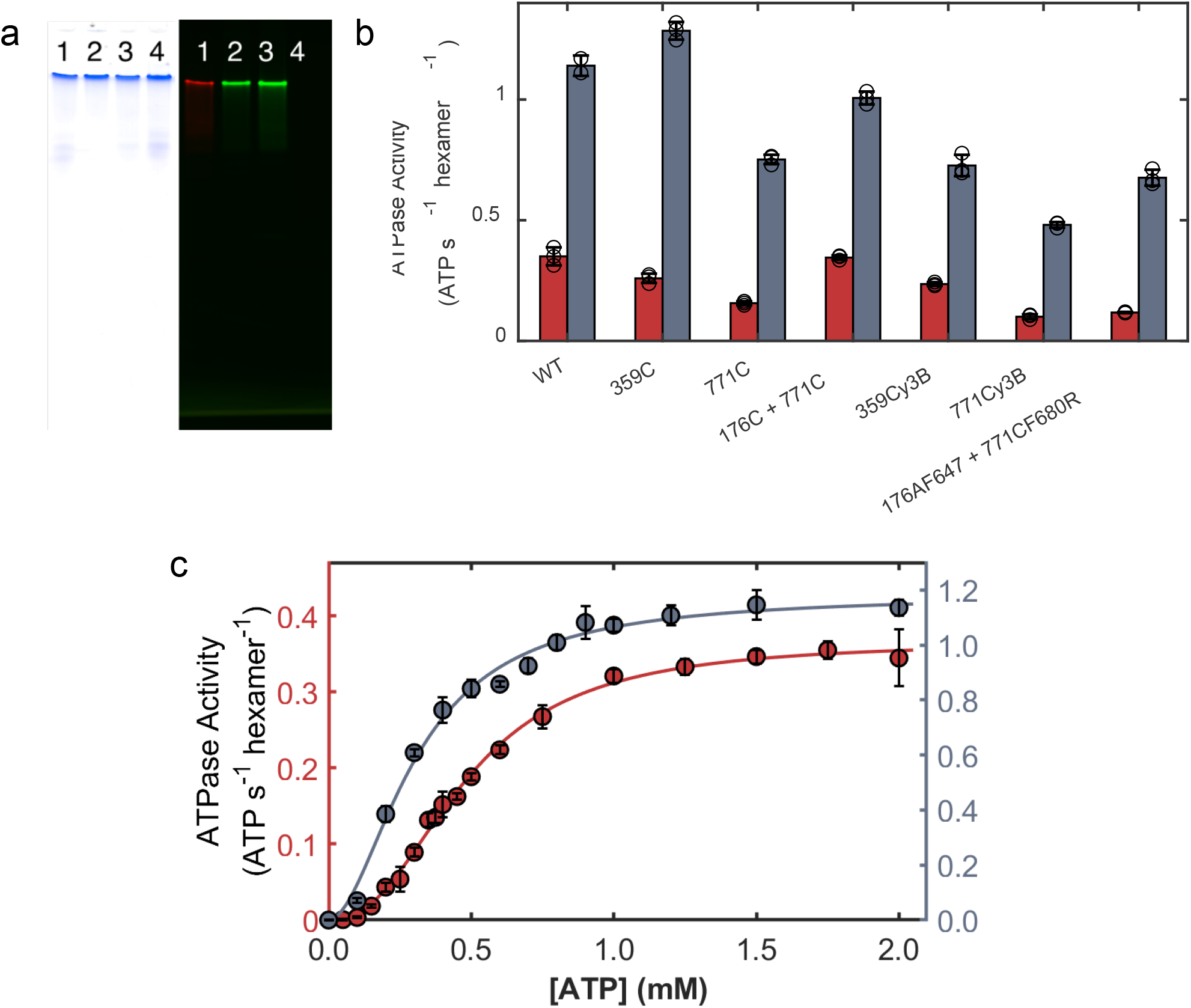
ClpB remains active after point mutations and fluorescence labeling. (**a**) Native gel electrophoresis (4 % acrylamide, with 10 mM MgCl_2_, 2 mM ATP at 70 V for 12 hours (4)) of reassembled fluorescently labeled double cysteine mutant ClpB (176 – AF647 + 771 – CF680R, lane 1), single cysteine mutants labeled with Cy3B (359, lane 2 and 771, lane 3), and wild-type (WT) ClpB (lane 4). Shown are the Coomassie Blue stain (left) and fluorescence (Typhoon scanner, right) following excitation at 635 nm (red) and 532 nm (green). As previously reported (4, 14), the presence of a high molecular weight single band demonstrates the successful hexameric assembly. The red dyes on the double-labeled ClpB exhibited fluorescence under red light, both ClpB variants labeled with Cy3B exhibited fluorescence under green light, while the WT ClpB experienced no fluorescence. (**b**) ATPase activity of ClpB at 22.5 °C in the presence of 2 mM ATP (red) and upon addition of 3 μM κ-casein (grey). The activity is shown for the WT ClpB, single cysteine mutants (359C, 771C), double mutant (176C + 771), and the fully-fluorescently labeled counterparts. (**c**) ATPase activity of WT ClpB as a function of ATP concentration without (red, left axis) and with (grey, right axis) the addition of 3 μM κ-casein. The solid lines represent the fits according to the Hill equation with Hill coefficients of 2.7 ± 0.1 and 1.9 ± 0.2 and *K*_m_ values of 0.42 ± 0.01 mM and 0.31 ± 0.02 mM, respectively. Activity values are similar to literature-reported values (4, 15).

**Figure S6.**
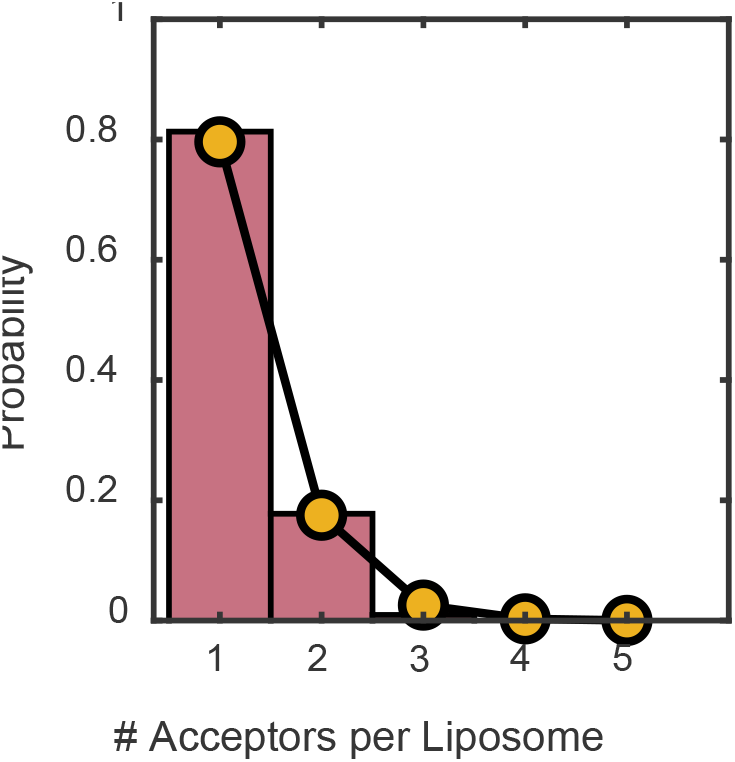
Photobleaching experiments on CF660R labeled casein demonstrate casein overwhelmingly contains at most a single acceptor. Histogram (red) of the number of photobleaching steps for 107 liposomes loaded with 1 μM CF660R-labeled casein, corresponding to an average number of dyes per liposome of 0.44. The orange points represent the expected probability based on the Poisson distribution for single-labeled casein, Pr(# acceptors) = Pr(*i* | *i* > 0) = Pois(i, λ)/(1 – Pois(0, λ), where Pois is the Poisson distribution, λ = 0.44, and *i* (≥ 1) is the number of acceptors. The agreement between the Poisson distribution and the photobleaching experiments demonstrates that the bulk of casein cannot contain more than one acceptor. If casein contained a significant fraction of double-labeled molecules there would be a significant increase in the occurrence of two or more acceptors per liposome.

**Figure S7.**
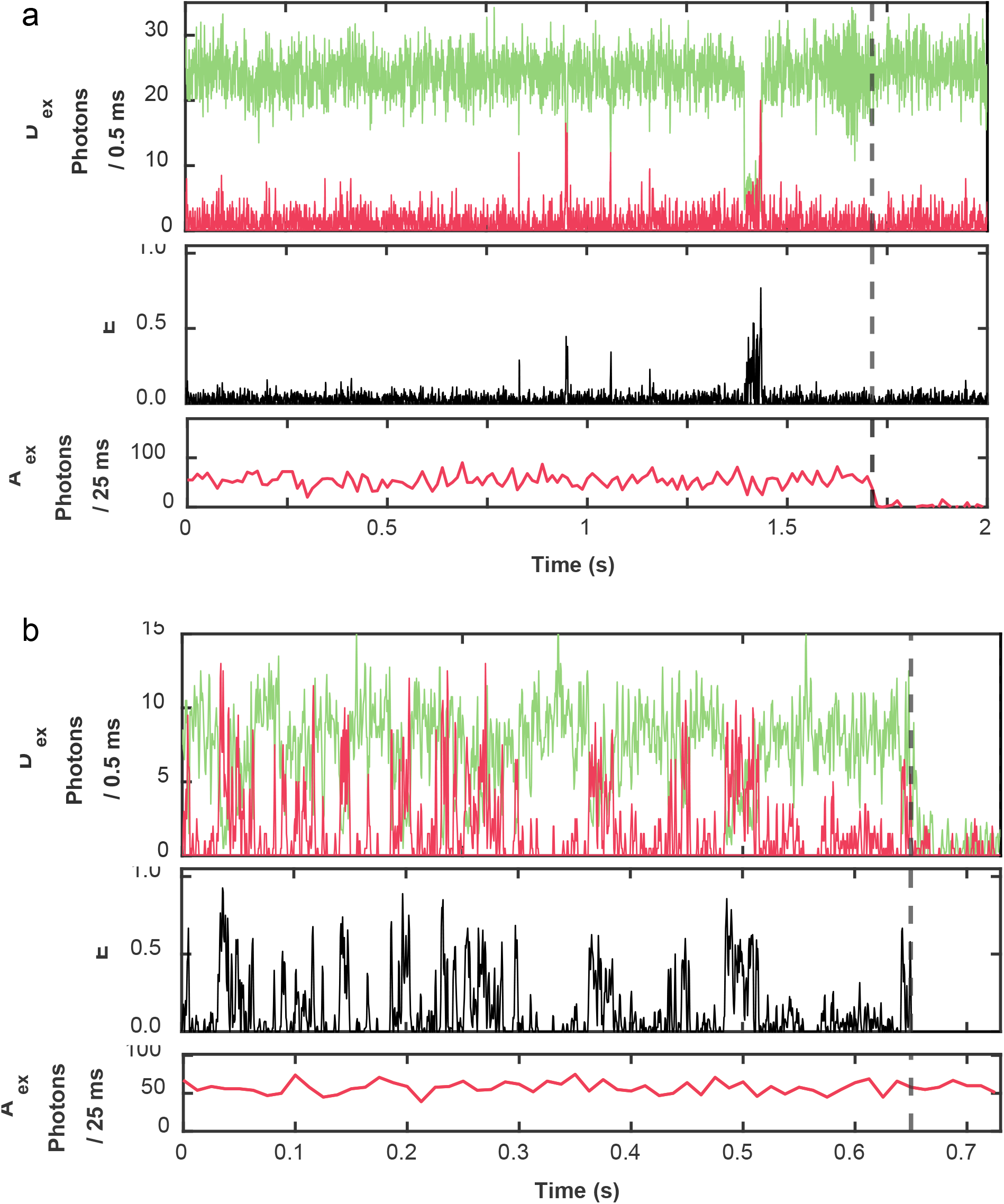
The acceptor remains fluorescent throughout the trajectory. Sample trajectories of translocation of casein through (**a**) NBD1 and (**b**) NBD2 labeled ClpB using pulsed interleaved excitation (500 nW donor, 100 nW acceptor). Shown are the fluorescence after donor excitation (D_ex_, top) of the donor (green) and acceptor (red), the apparent FRET efficiency (middle), and the acceptor fluorescence after direct acceptor excitation (A_ex_, bottom). The dashed line marks the photobleaching of the (**a**) acceptor and (**b**) the donor. The constant acceptor signal (before photobleaching) demonstrates the acceptor remains fluorescent throughout the trajectory.

**Figure S8.**
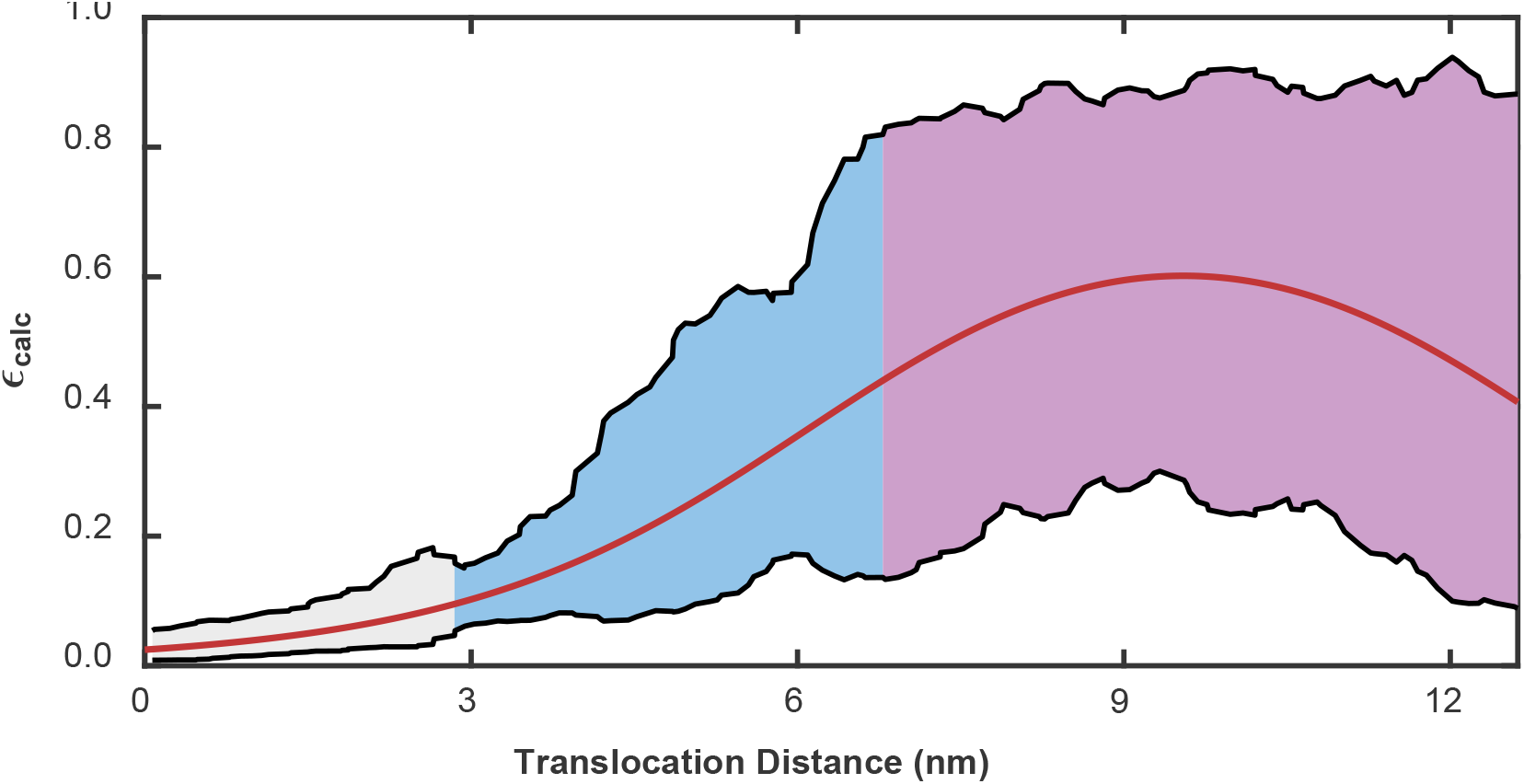
Calculated FRET efficiencies for NBD2 labeled ClpB demonstrate that high FRET efficiency events can only occur if casein passes through the NBD2 ring. A map of the expected FRET efficiency (ε_calc_) for an acceptor as it passes through ClpB with a donor located in the NBD2 (res. 771). The solid red line represents the centerline passing directly through the pore, while the black lines represent the boundaries of the pore. The shaded regions represent the *N*-terminal domain (NTD, grey) and the two NBDs (blue and purple).

**Figure S9.**
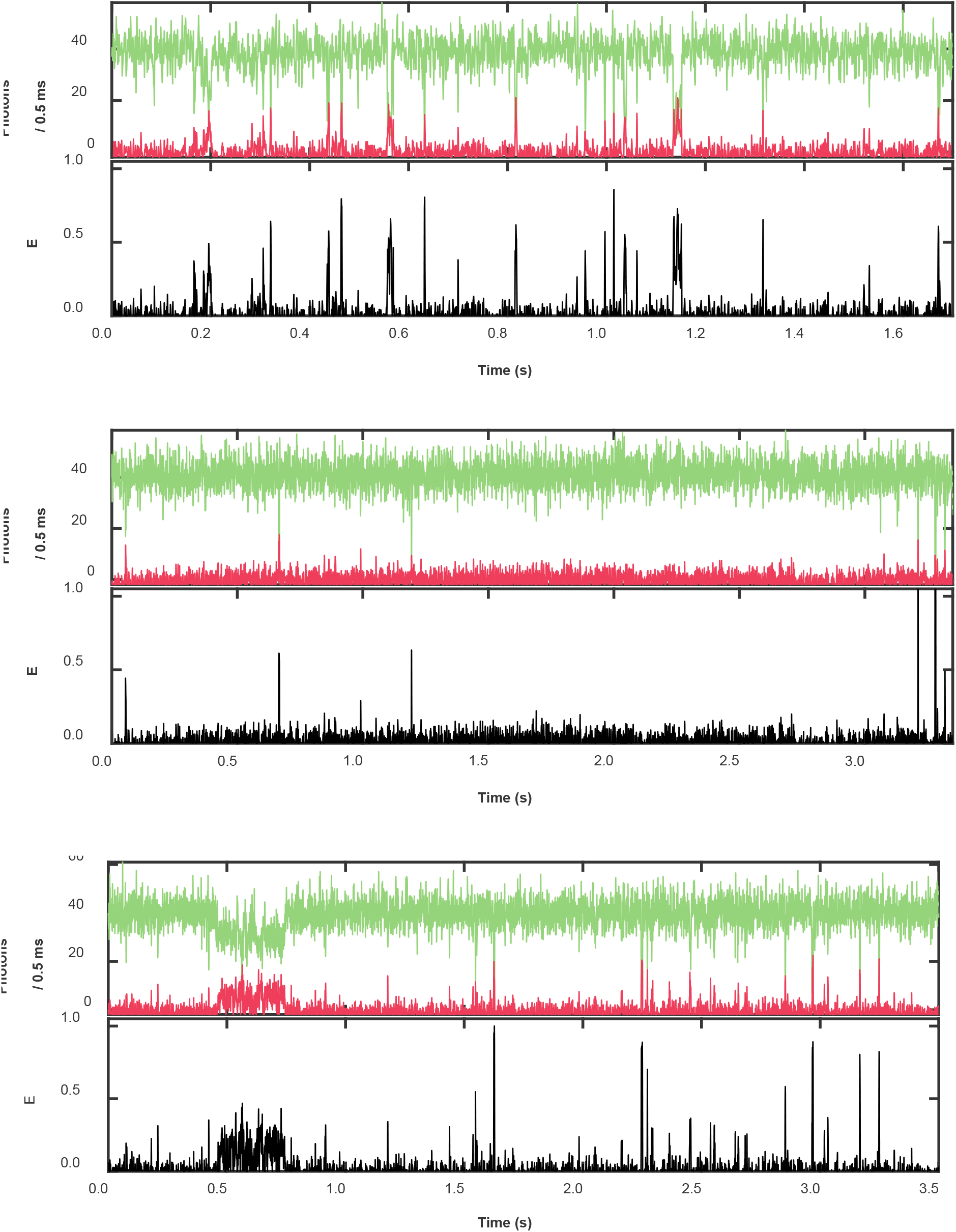
Sample trajectories of translocation of casein through NBD1 labeled ClpB in the presence of 2 mM ATP. Shown are the fluorescence (top panels) of the donor (green) and acceptor (red) and (bottom panels, black) the FRET efficiency.

**Figure S10.**
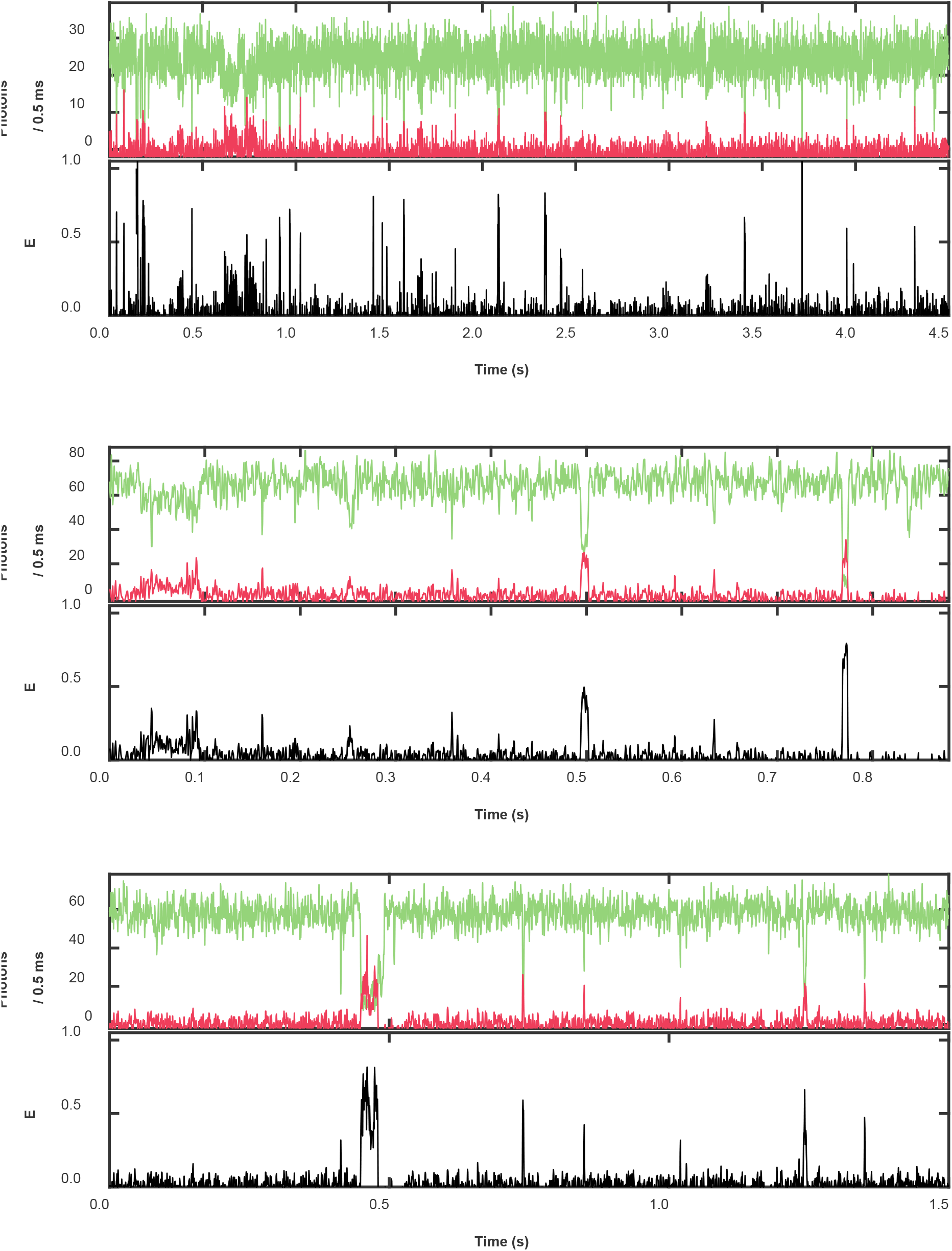
Sample trajectories of translocation of casein through NBD2 labeled ClpB in the presence of 2 mM ATP. Shown are the fluorescence (top panels) of the donor (green) and acceptor (red) and (bottom panels, black) the FRET efficiency.

**Figure S11.**
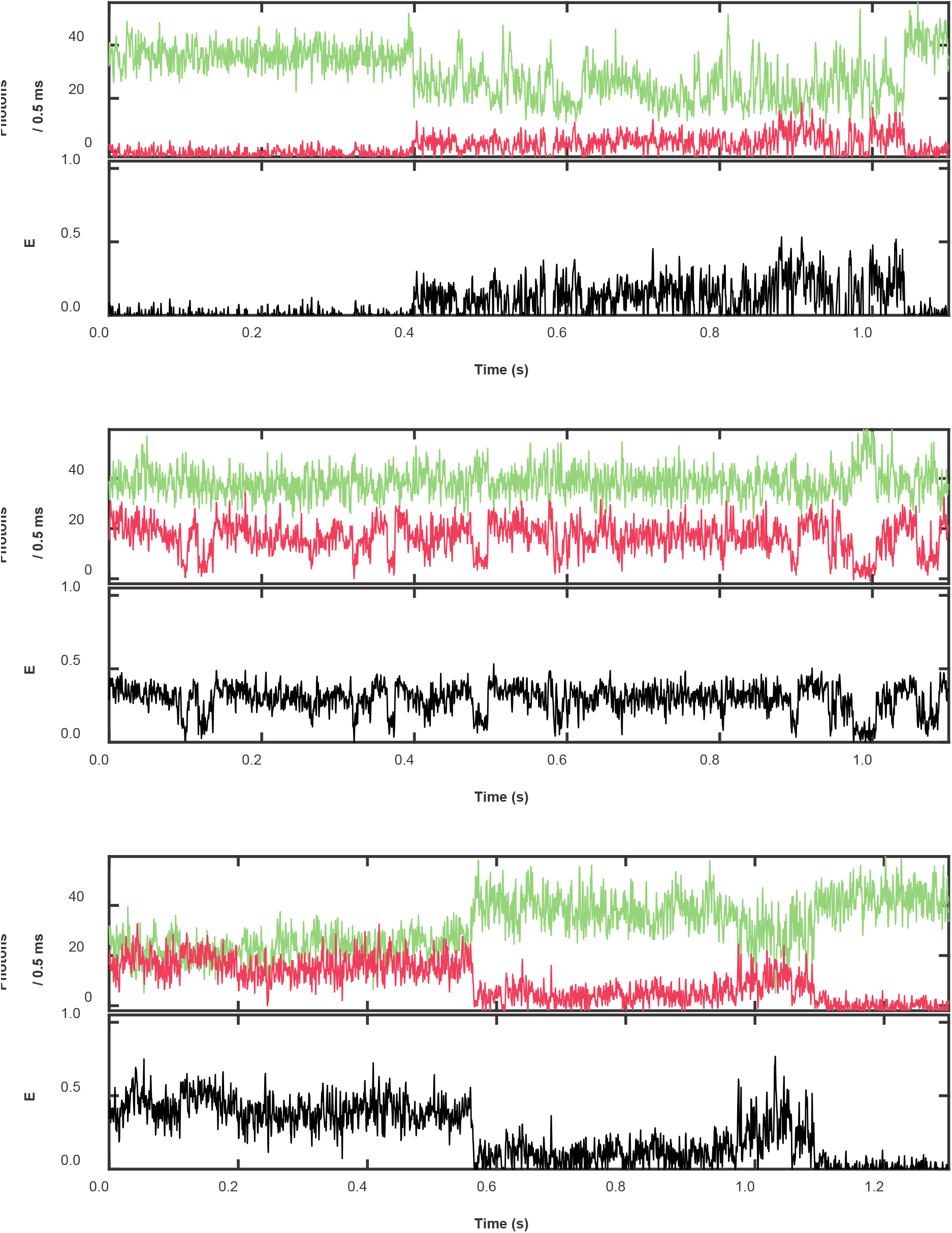
Sample trajectories of translocation of casein through NBD1 labeled ClpB in the presence of 2 mM ATPγS. Shown are the fluorescence (top panels) of the donor (green) and acceptor (red) and (bottom panels, black) the FRET efficiency.

**Figure S12.**
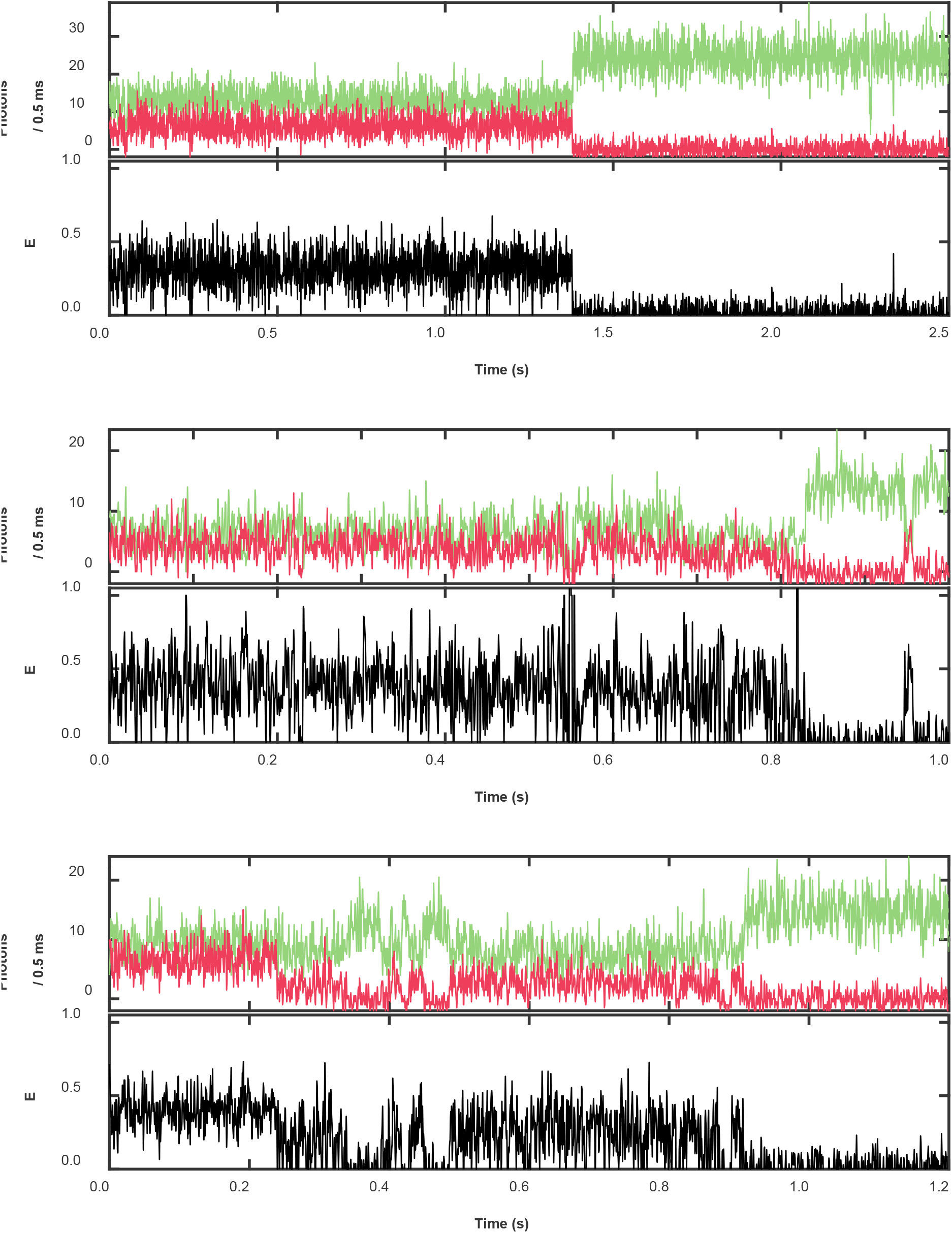
Sample trajectories of translocation of casein through NBD2 labeled ClpB in the presence of 2 mM ATPγS. Shown are the fluorescence (top panels) of the donor (green) and acceptor (red) and (bottom panels, black) the FRET efficiency.

**Figure S13.**
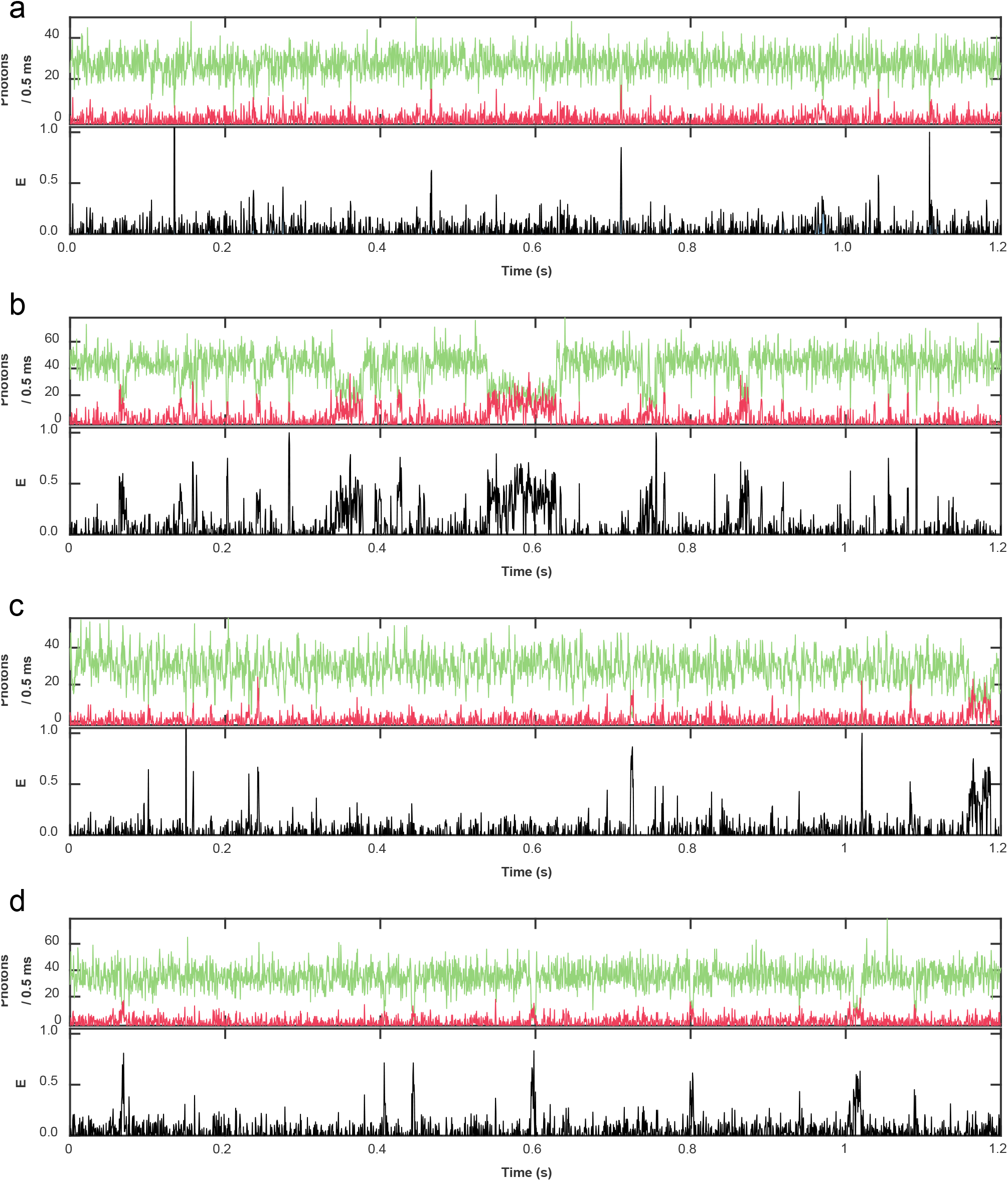
Sample trajectories of translocation of casein as a function of temperature. Shown are the fluorescence (top panels) of the donor (green) and acceptor (red) and (bottom panels, black) the FRET efficiency of translocation events through (**a**,**b**) NBD1 and (**c**,**d**) NBD2 labeled ClpB at (**a**,**c**) 32 and (**b**,**d**) 10 °C.

**Figure S14:**
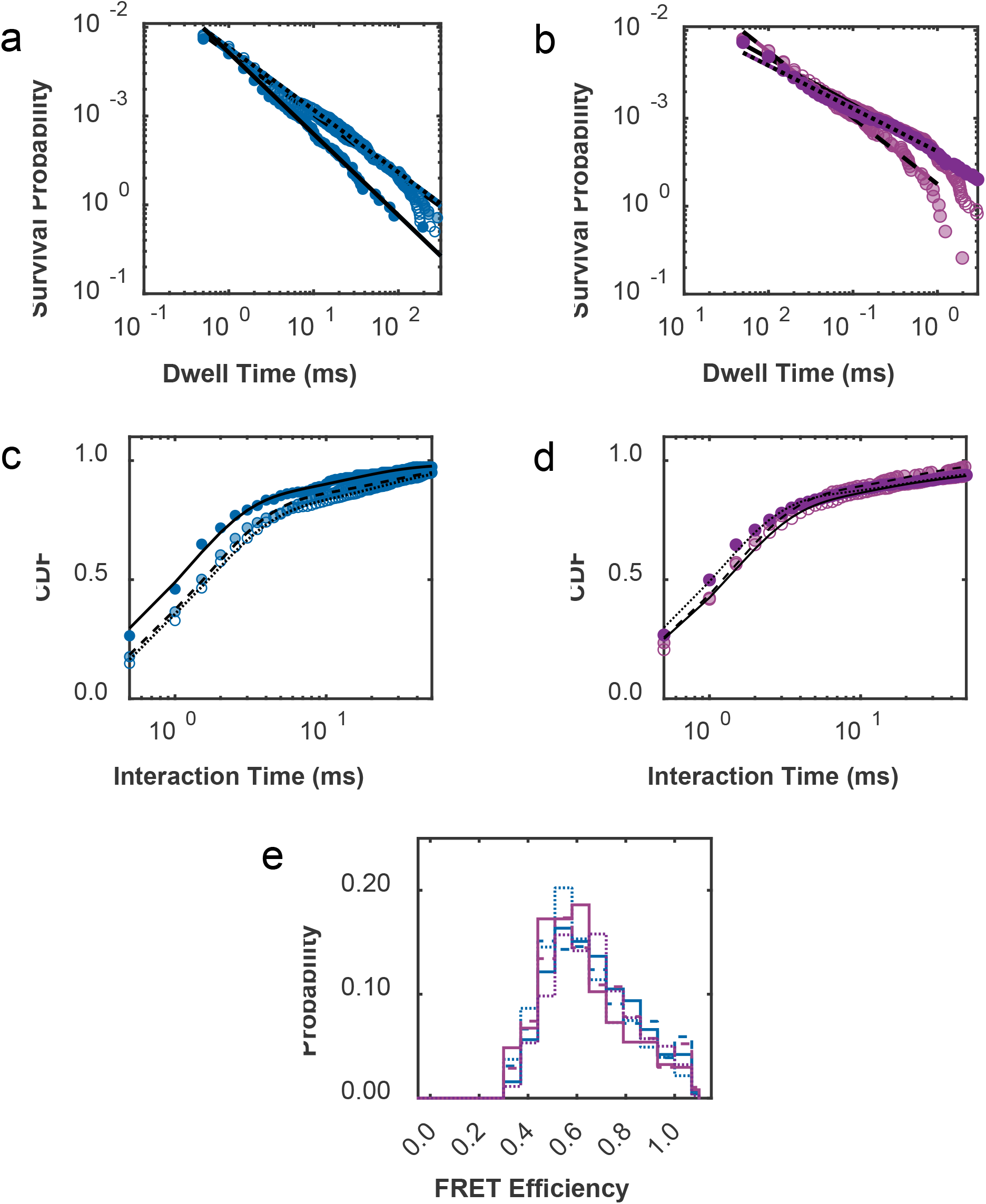
Event dwell time is weakly affected by temperature. (**a**,**b**) Survival probability and (**c**,**d**) cumulative distribution function (CDF) of the dwell time and (**e**) peak FRET efficiency for NBD1 (blue) and NBD2 (purple) labeled ClpB at temperatures of 32 (empty symbols, dotted lines), 22.5 (shaded symbols, dashed lines), and 10 °C (filled symbols, solid lines). The parameters retrieved from the (**a**,**b**) power-law and (**c**,**d**) exponential fits are provided in Tables S4 and S7.

**Figure S15.**
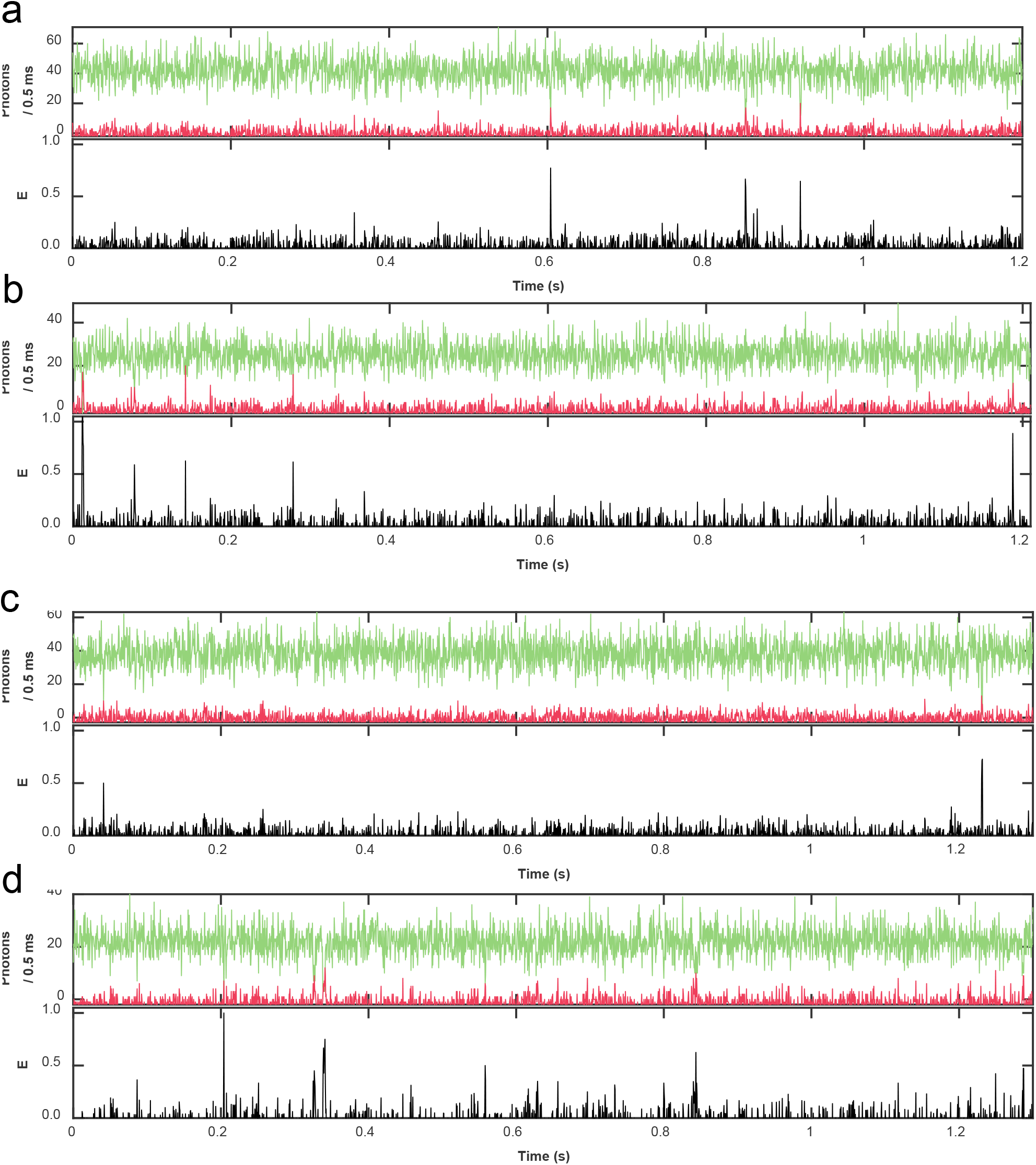
Sample trajectories of translocation of casein as a function of ATP concentration. Shown are the fluorescence (top panels) of the donor (green) and acceptor (red) and (bottom panels, black) the FRET efficiency of translocation events through (**a**,**b**) NBD1 and (**c**,**d**) NBD2 labeled ClpB in the presence of (**a**,**c**) 0.2 and (**b**,**d**) 0.4 mM ATP.

**Figure S16.**
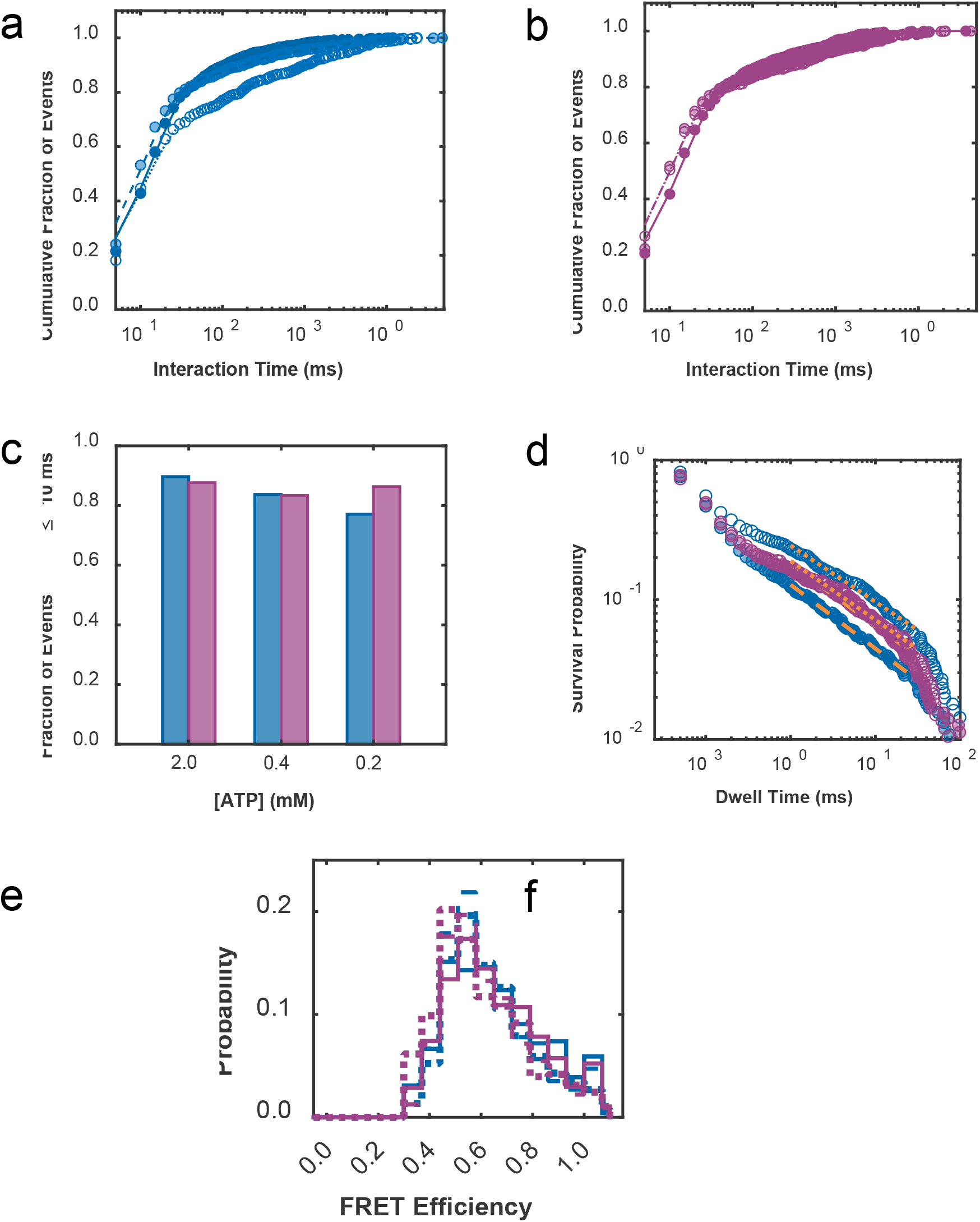
Event dwell time is weakly affected by ATP concentration. (**a**,**b**) Cumulative distribution of event times, (**c**) fraction of short events, (**d**) power-law distribution, and (**e**) distribution of peak FRET efficiencies at ATP concentrations of 2.0 (solid lines, filled symbols), 0.4 (shaded symbols, dashed lines), and 0.2 mM (hollow symbols, dotted lines) for ClpB labeled on NBD1 (blue) and NBD2 (purple). The lines in (**a**,**b**,**d**) represent fits, whose parameters are reported in Tables S4, S5, and S7.

**Figure S17.**
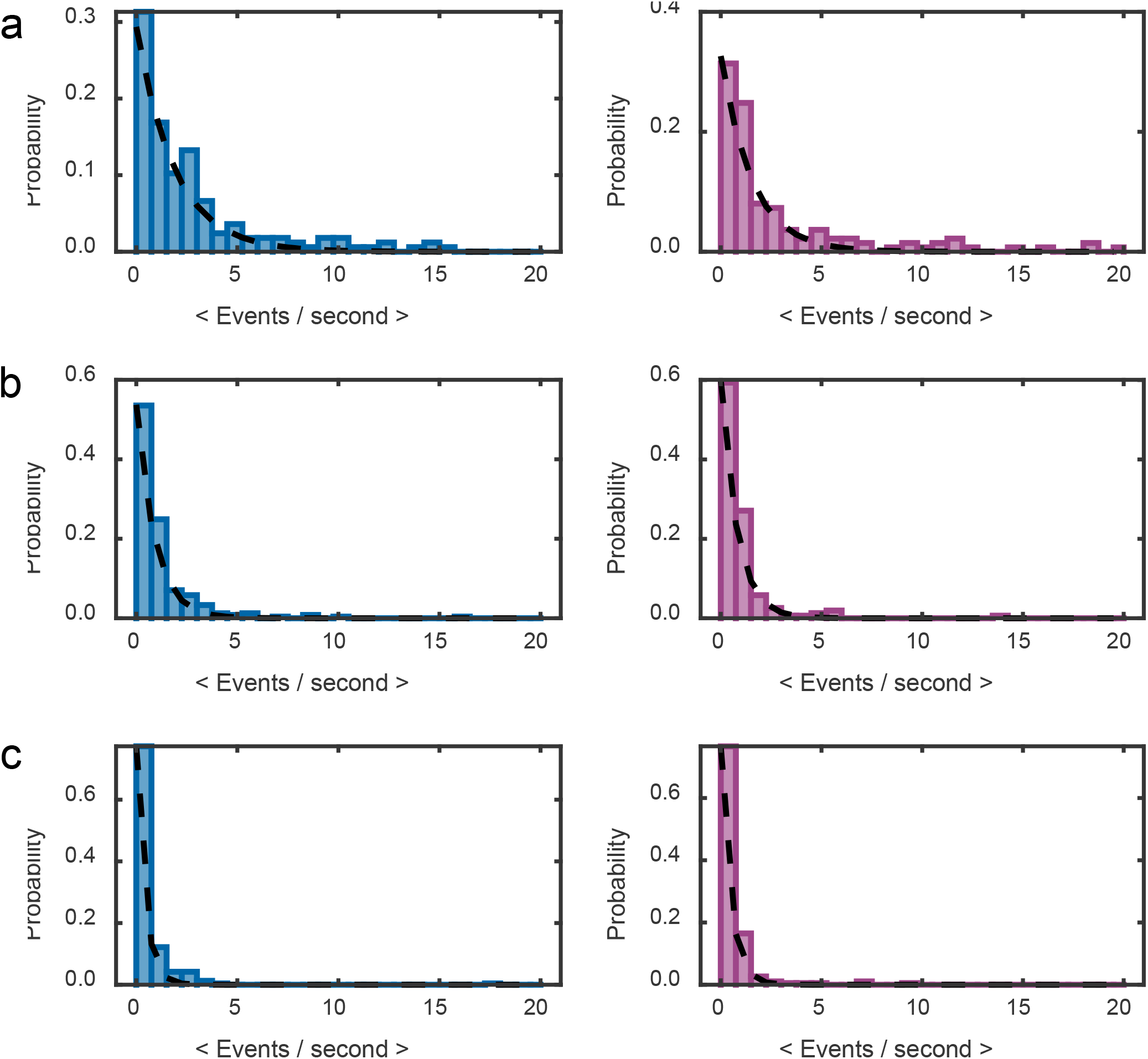
Event frequency directly correlates with ATP concentration. Histograms of the average number of events per second for individual ClpB molecules at ATP concentrations of (**a**) 2.0, (**b**) 0.4, (**c**) 0.2 mM ATP for ClpB labeled on NBD1 (left, blue) and NBD2 (right, purple). The dashed lines represent the exponential fits of the distributions. The parameters retrieved from the fits are reported in Table S8 and in Figure 3b (in the main text).

**Figure S18:**
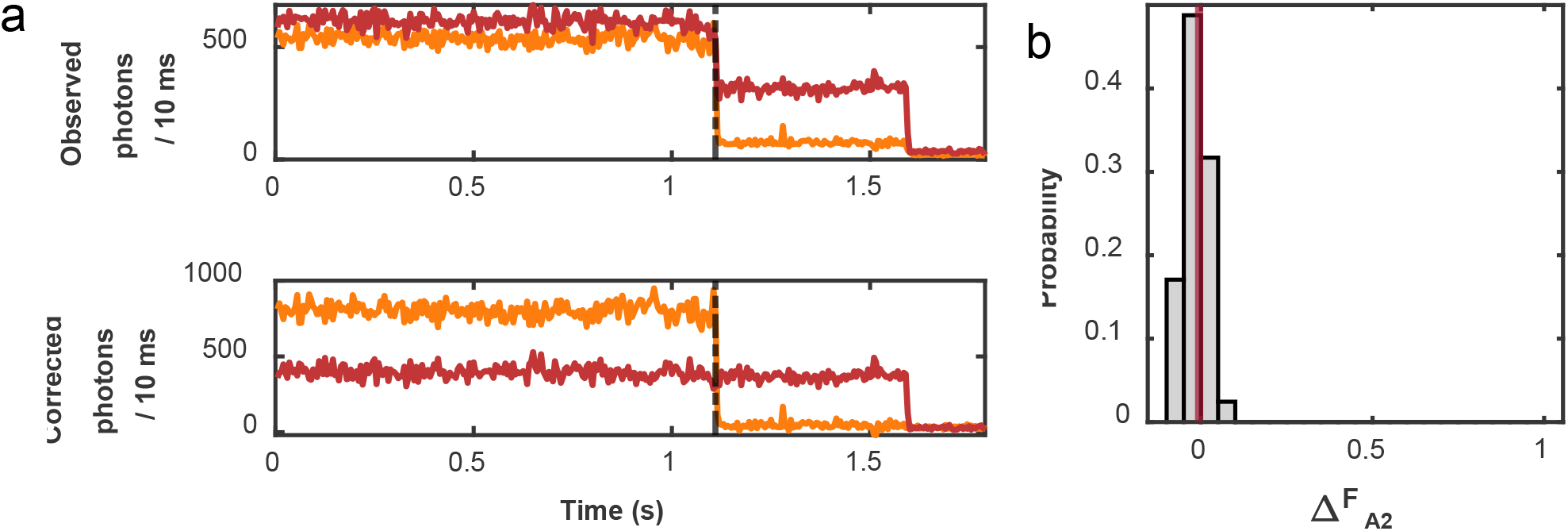
Acceptor photobleaching study on double-acceptor labeled ClpB demonstrates negligible inter-acceptor energy transfer. (**a**) Sample trajectory of a double-labeled ClpB molecule after excitation at 640 nm. Both acceptor 1 (A1, orange) and acceptor 2 (A2, red) were excited by the 640 nm laser. The top panel displays the uncorrected trajectory and the bottom the leak-corrected trajectory (See *Leak Factor Determination* in the SI). The black dashed line indicates the photobleaching of A1. Note the unchanged intensity of A2 in the lower panel, indicating negligible energy transfer between the acceptors. The difference in intensity between A1 and A2 originates from the difference in extinction coefficients (ε_640 nm_ = 64000 and 210000 M^-1^cm^-1^) and quantum yields (= 0.47 and 0.32). (**b**) Histogram of the relative change in fluorescence intensity of A2 after photobleaching of A1 (*ΔF*_A2_ = (<*N*_A2_>_before_ – <*N*_A2_>_after_)/<*N*_A2_>_after_, where the subscripts “before” and “after” denote the average intensities before and after photobleaching) for 41 double-labeled ClpB molecules. The red line indicates the mean value of 0.01, demonstrating no inter-acceptor energy transfer.

**Figure S19.**
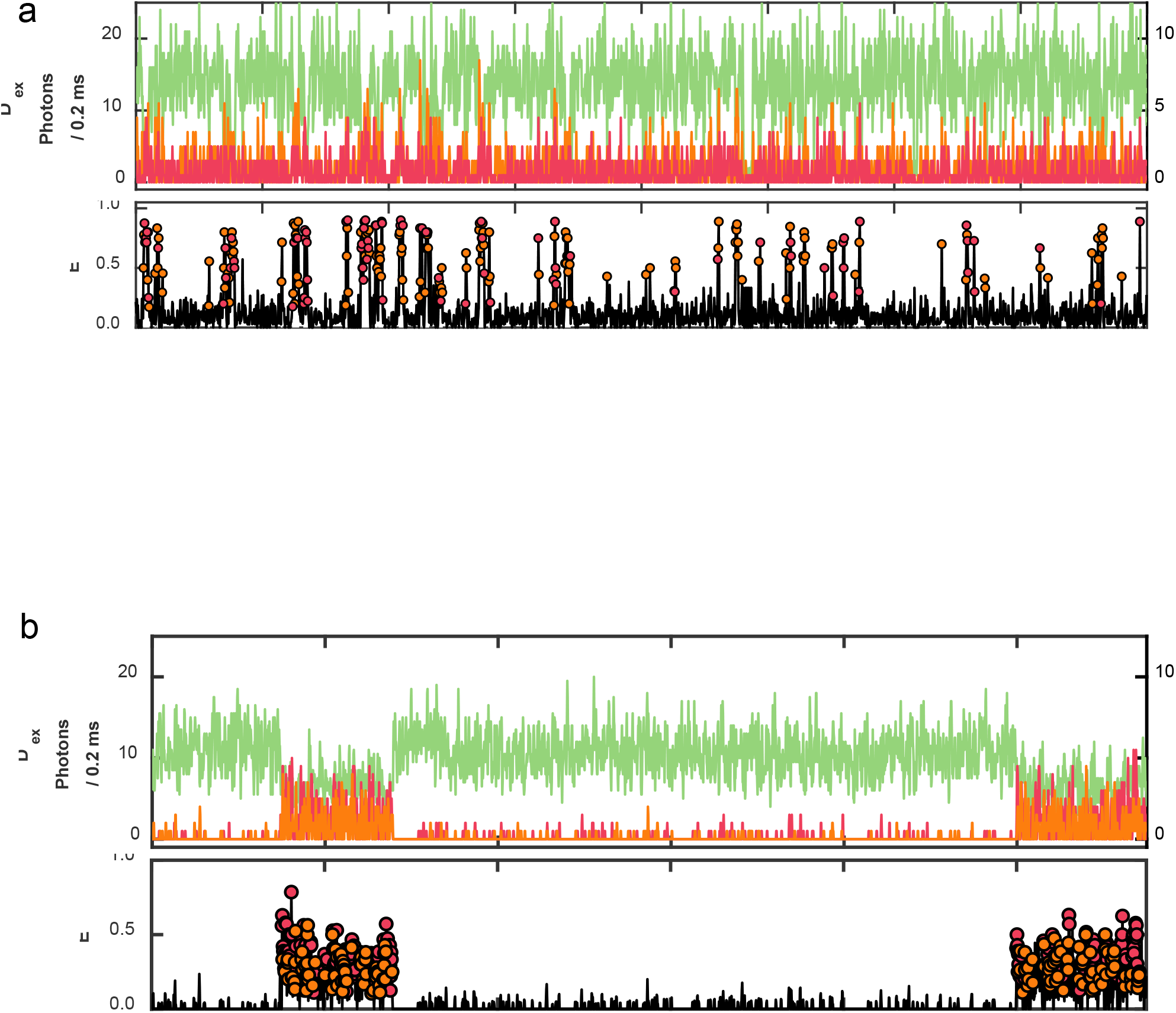
Sample trajectories of Cy3B-labeled casein translocating through a double-labeled ClpB. Trajectories are shown in the presence of (**a**) 2 mM ATP and (**b**) 2 mM ATPγS. The fluorescence of the donor (green, left axis) and acceptors (A1 – orange and A2 – red, right axis) after donor excitation (D_ex_) is shown in the top panel. *E*_app_ is shown in the middle panel, the colored circles indicate which acceptor is most dominant in each bin during the events. The constant leak-corrected acceptor emission (see Fig. S18) after acceptor excitation at 640 nm (bottom panels) demonstrates that both acceptors were active throughout the trajectory.

**Figure S20:**
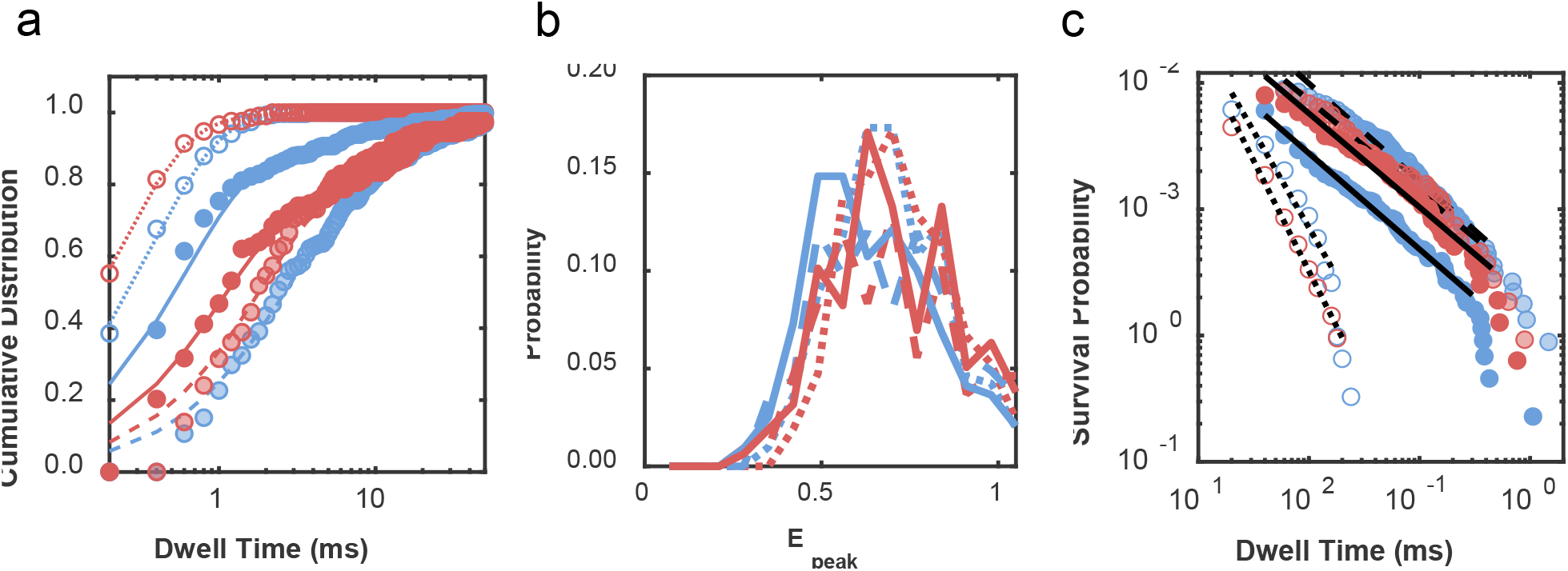
Characterization of the different types of interactions in the presence of ATP. Red solid-filled symbols and solid line – type I, blue solid-filled symbols and solid line – type II, red shaded symbols and dashed line – type III, blue shaded symbols and dashed line – type IV, red hollow symbols and dotted line – type V, and blue hollow symbols and dotted line – type VI. (**a**) Cumulative distribution of dwell times, (**b**) histogram of peak FRET efficiencies, and (**c**) power-law distributions. Parameters retrieved from the fits of (**a**) and (**c**) are provided in Table S6.

**Figure S21.**
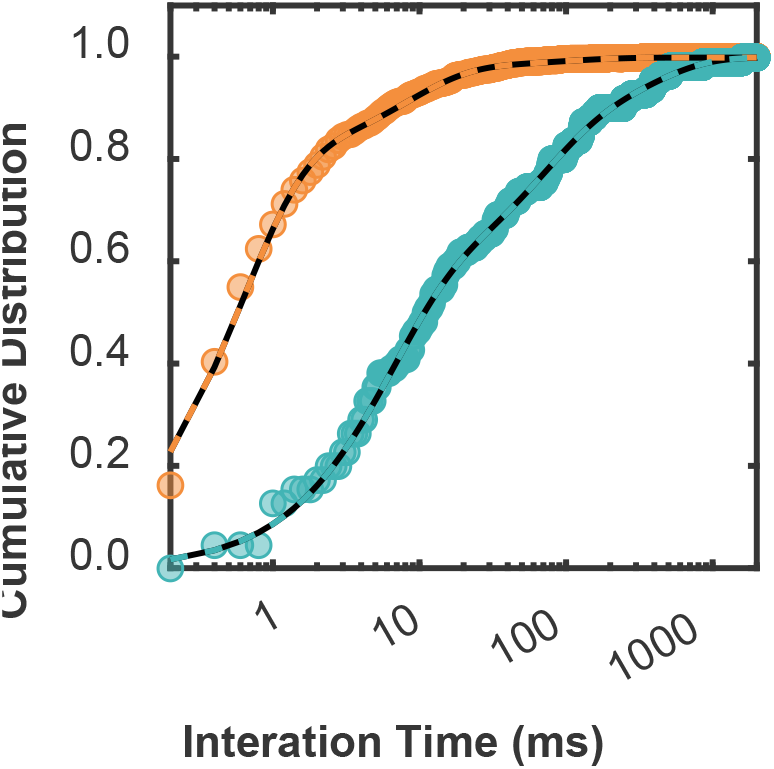
Cumulative dwell time histogram of all combined events from the three-color experiments. Cumulative distribution of dwell times in the presence of ATP (orange) and ATPγS (teal). The dashed lines represent the exponential fits corresponding to overall average dwell times of 3.8 and 76 ms with ATP or ATPγS, respectively. Fit parameters are provided in Table S7.

**Figure S22.**
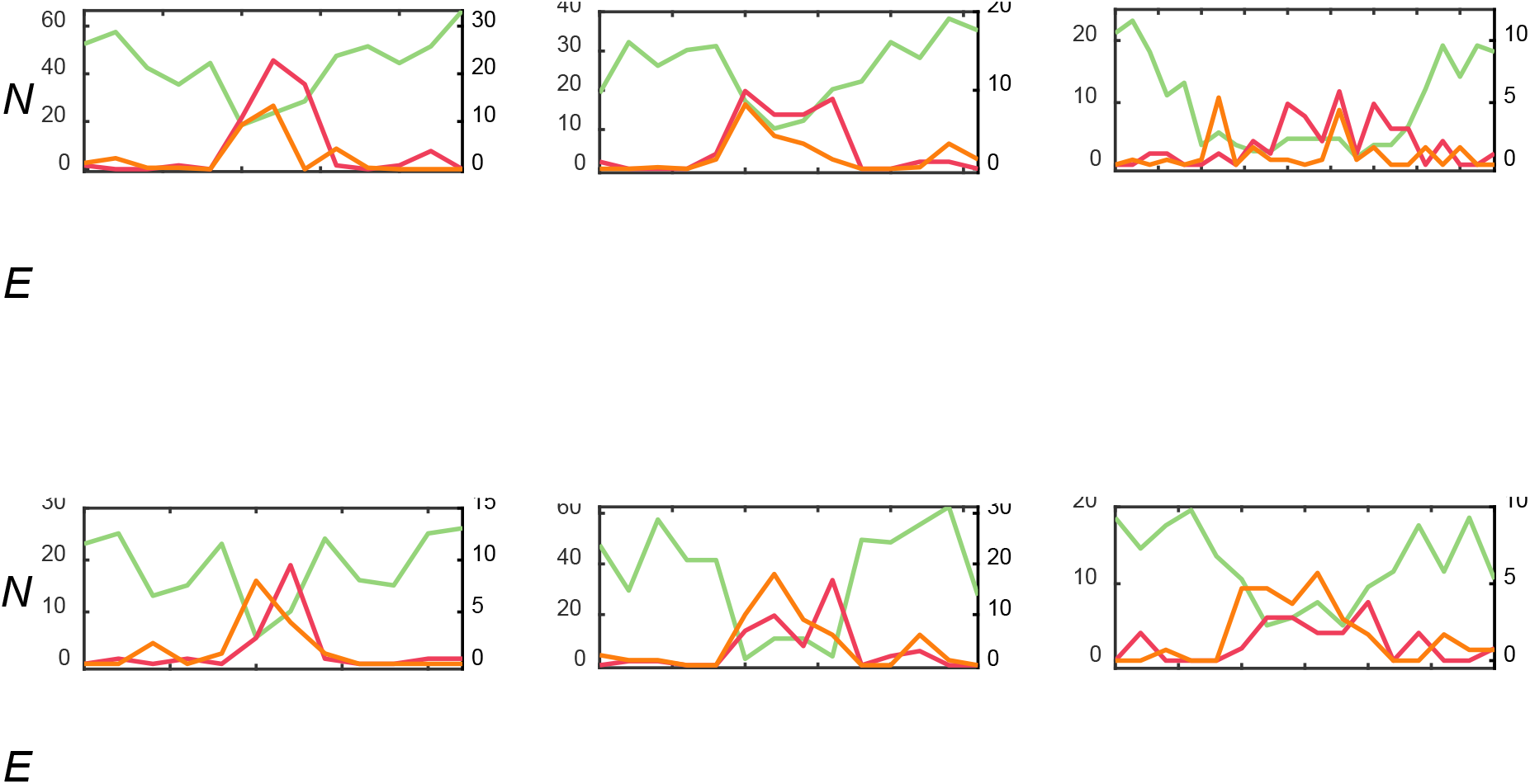
Samples of complete forward translocation type I events. The fluorescence of the donor (green, left axis), A1 (orange, right axis), and A2 (red, right axis) are shown in the upper panels with a bin size of 0.2 ms. The lower panels contain the apparent FRET efficiency for A1 (orange), A2 (red), and total (black). The colored circles indicate which acceptor is most dominant in each bin.

**Figure S23.**
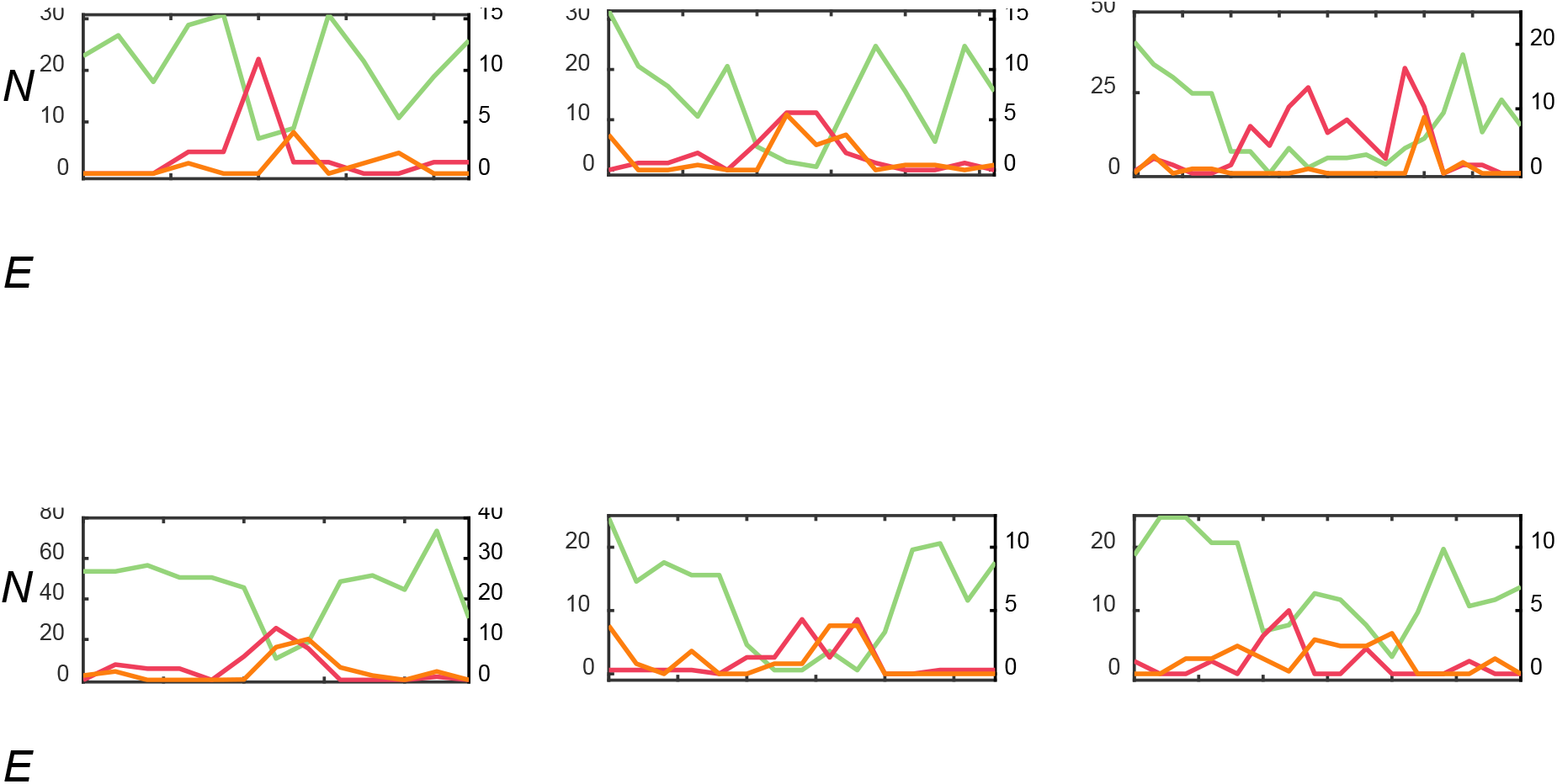
Samples of complete reverse translocation type II events. The fluorescence of the donor (green, left axis), A1 (orange, right axis), and A2 (red, right axis) are shown in the upper panels with a bin size of 0.2 ms. The lower panels contain the apparent FRET efficiency for A1 (orange), A2 (red), and total (black). The colored circles indicate which acceptor is most dominant in each bin.

**Figure S24.**
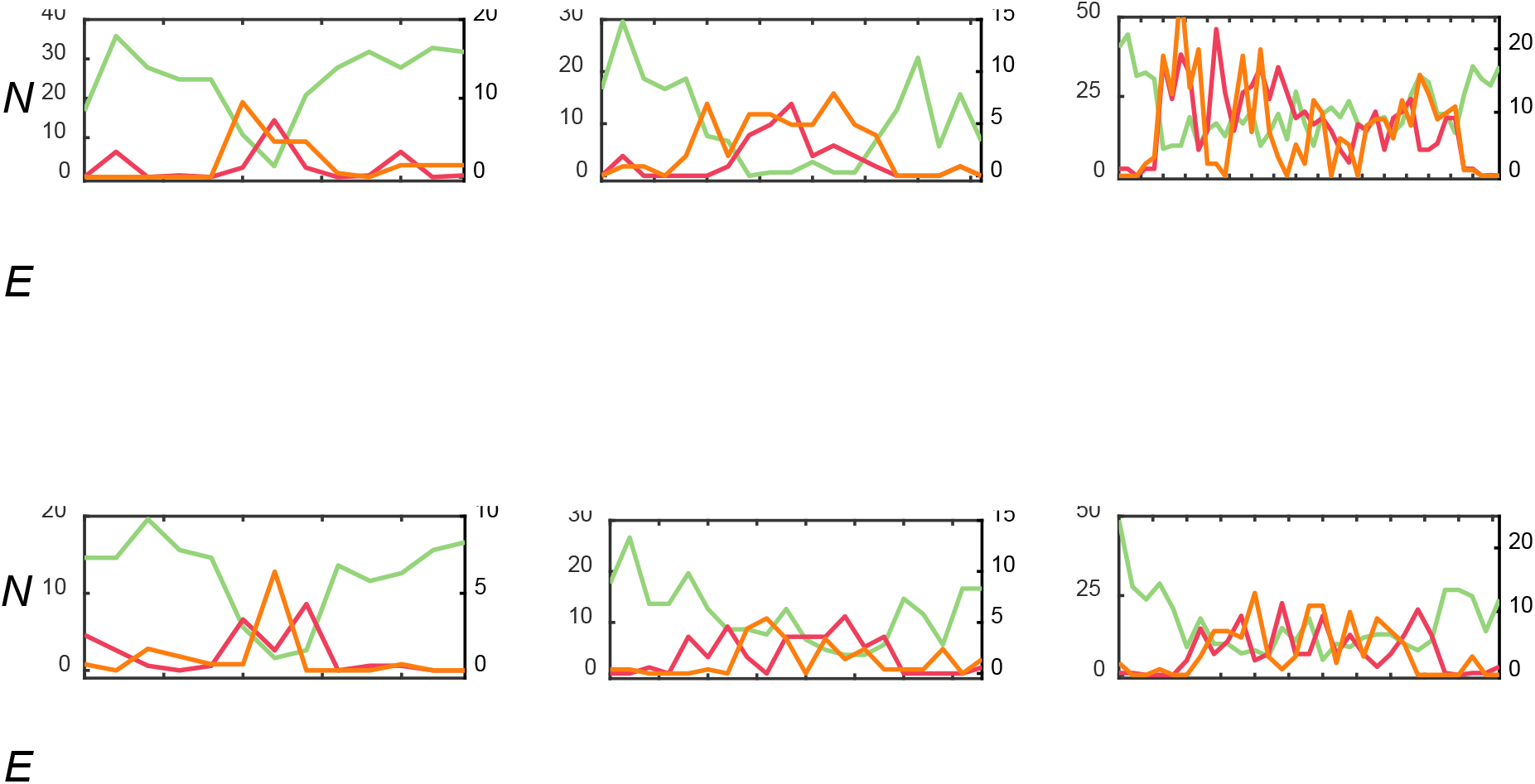
Samples of partial translocation types III (top) and IV (bottom) events. The fluorescence of the donor (green, left axis), A1 (orange, right axis), and A2 (red, right axis) are shown in the upper panels with a bin size of 0.2 ms. The lower panels contain the apparent FRET efficiency for A1 (orange), A2 (red), and total (black). The colored circles indicate which acceptor is most dominant in each bin.

**Figure S25.**
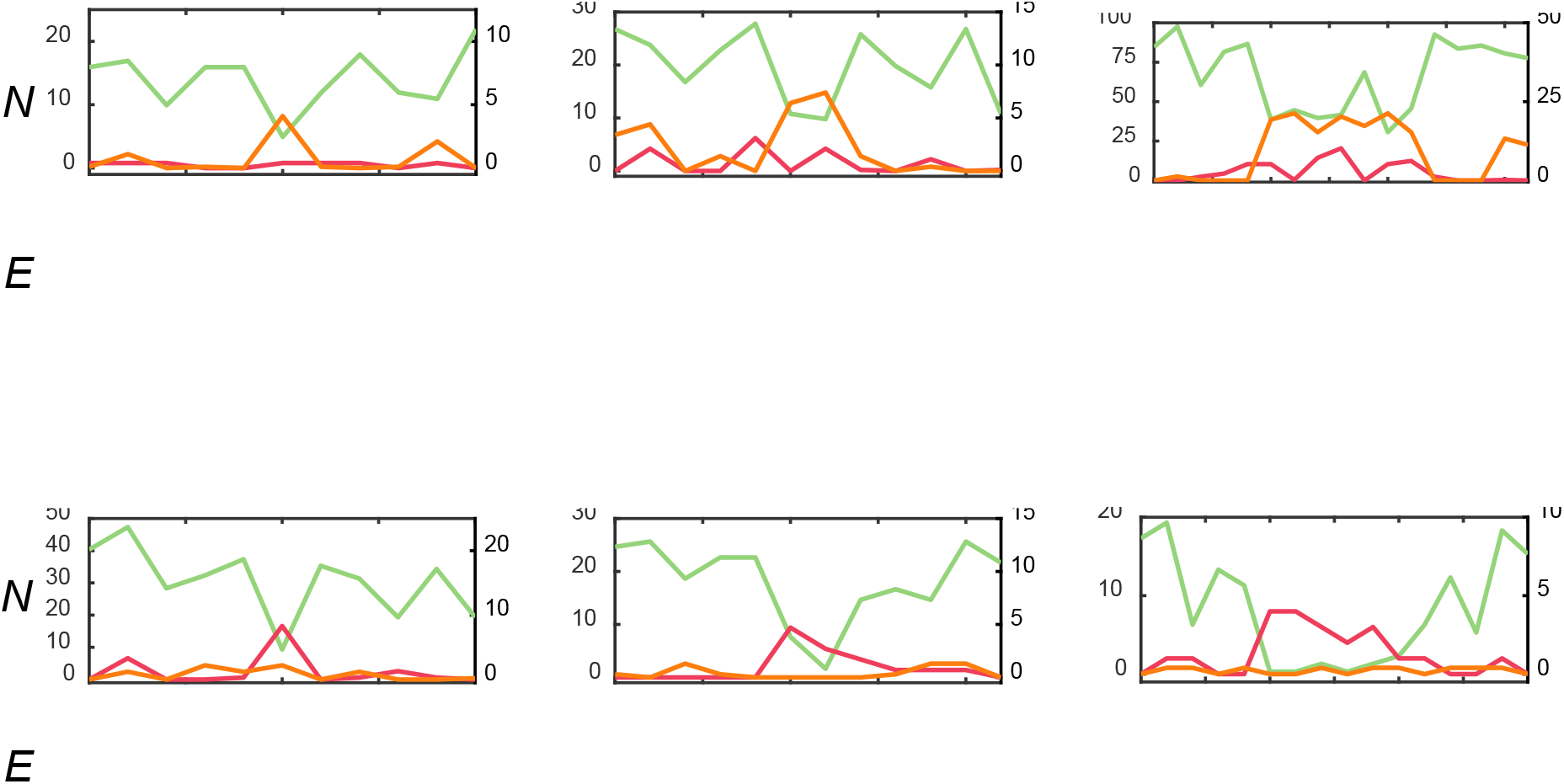
Samples of binding types V (top) and VI (bottom) events. The fluorescence of the donor (green, left axis), A1 (orange, right axis), and A2 (red, right axis) are shown in the upper panels with a bin size of 0.2 ms. The lower panels contain the apparent FRET efficiency for A1 (orange), A2 (red), and total (black). The colored circles indicate which acceptor is most dominant in each bin.

**Figure S26:**
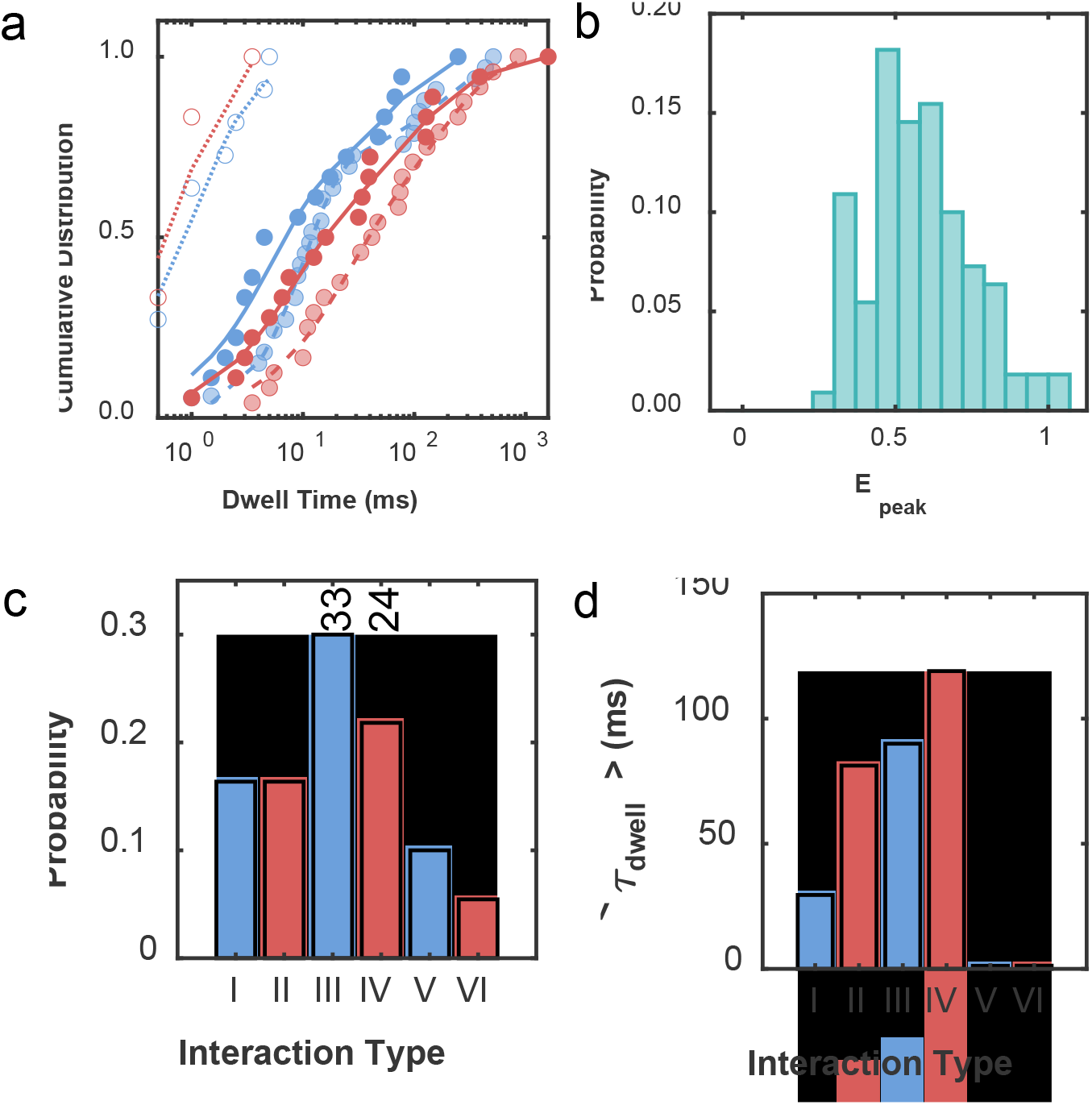
Characterization of the different types of events in the three-color experiments in the presence of ATPγS. (**a**) Cumulative distribution of dwell times for type I (red solid-filled symbols and solid line), II (blue solid-filled symbols and solid lines), III (red shaded symbols and dashed lines), IV (blue shaded symbols and dashed lines), V (red hollow symbols and dotted lines), and VI (blue hollow symbols and dotted lines). (**b**) Combined (due to the low number of interactions) peak FRET efficiencies. (**c**) Relative frequency (the numbers above the bars display the number of each type of event) and (**d**) average dwell time of each event type found from the fits in (**a**).

**Figure S27:**
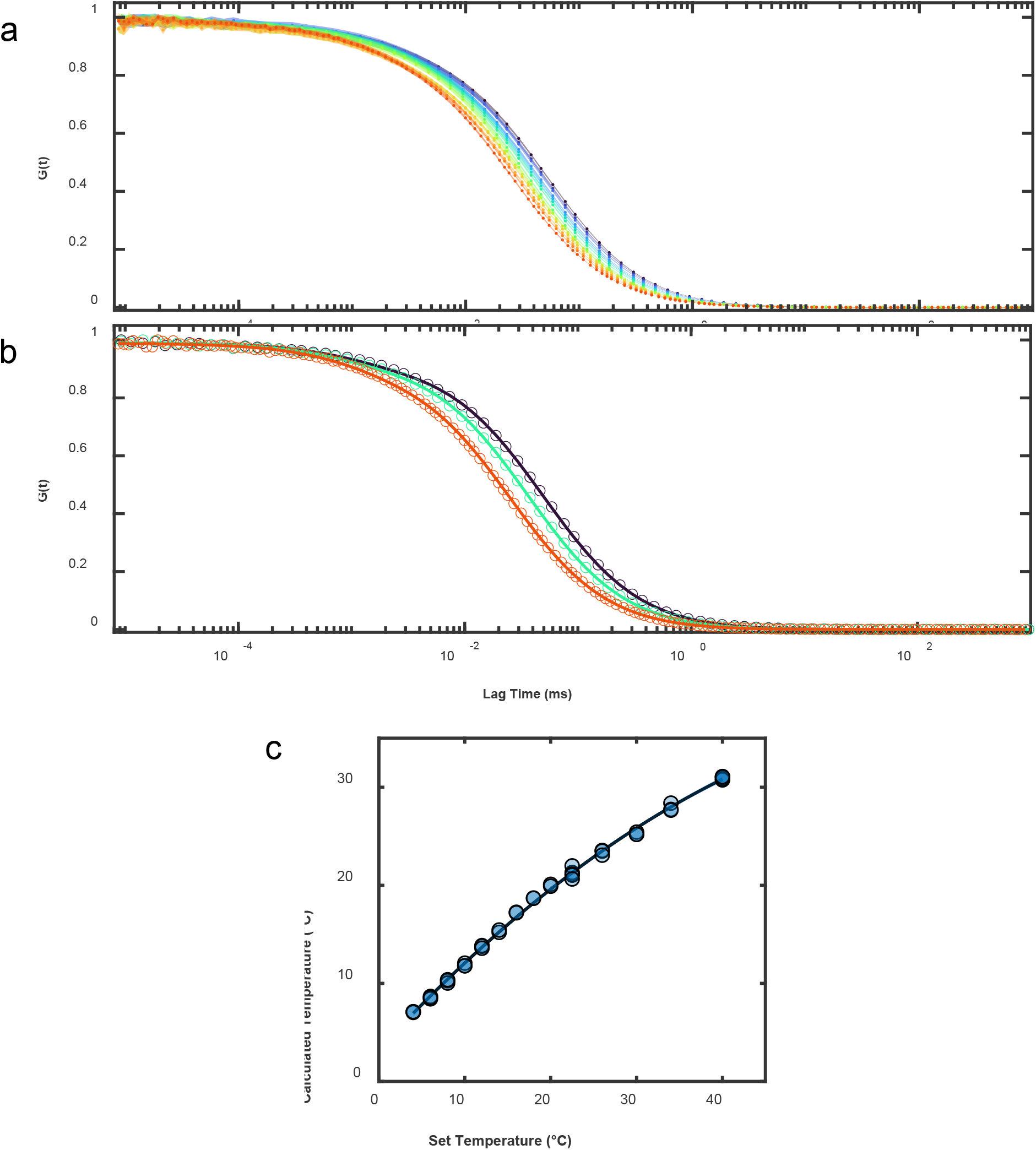
Calibration of the temperature cell. (**a**) Fluorescence correlation spectroscopy (FCS) curves of Cy3B in 25 mM HEPES, 25 mM KCl, 10 mM MgCl_2_ (pH 8). From (dark blue) top to (red) bottom, the temperature cell was set at 4, 6, 8, 10, 12, 14, 16, 20, 22.5, 26, 30, 34, 40 °C. The FCS curves were calculated using the PicoQuant SymPhoTime 64 software. The narrow shaded regions about the curves represent the uncertainty. (**b**) Sample fits (line) of the FCS curves (open circles) with Equation S16. To reduce clutter, only temperatures of 4, 16, and 40 °C are shown. (**c**) Calibration curve for the temperature retrieved from Equation S17 using the parameters retrieved from the fits of (**a**), corresponding to the temperature inside the confocal volume (Calculated Temperature), as a function of the temperature set on the Peltier controller. Each temperature measurement was repeated in triplicates. Parameters retrieved from the fits of FCS curves are provided in Table S12.

**Figure S28.**
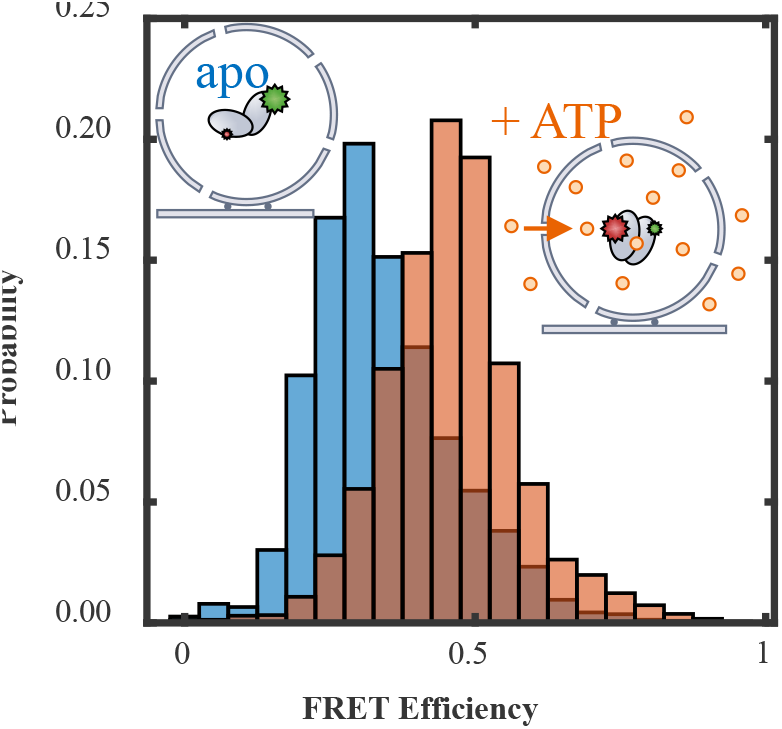
SOPC vesicles are permeable to ATP at 10 °C. Histograms of the FRET efficiency values of fluorescently labeled AK variant L82V (3) encapsulated in surface-immobilized 1-stearoyl-2-oleoyl-sn-glycero-3-phosphocholine (SOPC, 18:0-18:1 PC) liposomes at 10 °C without (blue) and with (orange) 2 mM ATP. AK was encapsulated in SOPC liposomes without nucleotides. After immobilization of the liposomes inside a flow cell, the temperature was turned to 10 °C for 15 minutes before data acquisition. Upon the addition of 2 mM ATP, the FRET efficiency histogram of AK, measured initially on 128 individual molecules, shifted to higher values (180 molecules), indicating a population shift that resulted from the SOPC membrane being permeable to ATP at 10 °C. Trajectories were monitored until photobleaching occurred using a circularly polarized 485 nm laser (ca. 200 nW).

**Figure S29.**
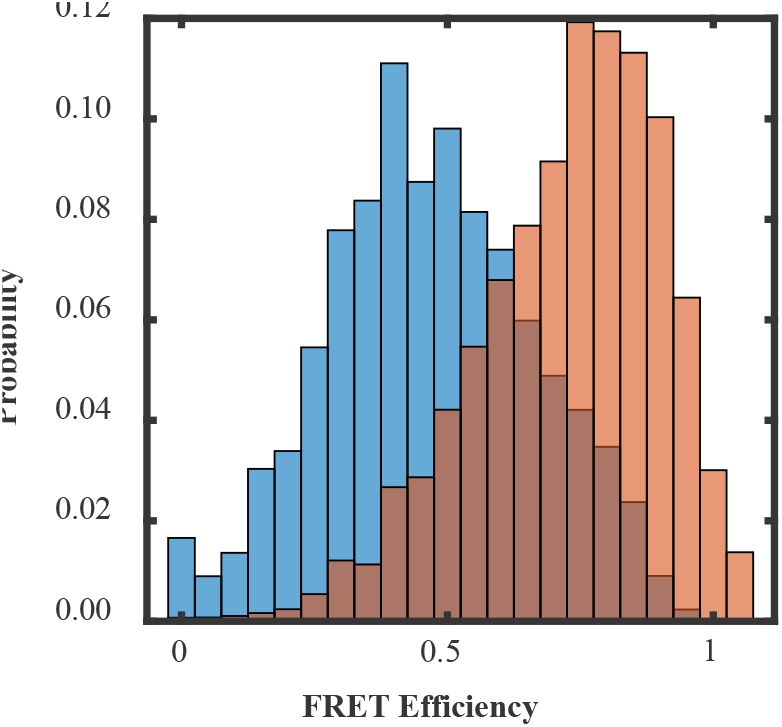
15:0 PC vesicles are permeable to ATP at 32 °C. Histograms of the FRET efficiencies fluorescently labeled AK L82V (3) encapsulated in surface-immobilized (1,2-dipentadecanoyl-sn-glycero-3-phosphocholine) 15:0 PC liposomes at 32 °C without (blue) and with (orange) 2 mM ATP. AK was encapsulated in 15:0 PC liposomes in the absence of ATP. The temperature was set to 32 °C for 20 minutes before data acquisition. Upon the addition of 2 mM ATP, the FRET efficiency histogram of AK, measured initially on 65 molecules, shifted to higher values (157 molecules), indicating that the 15:0 PC membrane was permeable to ATP at 32 °C. Trajectories were monitored until photobleaching occurred using a circularly polarized 485 laser (ca. 200 nW).

**Figure S30:**
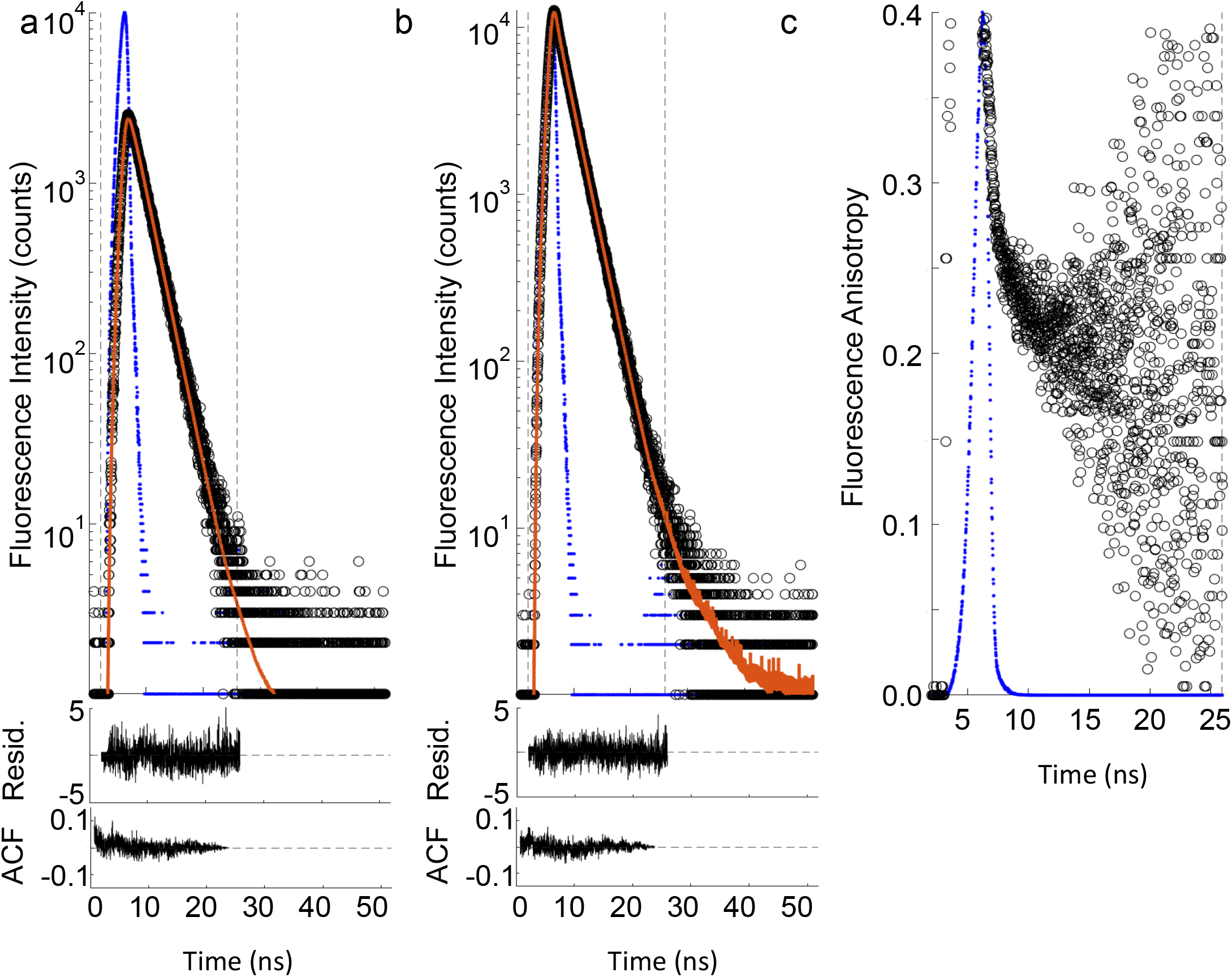
Global analysis of the time-resolved fluorescence anisotropy decay curves of ClpB 359. Fluorescence intensity (○) in the (**a**) *I*_VH_ and (**b**) *I*_VV_ channels and their corresponding fits to Equations S1 and S2 (orange lines) are given alongside (**c**) the fluorescence anisotropy decay. The dashed lines represent data limits used in the global analysis and the blue dots represent the instrument response function. The normalized residuals (Resid.) and autocorrelation functions (ACF) of the residuals are centered about zero, indicating a good fit. *χ*^2^ = 1.07.

**Figure S31.**
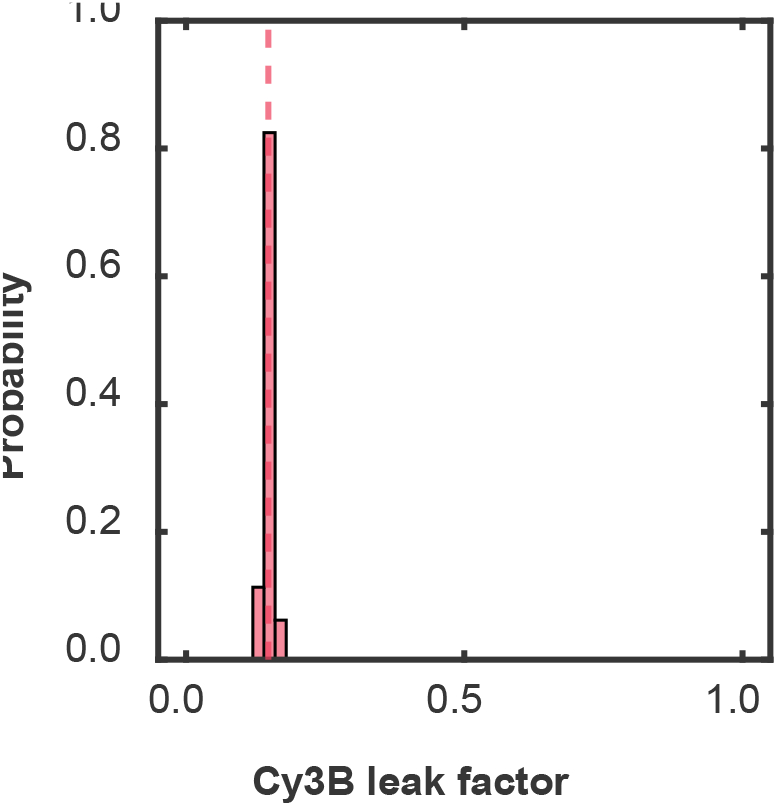
The per-molecule Cy3B average leak factor into the acceptor channel in two-color FRET experiments. The histogram is constructed from single-molecule data of 103 Cy3B labeled ClpB molecules, with an average value (dashed line) of 0.148.

**Figure S32.**
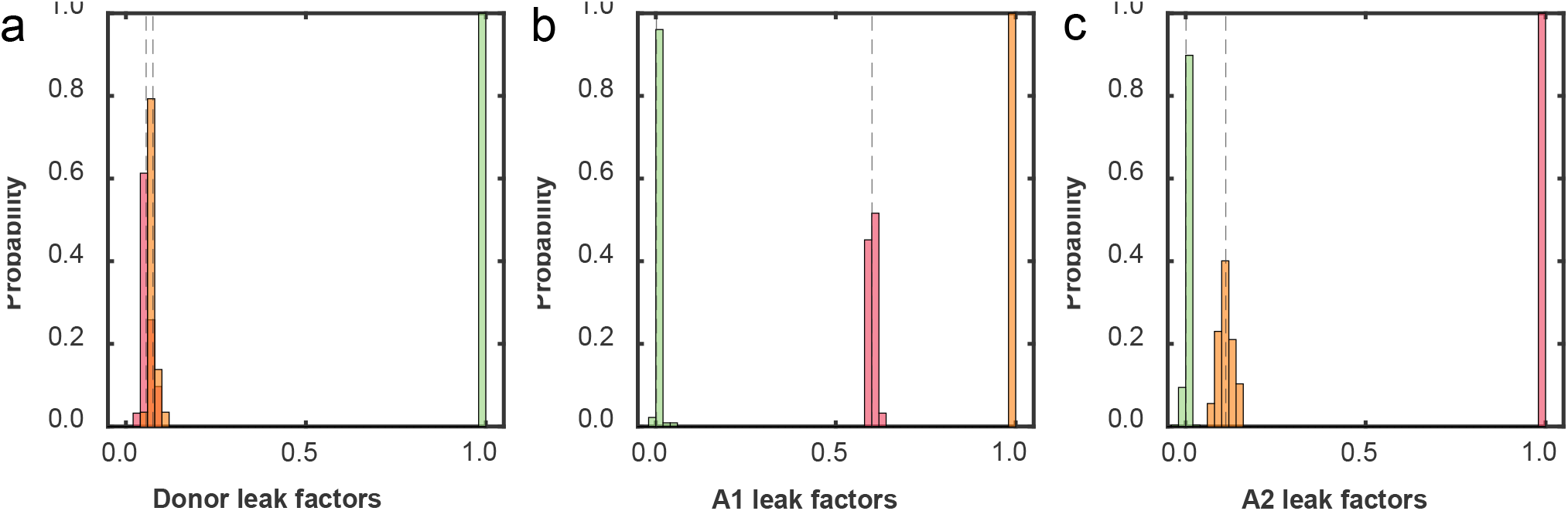
The per-molecule average leak factors in the three-color FRET experiments. Single-labeled (**a**) Cy3B (*N* = 154) casein, (**b**) AF647 ClpB (N = 240), and (**c**) CF680R ClpB (*N* = 289) molecules used in the three-color single-molecule experiments. The leak factors were taken as the average values of the distributions. The bar colors indicate the leak of the given species into the donor (Cy3B, green), acceptor 1 (AF647, orange), and acceptor 2 (CF680R, red) channels. The dashed lines represent the average values.

## Supporting Tables

**Table S1.** Summary of the number of labeled ClpB molecules sampled under various experimental conditions.

| Dye position | Nucleotide | Nucleotide Concentration (mM) | Temperature (°C) | Number of ClpB molecules | Number of events |
| --- | --- | --- | --- | --- | --- |
| NBD1 (359) | ATP | 2.0 | 10 | 84 | 1202 |
|  |  |  | 22.5 | 166 | 1537 |
|  |  |  | 32 | 74 | 532 |
|  |  | 0.4 | 22.5 | 164 | 744 |
|  |  | 0.2 |  | 173 | 1113 |
| | ATPγS | 2.0 | | 147 | 337 |
| NBD2 (771) | ATP | 2.0 | 10 | 67 | 696 |
|  |  |  | 22.5 | 137 | 1214 |
|  |  |  | 32 | 69 | 621 |
|  |  | 0.4 | 22.5 | 153 | 890 |
|  |  | 0.2 |  | 112 | 950 |
| | ATPγS | 2.0 | | 113 | 227 |
| NBD1 (176) + NBD2 (771) | ATP | 2.0 | 22.5 | 135 | 1445 |
| | ATPγS | 2.0 | | 110 | 135 |

**Table S2.** Dwell time characterization as a function of temperature and ATP concentration.

| Dye position | [ATP] (mM) | Temperature (°C) | Power Law Exponent<sup>a</sup> | Fraction of short events<sup>b</sup> | $\langle\tau_{\text{dwell}}\rangle$ (ms)<sup>c</sup> |
| --- | --- | --- | --- | --- | --- |
| NBD1 (359) | 2.0 | 10 | $1.70 \pm 0.01$ | 0.88 | $1.70 \pm 0.18$ |
| | | 22.5 | $1.69 \pm 0.02$ | 0.90 | $1.60 \pm 0.13$ |
| | | 32 | $1.92 \pm 0.01$ | 0.93 | $1.43 \pm 0.10$ |
| | 0.4 | 22.5 | $1.46 \pm 0.02$ | 0.83 | $1.87 \pm 0.07$ |
| | 0.2 | 22.5 | $1.40 \pm 0.01$ | 0.88 | $1.59 \pm 0.08$ |
| NBD2 (771) | 2.0 | 10 | $1.83 \pm 0.03$ | 0.89 | $1.55 \pm 0.08$ |
| | | 22.5 | $1.55 \pm 0.02$ | 0.86 | $1.58 \pm 0.14$ |
| | | 32 | $1.49 \pm 0.01$ | 0.88 | $1.37 \pm 0.09$ |
| | 0.4 | 22.5 | $1.42 \pm 0.02$ | 0.91 | $1.94 \pm 0.07$ |
| | 0.2 | 22.5 | $1.44 \pm 0.02$ | 0.88 | $1.54 \pm 0.09$ |
| NBD1 (176) + NBD2 (771) | 2.0 | 22.5 | $1.68 \pm 0.01^{\text{d}}$ | 0.91 | $1.32 \pm 0.21^{\text{d}}$ |

<sup>a</sup> Exponents retrieved from the dwell time survival functions.
<sup>b</sup> Fraction of the events which were $\leq 10$ ms in length.
<sup>c</sup> Lifetimes retrieved from the exponential fits of the short events ( $\leq 10$ ms).
<sup>d</sup> Overall values. A breakdown of the values for each event type is provided in Table S7.

**Table S3.** Parameters retrieved from the triexponential fits of the combined (NBD1 and NBD2 labeled ClpB) two-color dwell time CDF.

| Nucleotide | $a_1$ | $\tau_1$ (ms) | $a_2$ | $\tau_2$ (ms) | $a_3$ | $\tau_3$ (ms) | Weighted Average (ms) |
| --- | --- | --- | --- | --- | --- | --- | --- |
| ATP | 0.86 | 1.52 | 0.11 | 32.3 | 0.02 | 780 | 21.8 |
| ATPγS | 0.30 | 27.2 | 0.33 | 374 | 0.37 | 1670 | 749 |

Note: The $a_i$ and $\tau_i$ values are the preexponential factors and lifetimes for the $i^{\text{th}}$ exponent, respectively.

**Table S4.**
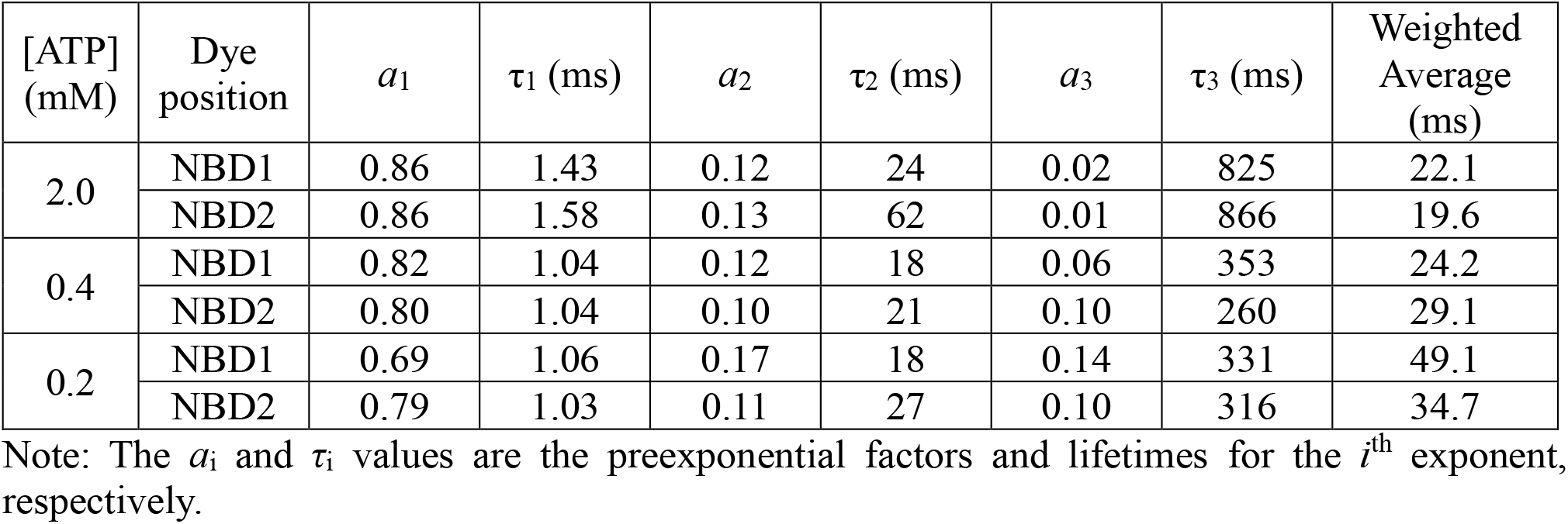
Parameters retrieved from the triexponential fits of the two-color dwell time CDF as a function of ATP concentration.

| [ATP] (mM) | Dye position | $a_1$ | $\tau_1$ (ms) | $a_2$ | $\tau_2$ (ms) | $a_3$ | $\tau_3$ (ms) | Weighted Average (ms) |
| --- | --- | --- | --- | --- | --- | --- | --- | --- |
| 2.0 | NBD1 | 0.86 | 1.43 | 0.12 | 24 | 0.02 | 825 | 22.1 |
|  | NBD2 | 0.86 | 1.58 | 0.13 | 62 | 0.01 | 866 | 19.6 |
| 0.4 | NBD1 | 0.82 | 1.04 | 0.12 | 18 | 0.06 | 353 | 24.2 |
|  | NBD2 | 0.80 | 1.04 | 0.10 | 21 | 0.10 | 260 | 29.1 |
| 0.2 | NBD1 | 0.69 | 1.06 | 0.17 | 18 | 0.14 | 331 | 49.1 |
|  | NBD2 | 0.79 | 1.03 | 0.11 | 27 | 0.10 | 316 | 34.7 |

Note: The $a_i$ and $\tau_i$ values are the preexponential factors and lifetimes for the $i^{\text{th}}$ exponent, respectively.

**Table S5.** Parameters retrieved from the exponential fits of the frequency of events as a function of ATP concentration.

| Dye position | [ATP] (mM) | ATPase Activity<sup>a</sup><br>(ATP s<sup>-1</sup> hexamer<sup>-1</sup>) | Event Frequency<sup>b</sup> (s<sup>-1</sup>) |
| --- | --- | --- | --- |
| NBD1 (359) | 2.0 | 1.14 ± 0.03 | 2.09 ± 0.09 |
|  | 0.4 | 0.76 ± 0.05 | 1.03 ± 0.06 |
|  | 0.2 | 0.39 ± 0.03 | 0.48 ± 0.03 |
| NBD2 (771) | 2.0 | 1.14 ± 0.03 | 1.63 ± 0.12 |
|  | 0.4 | 0.76 ± 0.05 | 0.74 ± 0.02 |
|  | 0.2 | 0.39 ± 0.03 | 0.49 ± 0.01 |

<sup>a</sup> With the addition of 3 μM κ-casein.
<sup>b</sup> Average number of events per unit time.

**Table S6.** Summary of parameters characterizing the different types of events.

| Event Type | 2 mM ATP | | | | 2 mM ATPγS | | |
| --- | --- | --- | --- | --- | --- | --- | --- |
| | Fraction | $N^a$ | $\tau_{\text{dwell}}$ (ms)<sup>b</sup> | Power Law Exponent<sup>c</sup> | Fraction | $N^a$ | $\tau_{\text{dwell}}$ (ms)<sup>b</sup> |
| I | 0.30 | 438 | $2.1 \pm 0.2$ | 1.78 | 0.17 | 18 | $28.0 \pm 2.3$ |
| II | 0.11 | 158 | $6.6 \pm 1.9$ | 1.73 | 0.17 | 18 | $81.2 \pm 16.3$ |
| III | 0.16 | 225 | $11.3 \pm 2.1$ | 1.76 | 0.31 | 33 | $90.0 \pm 4.4$ |
| IV | 0.07 | 108 | $6.2 \pm 0.5$ | 1.74 | 0.23 | 24 | $119.0 \pm 17.6$ |
| V | 0.21 | 306 | $0.4 \pm 0.1$ | 2.53 | 0.08 | 11 | $1.2 \pm 0.1$ |
| VI | 0.15 | 210 | $0.3 \pm 0.1$ | 2.75 | 0.04 | 6 | $1.3 \pm 0.1$ |

<sup>a</sup> The number of event types observed. Note the limited sampling in the presence of ATPγS due to the long dwell times.
<sup>b</sup> Calculated from the exponential fit of the CDF for the different events.
<sup>c</sup> Exponents retrieved from the dwell time survival functions.

**Table S7.** Parameters retrieved from the triexponential fit of the three-color dwell time CDF in the presence of ATP and ATPγS.

| Nucleotide | $a_1$ | $\tau_1$ (ms) | $a_2$ | $\tau_2$ (ms) | $a_3$ | $\tau_3$ (ms) | Weighted Average (ms) |
| --- | --- | --- | --- | --- | --- | --- | --- |
| ATP | 0.79 | 0.6 | 0.19 | 8.0 | 0.02 | 88 | 3.8 |
| ATPγS | 0.53 | 6.0 | 0.30 | 60.5 | 0.16 | 332 | 75.9 |

Note: The $a_i$ and $\tau_i$ values are the preexponential factors and lifetimes for the $i^{\text{th}}$ exponent, respectively.

**Table S8.**
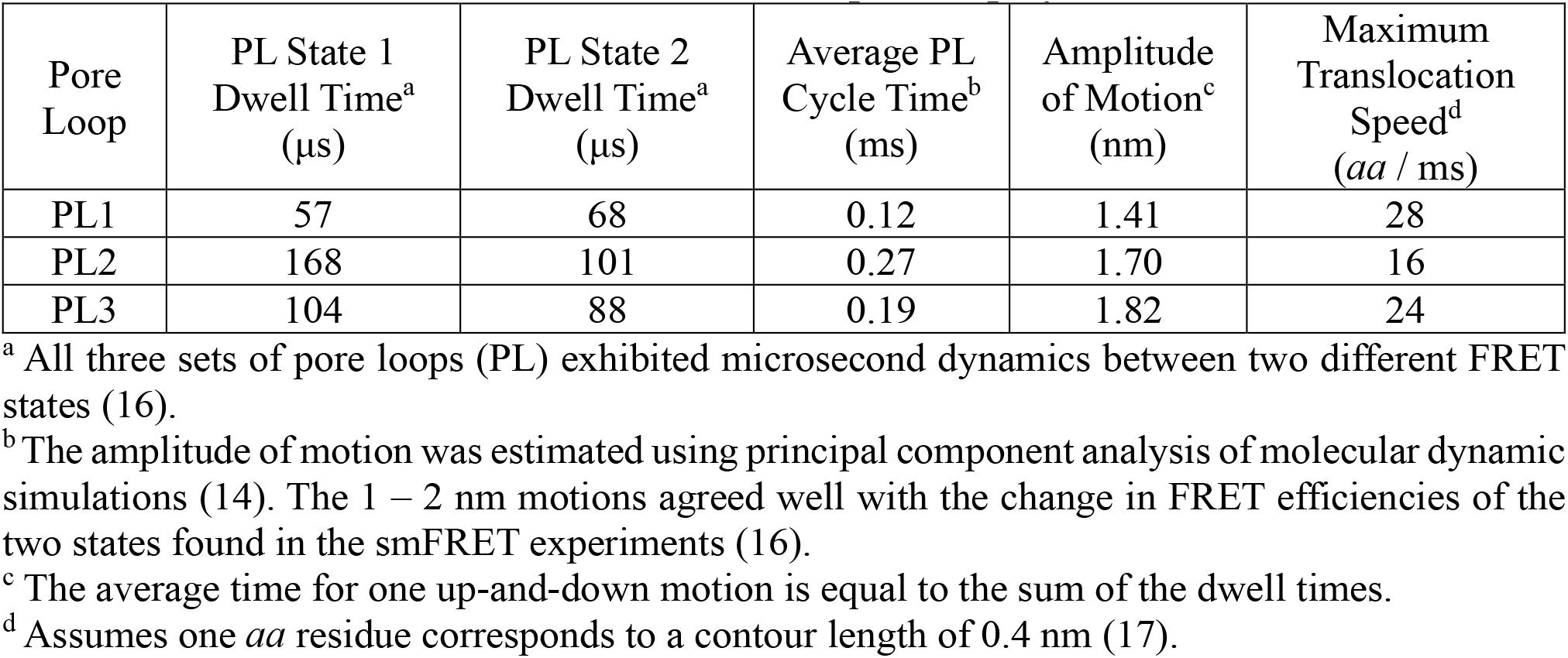
Translocation velocities estimated from pore loop dynamics.

**Table S9.** Size distribution of extruded liposomes.

| Repeat<sup>a</sup> | z-Average Diameter (nm) | Particle Dispersity Index |
| --- | --- | --- |
| 1 | 119 | 0.04 |
| 2 | 119 | 0.05 |
| 3 | 118 | 0.07 |

<sup>a</sup> Repeats were taken from independently prepared solutions of liposomes.

**Table S10.** Summary of the steady-state fluorescence anisotropy of fluorescently labeled ClpB.

| Species | Steady-State Anisotropy | $r_{SS,calc}^a$ |
| --- | --- | --- |
| Free dye | $0.097 \pm 0.01$ | - |
| ClpB 359CyB | $0.271 \pm 0.01$ | 0.26 |
| ClpB 771Cy3B | $0.277 \pm 0.01$ | 0.30 |
| ClpB 771CF680R | $0.227 \pm 0.02$ | - |

<sup>a</sup> Calculated steady-state anisotropy value (Equation S5) using parameters retrieved from the time-resolved fluorescence analysis (Table S11).

**Table S11.** Parameters retrieved from the global analysis of the fluorescence anisotropy decays of Cy3B-labeled ClpB.

| Parameter | ClpB 359Cy3B | ClpB 771Cy3B |
| --- | --- | --- |
| $\chi^2$ | 1.07 | 1.11 |
| $\tau_D$ (ns) | 2.7 | 2.8 |
| $G$ | 0.51 | |
| $\phi_F$ (ns)<sup>†</sup> | 0.95 | 1.11 |
| contribution ($\alpha$) of the fast tumbling time | 0.45 | 0.31 |
| $\phi_P$ (ns)<sup>††</sup> | 210 | |
| $r_{SS,calc}^a$ | 0.26 | 0.30 |

<sup>†</sup> Fast tumbling time associated with rotational freedom of the dye.
<sup>††</sup> The slow tumbling time of ClpB occurs on a time scale much slower than the decay of Cy3B. A tumbling time of 210 ns, predicted by the Perrin equation for a 12 nm diameter sphere, was used as a fixed value in the analysis (18).
<sup>a</sup> Calculated steady-state fluorescence anisotropy values (Equation S5).

**Table S12:** Parameters retrieved from the FCS curves of Cy3B at different temperatures. A description of the model (Equations S15 and S16) and parameters is provided above in the SI text under *Fluorescence correlation spectroscopy*).

| Set T (°C)<sup>a</sup> | $\omega$ | $f_T$ | $\tau_T$ (μs) | $N$ | $\tau_D$ (μs) | $G_\infty \times 10^4$ | Cell T (°C)<sup>b</sup> |
| --- | --- | --- | --- | --- | --- | --- | --- |
| 4 | 6.3 | $0.064 \pm 0.001$ | $1.0 \pm 0.1$ | $1.07 \pm 0.01$ | $49.8 \pm 0.1$ | $-2 \pm 2$ | $7.08 \pm 0.04$ |
| 6 | | $0.061 \pm 0.001$ | $1.0 \pm 0.2$ | $1.07 \pm 0.05$ | $47.4 \pm 0.2$ | $-1 \pm 2$ | $8.6 \pm 0.1$ |
| 8 | | $0.061 \pm 0.003$ | $1.0 \pm 0.1$ | $1.07 \pm 0.04$ | $44.9 \pm 0.3$ | $-3 \pm 1$ | $10.2 \pm 0.2$ |
| 10 | | $0.062 \pm 0.002$ | $1.0 \pm 0.2$ | $1.07 \pm 0.05$ | $42.7 \pm 0.2$ | $-1 \pm 2$ | $11.9 \pm 0.2$ |
| 12 | | $0.060 \pm 0.001$ | $1.1 \pm 0.2$ | $1.07 \pm 0.02$ | $40.3 \pm 0.2$ | $-2 \pm 2$ | $13.7 \pm 0.1$ |
| 14 | | $0.060 \pm 0.001$ | $1.0 \pm 0.1$ | $1.07 \pm 0.03$ | $38.5 \pm 0.2$ | $-2 \pm 1$ | $15.3 \pm 0.2$ |
| 16 | | $0.059 \pm 0.001$ | $0.8 \pm 0.1$ | $1.07 \pm 0.03$ | $36.4 \pm 0.1$ | $-1 \pm 1$ | $17.22 \pm 0.04$ |
| 18 | | $0.058 \pm 0.001$ | $0.9 \pm 0.1$ | $1.06 \pm 0.01$ | $34.9 \pm 0.1$ | $-2 \pm 1$ | $18.68 \pm 0.02$ |
| 20 | | $0.054 \pm 0.002$ | $0.8 \pm 0.1$ | $1.07 \pm 0.02$ | $33.6 \pm 0.1$ | $0 \pm 1$ | $20.0 \pm 0.1$ |
| 22.5 | | $0.071 \pm 0.014$ | $0.8 \pm 0.2$ | $1.09 \pm 0.21$ | $32.2 \pm 0.3$ | $-1 \pm 2$ | $21.2 \pm 0.5$ |
| 26 | | $0.081 \pm 0.001$ | $0.8 \pm 0.1$ | $1.10 \pm 0.04$ | $30.1 \pm 0.2$ | $0 \pm 1$ | $23.4 \pm 0.3$ |
| 30 | | $0.078 \pm 0.002$ | $0.8 \pm 0.1$ | $1.10 \pm 0.04$ | $28.6 \pm 0.1$ | $1 \pm 2$ | $25.3 \pm 0.1$ |
| 34 | | $0.076 \pm 0.001$ | $1.0 \pm 0.2$ | $1.09 \pm 0.03$ | $26.7 \pm 0.3$ | $1 \pm 1$ | $27.9 \pm 0.4$ |
| 40 | | $0.077 \pm 0.001$ | $1.1 \pm 0.2$ | $1.10 \pm 0.05$ | $24.7 \pm 0.1$ | $2 \pm 1$ | $32.0 \pm 0.2$ |

<sup>a</sup> Temperature set on the Peltier control unit.
<sup>b</sup> Temperature calculated (Equation S17) inside the confocal volume.

**Table S13.**
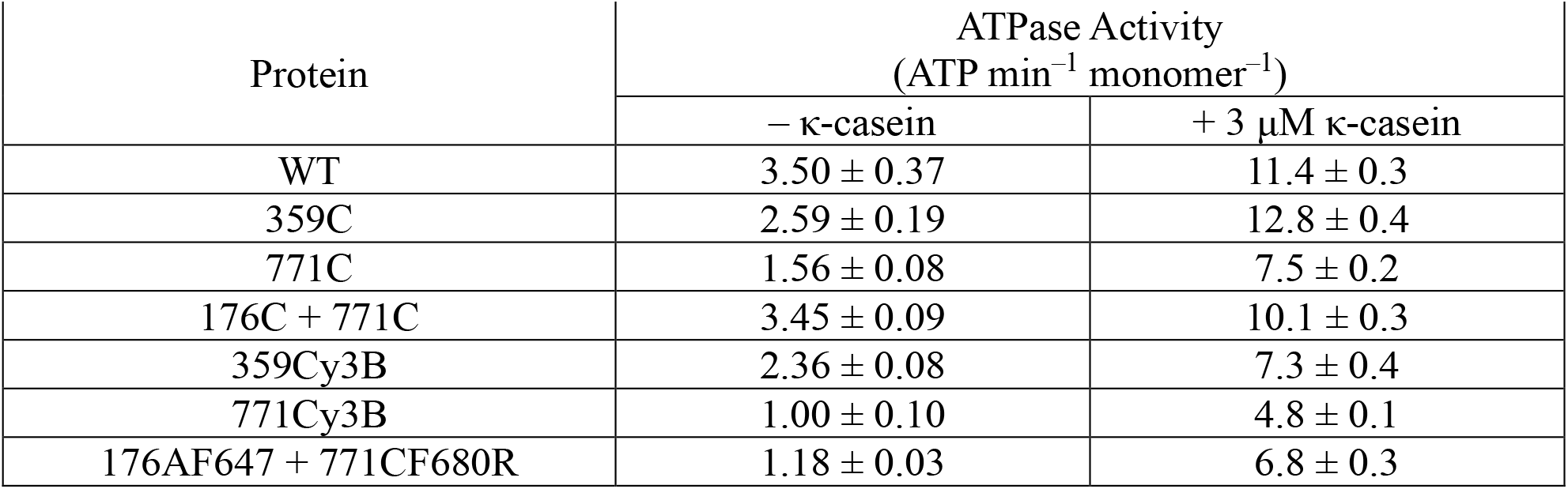
ATPase activity of ClpB in the presence of 2 mM ATP.

| Protein | ATPase Activity<br>(ATP min<sup>-1</sup> monomer<sup>-1</sup>) |  |
| --- | --- | --- |
| | – κ-casein | + 3 μM κ-casein |
| WT | $3.50 \pm 0.37$ | $11.4 \pm 0.3$ |
| 359C | $2.59 \pm 0.19$ | $12.8 \pm 0.4$ |
| 771C | $1.56 \pm 0.08$ | $7.5 \pm 0.2$ |
| 176C + 771C | $3.45 \pm 0.09$ | $10.1 \pm 0.3$ |
| 359Cy3B | $2.36 \pm 0.08$ | $7.3 \pm 0.4$ |
| 771Cy3B | $1.00 \pm 0.10$ | $4.8 \pm 0.1$ |
| 176AF647 + 771CF680R | $1.18 \pm 0.03$ | $6.8 \pm 0.3$ |

**Table S14.** Parameters retrieved from the fit of the ATPase activity by the Hill equation.

| Conditions | $K_m$ (μM) | $n$ |
| --- | --- | --- |
| – κ-casein | $0.42 \pm 0.01$ | $2.74 \pm 0.11$ |
| + 3 μM κ-casein | $0.31 \pm 0.02$ | $1.87 \pm 0.17$ |

